# ClusTrast: a short read *de novo* transcript isoform assembler guided by clustered contigs

**DOI:** 10.1101/2022.01.02.473666

**Authors:** Karl Johan Westrin, Warren W. Kretzschmar, Olof Emanuelsson

**Affiliations:** Science for Life Laboratory, Department of Gene Technology, KTH Royal Institute of Technology, SE-171 65, Solna, Sweden; Center for Hematology and Regenerative Medicine (HERM), Department of Medicine Huddinge, Karolinska Institute, SE-141 52, Flemingsberg, Sweden

## Abstract

**Background:** Transcriptome assembly from RNA-sequencing data in species without a reliable reference genome has to be performed *de novo*, but studies have shown that *de novo* methods often have inadequate ability to reconstruct transcript isoforms. We address this issue by constructing an assembly pipeline whose main purpose is to produce a comprehensive set of transcript isoforms.

**Results:** We present the *de novo* transcript isoform assembler ClusTrast, which takes short read RNA-seq data as input, constructs a primary assembly, clusters a set of guiding contigs, aligns the short reads to the guiding contigs, assembles each clustered set of short reads individually, and merges the primary and clusterwise assemblies into the final assembly. We tested ClusTrast on real datasets from six eukaryotic species, and showed that ClusTrast reconstructed more expressed known isoforms than any of the other tested *de novo* assemblers, at a moderate reduction in precision. For recall, ClusTrast was on top in the lower end of expression levels (<15% percentile) for all tested datasets, and over the entire range for almost all datasets. Reference transcripts were often (35–69% for the six datasets) reconstructed to at least 95% of their length by ClusTrast, and more than half of reference transcripts (58–81%) were reconstructed with contigs that exhibited polymorphism, measuring on a subset of reliably predicted contigs. ClusTrast recall increased when using a union of assembled transcripts from more than one assembly tool as primary assembly.

**Conclusion:** We suggest that ClusTrast can be a useful tool for studying isoforms in species without a reliable reference genome, in particular when the goal is to produce a comprehensive transcriptome set with polymorphic variants.

## 1 Background

In eukaryotes, many genes can produce RNA transcripts of differing base sequences called transcript isoforms. Transcript isoforms are created by alternative transcriptional start sites, splicing, or polyadenylation. Controlling transcript isoform expression is one way for a cell to regulate protein expression and thereby its behavior (Wang et al., 2008; Barbosa-Morais et al., 2012; Floor and Doudna, 2016). Changes in transcript isoform expression have been associated with developmental changes and tissue specificity in eukaryotes, and disease in humans (Fackenthal and Godley, 2008; Sterne-Weiler and Sanford, 2014; Xiong et al., 2015; Akhter et al., 2018). Thus, it is often important to clarify not only what genes are expressed but also which transcript isoforms are expressed.

The expression of genes and transcripts is often studied by RNA-sequencing, where short reads (SRs) derived from massively parallel shotgun sequencing are aligned to an organism’s reference genome. With this approach, reconstructing transcripts is possible by using the reference genome as a guide (Garber et al., 2011). However, many non-model organisms do not have a high-quality reference genome available. In such cases, a commonly used approach is *de novo* assembly in which transcripts are assembled from the reads only. The assembled transcripts are sometimes referred to as contigs or reconstructed transcripts. Popular tools to perform *de novo* transcriptome assembly include Trans-ABySS (Robertson et al., 2010), Trinity (Grabherr et al., 2011), Oases (Schulz et al., 2012), and SOAP-denovo-Trans (Xie et al., 2014). An overview of current transcriptome assemblers is available, e.g., in Hölzer and Marz (2019). According to that study, the best performing assemblers were Trans-ABySS, Trinity and rnaSPAdes (Bushmanova et al., 2019).

In principle, *de novo* transcriptome assemblers can also reconstruct transcript isoforms of the expressed genes, but in practice their sensitivity is poor. In *Mus musculus*, Schulz et al. (2012) reported that Oases, Trans-ABySS, and Trinity assembled 1.21, 1.25, and 1.01 transcripts per gene, respectively, whereas a reference-based assembler reconstructed 1.56 transcripts per gene. Bushmanova et al. (2019) also observed poor transcript isoform reconstruction performance of transcriptome assembly methods: while their method, rnaSPAdes, outperformed the other compared assemblers in gene reconstruction in *Mus musculus*, it assembled only 1.02 transcripts per gene. In the same comparison, Trinity managed to assemble the most transcripts, with a ratio of 1.11 transcripts per gene. The insufficient ability of current *de novo* transcriptome assembly approaches to reconstruct all expressed transcript isoforms of a gene was evident to us in our work on the DAL19 gene in spruce, *Picea abies* (Akhter et al., 2018): Only one out of four confirmed DAL19 transcript isoforms was reconstructed to at least 90% using Oases and two using Trinity. We performed a directed assembly that managed to reconstruct three of the four transcript isoforms, but this method did not scale to whole transcriptome assembly. These examples, and others, e.g. Hayer et al. (2015) and Thind et al. (2021), demonstrate that there is still much room for improvement in *de novo* transcript isoform assembly.

Another observation concerns the imperfect overlap between the sets of reconstructed transcripts from different *de novo* assembly tools. Smith-Unna et al. (2016) noted that out of Oases, Trinity, and SOAP-denovo-Trans, each assembler reconstructed a large number of *bona fide* transcripts that neither of the other assemblers managed to reconstruct. They concluded that combining assembly methods may be an effective way to improve the detection rate of transcripts.

We report the *de novo* transcriptome assembler ClusTrast, which builds upon our previous experience of transcript isoform assembly (Akhter et al., 2018). The main purpose of ClusTrast is to provide a comprehensive set of transcript isoforms, using only sequence reads as input, and with the explicit intent to prioritize recall. The ClusTrast pipeline combines two assembly methods, Trans-ABySS and Shannon, incorporates a novel approach to clustering guiding contigs, assigns short reads to the clusters, and finally performs a clusterwise assembly of the clustered short reads. We assessed transcript isoform reconstruction performance of ClusTrast and several *de novo* transcriptome assemblers in six eukaryotic organisms and found that ClusTrast reconstructed more known transcript isoforms than any other assembler and reconstructed unknown (including misassembled) transcripts at a rate comparable to other assemblers. .

## 2 Implementation

### 2.1 ClusTrast method

We developed an approach for transcriptome assembly from short reads called ClusTrast.

#### 2.1.1 Overview

Figure 1 shows a flowchart of the ClusTrast pipeline. The only required input to ClusTrast is a file with short RNA-seq reads, referred to as SRs, short reads or SR RNA-seq. Guiding contigs (GCs) is an optional input. Supplementary Figure S.1 illustrates an example of how the method works.

**Figure 1:**
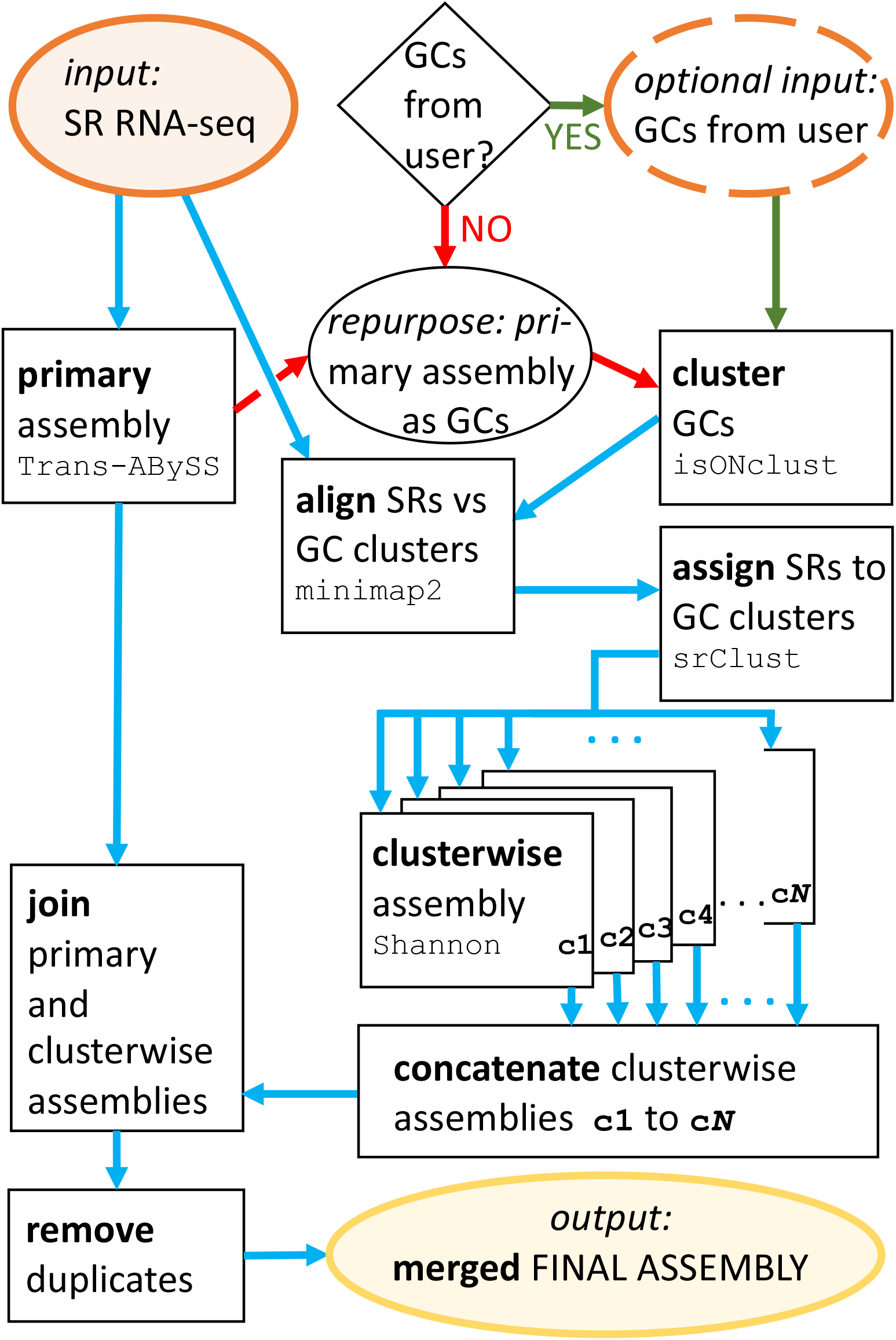
The ClusTrast pipeline. Orange ovals denote input data, black squares denote actions taken by ClusTrast, blue arrows denote intermediate data file transfers, yellow oval denotes output final assembly, and the black rhomboid and oval handle the guiding contig user input status. The only required input to ClusTrast is a short read RNA-seq data set (orange oval in top left corner). Guiding contigs may optionally be input by the user (dashed orange oval in top right corner, and green arrows followed if provided. They may also be used as primary assembly (not depicted)). If guiding contigs are not provided by the user, the primary assembly is repurposed and used also as guiding contigs (red arrows). SR = short reads, GCs = guiding contigs.

#### 2.1.2 Primary assembly and guiding contigs (GCs)

In ClusTrast, Trans-ABySS (Robertson et al., 2010) is employed to create a “primary assembly” from the short reads. The primary assembly will by default be used as the set of guiding contigs (GCs) in ClusTrast. The guiding contigs are used in the next step to assign the short reads to clusters (2.1.3). Guiding contigs may also be provided separately by the user, and could then also serve as primary assembly if desired. The primary assembly is by default merged into the final set of assembled transcripts (2.1.5).

Trans-ABySS is one of the leading *de novo* transcriptome assemblers according to Hölzer and Marz (2019). They evaluated the original strategy of Trans-ABySS, which used several different *k*-mers and merged the resulting assemblies, in order to get both recall from small values of *k* and precision from high values of *k*. Since a single-*k* run uses much less memory (or is substantially faster) than a multi-*k* run, we tried both strategies with ClusTrast. In this report, we have appended **-M** to the name of a method if it used a multi-*k* strategy. We also tried other assemblers as potential primary assemblers for ClusTrast (section 3.3).

#### 2.1.3 (a) Clustering of guiding contigs and (b) Assigning short reads to clusters

**(a)** The clustering of the guiding contigs is performed with isONclust (Sahlin and Medvedev, 2020), a tool originally developed for clustering of long reads (from PacBio or ONT sequencing technologies) into gene families. It uses a greedy algorithm for the clustering and handles variable error rates by the means of the quality values in the FASTQ input files. When the set of guiding contigs is in FASTA-format, ClusTrast will convert it to FASTQ-format with a static quality.

**(b)** The short reads are aligned to the guiding contigs with minimap2 (Li, 2018), using the preset option -x sr, intended for short read alignment, but included secondary alignments. Secondary alignments can optionally be excluded in ClusTrast. Next, the short reads are assigned to the guiding contig clusters based on the alignment results. If a short read *x* is aligned to guiding contigs *X*_1_ and *X*_2_, and *X*_1_ belongs to cluster *n*_1_ and *X*_2_ belongs to cluster *n*_2_, *x* will be included in *n*_1_ and *n*_2_. Reads not aligining to any guiding contig are put in a separate cluster. Thus, the short reads have now been clustered. A read can only occur once per cluster. See also Supplementary Section C.2.

#### 2.1.4 Clusterwise assembly

The clusterwise assembly in ClusTrast is performed by the transcriptome assembler Shannon (Kannan et al., 2016) (also used in refShannon, a genome-guided transcript assembler (Mao et al., 2020)), and aims to be information theoretically optimal. Kannan et al. claim that Shannon can finish in linear time given (*i*) sufficient diversity of transcript abundance and (*ii*) no loops in the graph, but they do not address how it will deal with datasets not meeting these criteria. However, dividing the reads in the short read dataset into clusters before assembly will reduce the complexity of each individual assembly and lower the risk of violating these requirements. Because of this, and its aim to reconstruct as many transcripts as possible, Shannon is used for the clusterwise assemblies in ClusTrast.

#### 2.1.5 Merging the primary and clusterwise assemblies

The final steps of ClusTrast are to join the concatenated clusterwise assemblies with the primary assembly, and next, to remove duplicate instances of reconstructed transcripts. Similar but non-identical reconstructed transcripts are kept, because of possible polymorphic variants that we want ClusTrast to retain (see also section 3.1.5). The output of ClusTrast is the merged final assembly.

### 2.2 Datasets and annotations

#### 2.2.1 Short read RNA-seq datasets for assembly generation

We evaluated the *de novo* transcriptome assemblies using the NCBI SRA datasets in Table 1. They were all non-stranded paired-end short read RNA-seq datasets. We pre-processed the datasets with fastp (Chen et al., 2018) with default parameters, which means removal of any remaining adapter sequences, quality pruning (max 40% of the bases were allowed to have base quality < 16, and at most five Ns per read), and exclusion of reads that ended up shorter than 15bp (see the supplementary material for details).

**Table 1:** Short read RNA-seq datasets accessed from the NCBI SRA database. RL=read length in bases. Species column, indicated in bold is the name to which the data set is referred to throughout this article. RPs=million read pairs, before pre-processing (on the left) and after pre-processing (on the right).

| SRA ID | RL | Species |  | RPs |  |
| --- | --- | --- | --- | --- | --- |
| SRR5133163.1 | $2 \times 150$ | **Human** | *Homo sapiens* | 29.51 | 29.05 |
| SRR8632985 | $2 \times 76$ | **Mouse** | *Mus musculus* | 31 | 30 |
| SRR11341576 | $2 \times 150$ | **Rice** | *Oryza sativa* | 24 | 23 |
| SRR11278019 | $2 \times 126$ | **Arabidopsis** | *Arabidopsis thaliana* | 11.2 | 11.1 |
| SRR10728575 | $2 \times 150$ | **Zebrafish** | *Danio rerio* | 21 | 20 |
| SRR5986240.1 | $2 \times 150$ | **Poplar** | *Populus trichocarpa* | 25.1 | 24.4 |

#### 2.2.2 Reference datasets for assembly evaluation

We downloaded reference genome sequences as well as reference transcript annotations from Ensembl for each of the six species. We used the GTF file annotations of genes and transcripts, not including the abinitio annotations. We estimated the expression of all reference transcripts in each of the six datasets using RSEM (Li and Dewey, 2011) and defined a transcript isoform as expressed if the transcripts per million (TPM) reported by RSEM was greater than zero. Versions and commands for RSEM are listed in the supplementary material. We defined a gene as expressed if at least one of the transcript isoforms associated with that gene was expressed. The versions of the annotations used and the number of genes and isoforms we detected in each dataset are shown in Table 2.

**Table 2:**
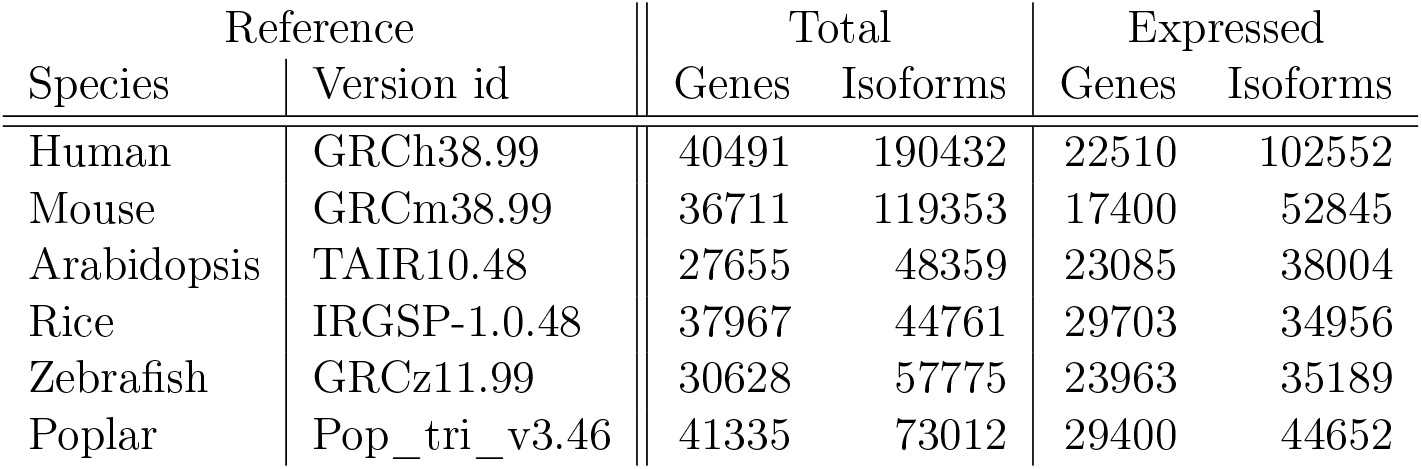
Reference transcriptome sequence, genome sequence and annotation versions accessed from Ensembl (https://www.ensembl.org/index.html and https://plants.ensembl.org/index.html for non-plant and plant species, respectively), where the Version id suffix shows the Ensembl version. The number of genes and isoforms are counted from the reference transcriptome. Genes and isoforms are considered expressed if TPM>0 as calculated by RSEM on the datasets in Table 1.

| Species | Reference<br>Version id | Total |  | Expressed |  |
| --- | --- | --- | --- | --- | --- |
|  |  | Genes | Isoforms | Genes | Isoforms |
| Human | GRCh38.99 | 40491 | 190432 | 22510 | 102552 |
| Mouse | GRCm38.99 | 36711 | 119353 | 17400 | 52845 |
| Arabidopsis | TAIR10.48 | 27655 | 48359 | 23085 | 38004 |
| Rice | IRGSP-1.0.48 | 37967 | 44761 | 29703 | 34956 |
| Zebrafish | GRCz11.99 | 30628 | 57775 | 23963 | 35189 |
| Poplar | Pop_tri_v3.46 | 41335 | 73012 | 29400 | 44652 |

### 2.3 Transcriptome assembly generation

We assembled the transcriptomes for all six datasets (Table 1) using Trans-ABySS (Robertson et al., 2010), Trinity (Grabherr et al., 2011), Oases (Schulz et al., 2012), SOAP-denovo-Trans (Xie et al., 2014), BinPacker (Liu et al., 2016), Shannon (Kannan et al., 2016), rnaSPAdes (Bushmanova et al., 2019), TransLiG (Liu et al., 2019), RNA-Bloom (Nip et al., 2020), and ClusTrast. We used each assembler’s own default parameters. Trans-ABySS and Oases can be run in a “multi-*k*” mode where the assembler is first run with a single *k*-mer (“single-*k*” mode; where a *k*-mer is a substring, with fixed length *k*, of a read) for several different *k*-mers and the resulting assemblies are merged into a single assembly. We used both the single-*k* and multi-*k* strategies for these two assemblers. We append **-M** to the name of a method if it uses a multi-*k* strategy, and **-S** if it uses a single-*k* strategy. Oases-**M** uses by default all odd *k*-mers from 19 to 31, but it only finished within less than 58 hours on the mouse and arabidopsis datasets. On the rice dataset, it finished after ∼400 hours. Therefore, for human and zebrafish, we used only Oases-**S** with *k* = 31. The program versions and the executed commands are listed in the supplementary material.

We also generated a concatenated assembly from the Trans-ABySS and Shannon transcrip-tomes, referred to as TrAB+Sh, to examine if the clustering approach of ClusTrast improves the assembly quality.

### 2.4 Transcriptome assembly evaluation

We evaluated the transcriptome assemblies by estimating precision (positive predicted value, PPV) and recall (sensitivity or true positive rate, TPR). For this, we used the reference based transcriptome comparison tools Conditional Reciprocal Best BLAST (CRBB) (Aubry et al., 2014), as implemented in the TransRate package (Smith-Unna et al., 2016), and SQANTI (Tardaguila et al., 2018). Versions and commands for these tools can be found in the supplementary material. We only used reference transcripts that were considered expressed (2.2.2). All assembled transcripts were considered expressed, since they were reconstructed from actual RNA-seq data.

#### 2.4.1 Using SQANTI in evaluation

We used SQANTI (Structural and Quality Annotation of Novel Transcript Isoforms) (Tardaguila et al., 2018) to classify assembled transcripts according to their splice junction matches with reference genes and transcript isoforms. When an assembled transcript is anti-sense to an an-notated gene, SQANTI will classify that transcript as anti-sense. We extracted all transcripts classified as anti-sense, reverse-complemented them, and then reclassified them with SQANTI. When an assembled transcript and a reference isoform have the same number of exons and same splice junctions, then SQANTI classifies it as a full splice match (FSM). When the assembled transcript has fewer exons than the reference but the splice junctions in the assembled transcript all exist in the reference, it is classified as an incomplete splice match (ISM). In order for SQANTI to classify an assembled transcript as an ISM, all junctions in the assembled transcript must match the reference, but the exact start and end can differ. In case there are several possible consistent reference isoforms, SQANTI assigns the assembled transcript to the shortest of the matching references. Assembled transcripts classified by SQANTI as novel in catalog (NIC, when the splice junctions are known but there is a novel combination) and novel not in catalog (NNC, with novel splice junctions) were not classified as true positives.

For recall, we counted all expressed reference isoforms with at least one assembled transcript that SQANTI classified as FSM or ISM (and with a certain fraction, 0.25-1.0, of the exons covered) as a true positive, and divided the total number of true positives by the total number of expressed reference isoforms. For precision, we counted each assembled isoform classified by SQANTI as FSM or ISM and covering at least a certain fraction (from 0.25 to 1.0) of the reference exons as a true positive, and divided the total number of true positives by the total number of assembled isoforms.

#### 2.4.2 Using CRBB in evaluation

We used CRBB (Conditional Reciprocal Best BLAST) (Aubry et al., 2014) to classify assembled transcripts according to their similarity to reference transcripts. To this end, we used TransRate (Smith-Unna et al., 2016), which in turn used BLAST (Altschul et al., 1990) to align each assembled transcript to the set of reference transcripts, and each reference transcript to the set of assembled transcripts. By using all transcripts which are top hits in both BLAST alignments reciprocally, an appropriate E-value cutoff is calculated. Transcripts with lower E-values than this cutoff are then considered CRBB hits. We defined recall as the proportion of reference transcripts that have a CRBB hit covering the reference transcript to at least 25%–100%. We defined precision as the proportion of assembled transcripts that are a CRBB hit covering the reference isoform to at least 25%–100%.

## 3 Results

### 3.1 Transcriptome assembly evaluation

Transcriptome assemblies for all compared assemblers, including ClusTrast, were generated as described in 2.3. Basic statistics of all assembled transcriptomes are available in Supplementary Table S.2–S.7. We collectively refer to all tested approaches as “assemblers”, although assembly pipeline (e.g., ClusTrast) or concatenation (TrAB+Sh) may be more accurate.

#### 3.1.1 Evaluation with SQANTI

We investigated how recall and precision changed when we varied the proportion of exons that an assembled transcript needs to recover in order to be considered a true positive. As this proportion was relaxed for the ISM classifications from 1.0 to 0.25 (for the FSM category it is by definition 1.0), the recall (Figure 2) and precision (Figure 3) increased. ClusTrast-**M** had the highest SQANTI recall of any assembler for all of the six datasets over the entire range (except roughly tied with TrAB+Sh for arabidopsis). The assembler with the highest precision varied across datasets; it was RNA-Bloom in human, ClusTrast-**M** in rice and (with TransLiG) poplar, Oases-**M** in mouse, and TransLiG in arabidopsis and zebrafish.

**Figure 2:**
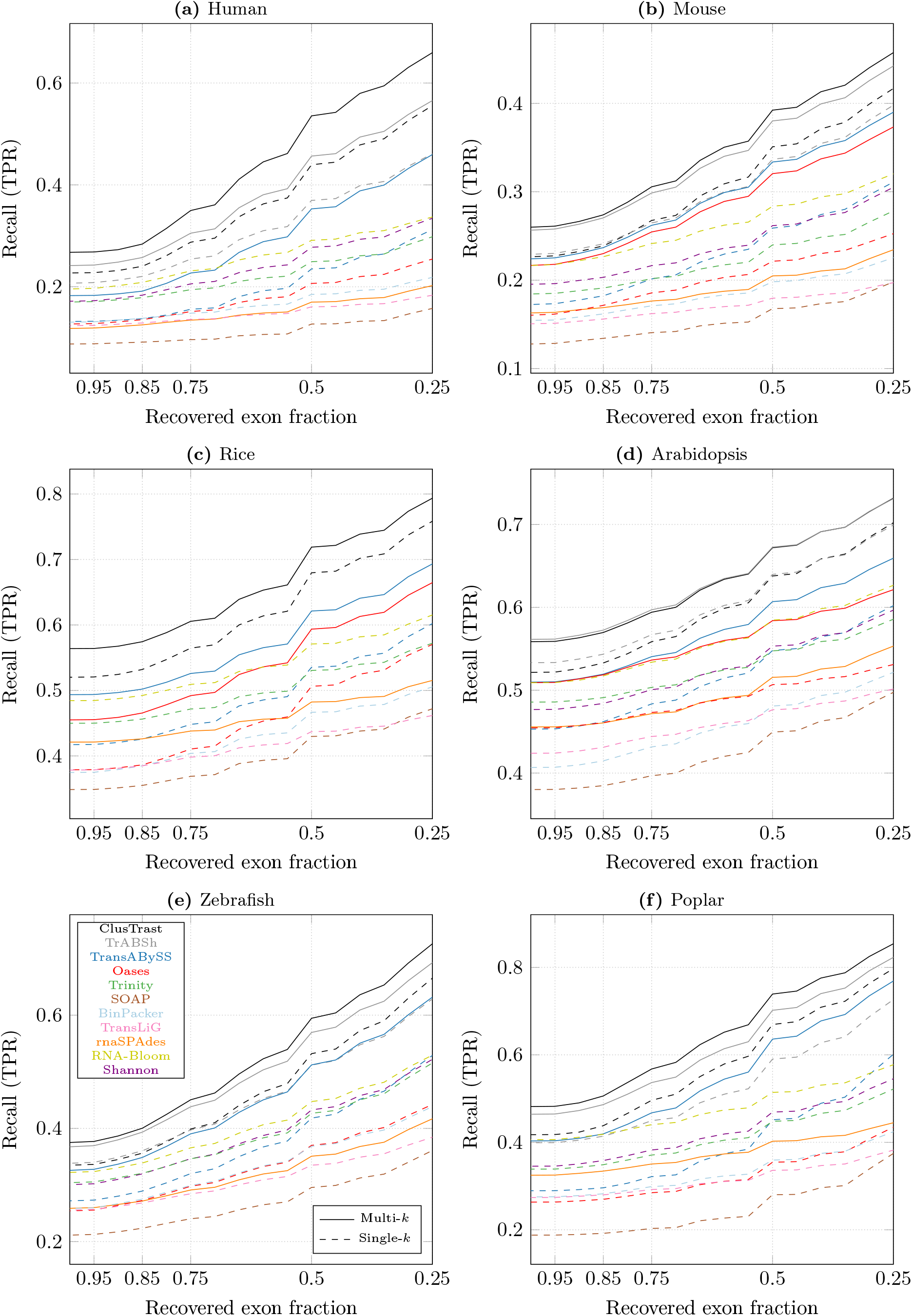
Proportion of reference isoforms with at least one SQANTI classification of FSM or ISM vs. the cumulative proportion of exons recovered by the assembly.

**Figure 3:**
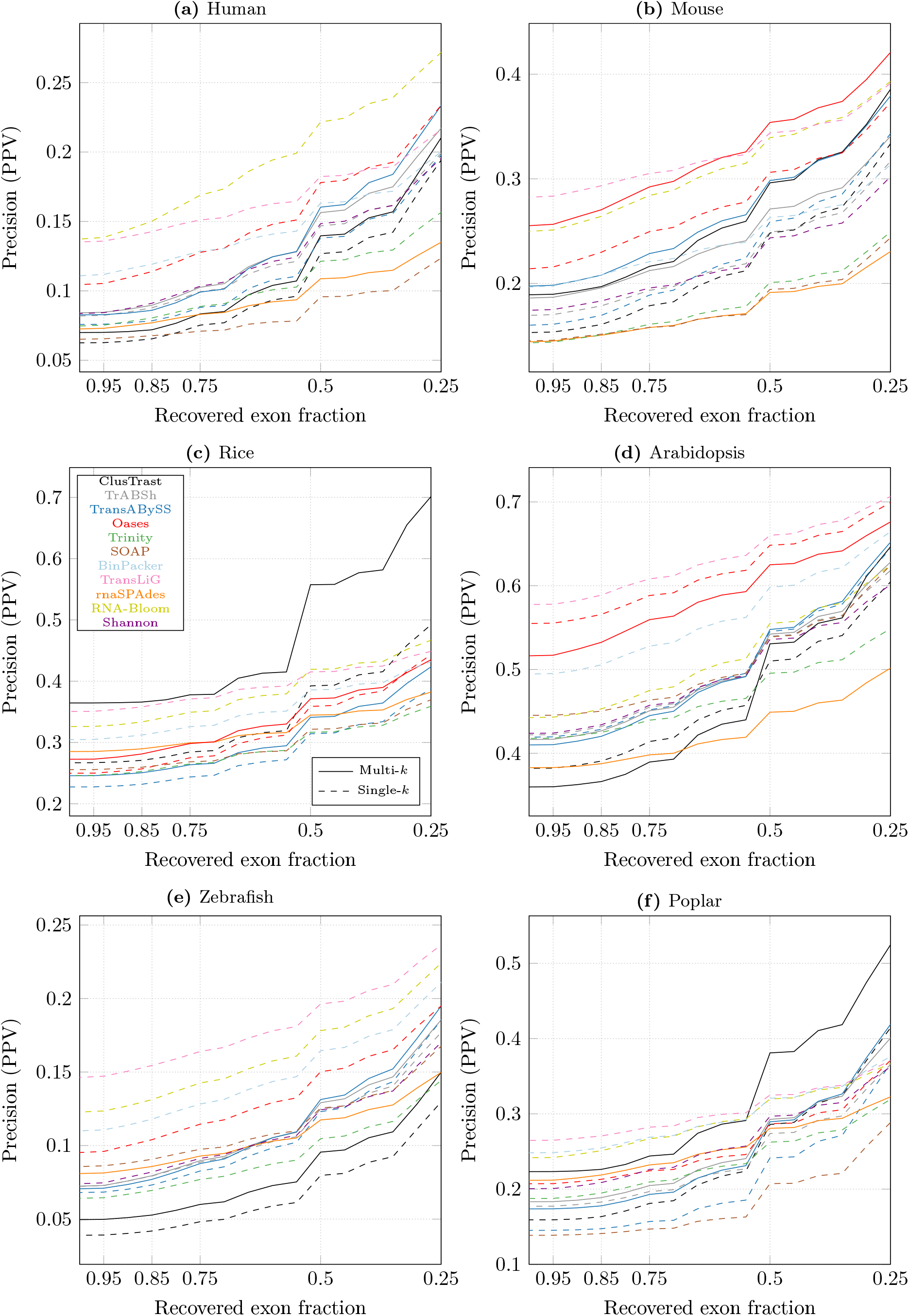
Proportion of reconstructed isoforms classified by SQANTI as FSM or ISM vs. the cumulative proportion of recovered exons from the reference.

Fixing the proportion at 0.5 (i.e., at least 50% of exons recovered for ISM), ClusTrast-**M** detected more transcript isoforms than the established assemblers Trinity (1.23–2.15 fold increase), Oases-**S**(1.33–2.59 fold increase), and Trans-ABySS-**M**(1.1–1.52 fold increase) (Supplementary Figure S.2 and Supplementary Table S.14). Precision was comparable to Trinity (0.9–1.76 fold change), Oases-**S**(0.78–1.5 fold change) and Trans-ABySS-**M**(0.73–1.63 fold change).

#### 3.1.2 Evaluation with CRBB

We investigated CRBB recall and precision over the same proportion of required recovered exons as for SQANTI and observed an increase in recall and precision as this proportion was decreased from 1.0 to 0.25. We observed some changes in the relative ordering of assemblers as shown in Figure 4 (CRBB recall) and Figure 5 (CRBB precision). In particular, rnaSPAdes performance levelled off in the lower end.

**Figure 4:**
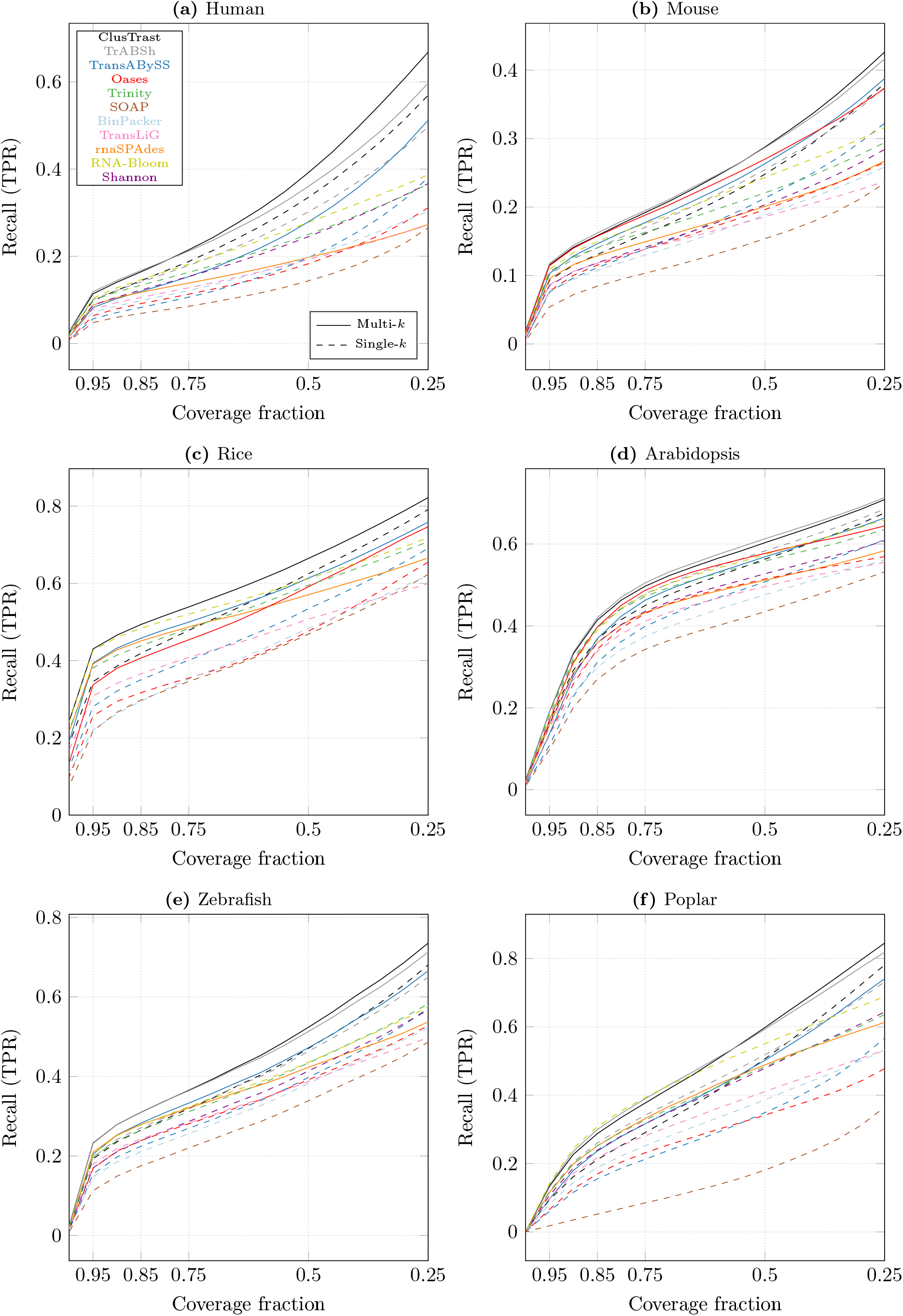
Proportion of references with a CRBB hit vs. the cumulative proportion of recovered reference length.

**Figure 5:**
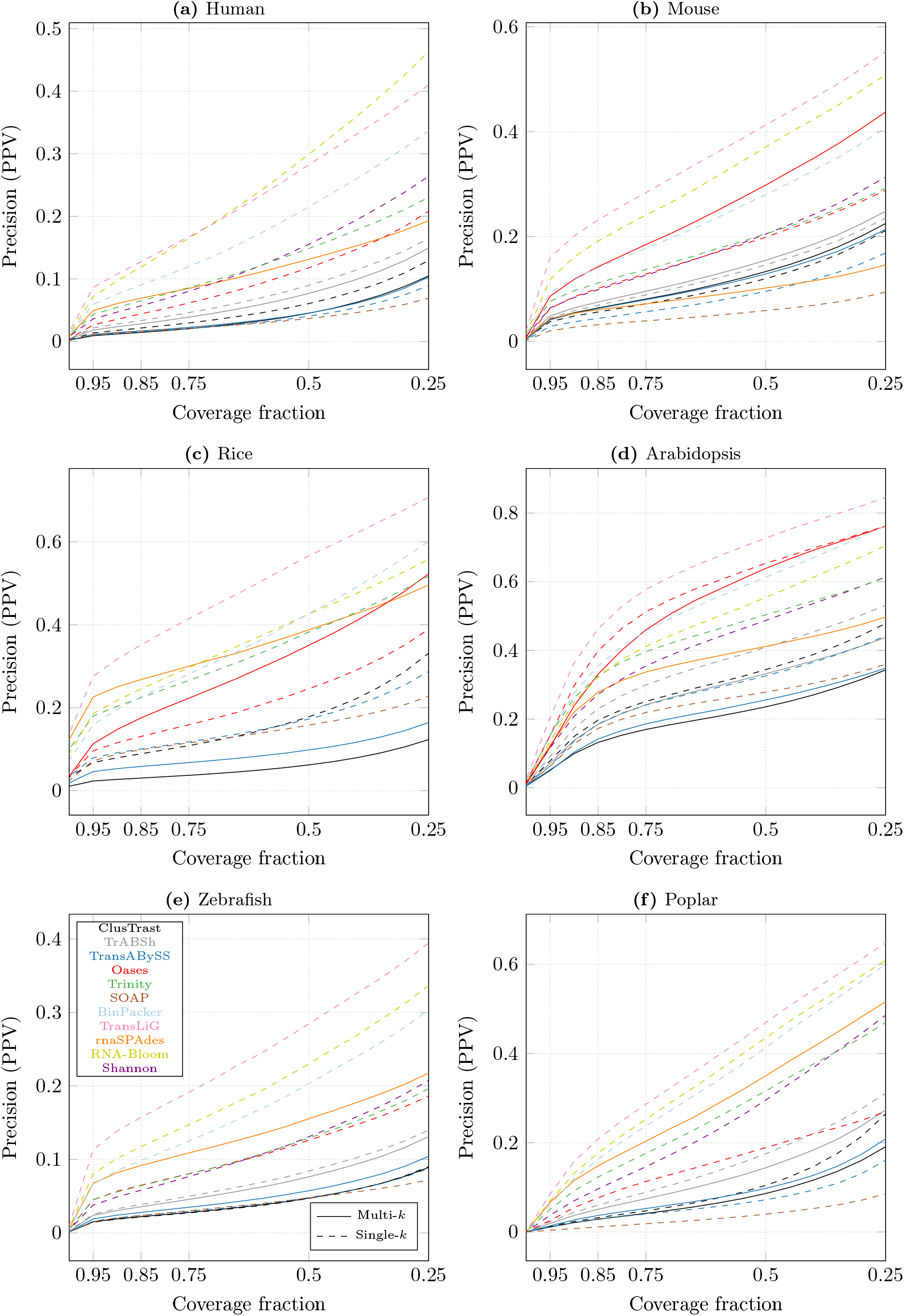
Proportion of reconstructed isoforms with a CRBB hit vs. the cumulative proportion of recovered reference length.

Fixing the proportion at 0.5, CRBB recall was higher for ClusTrast-**M** than for Trinity (1.01–1.34 fold increase) across all datasets, but not compared to all assemblers (Supplementary Figure S.3 and Supplementary Table S.15). ClusTrast-**M** performed the best on human, mouse, rice, and zebrafish, it ranked second for arabidopsis, while its ranking varied from first to third as the required reference transcript coverage decreased. ClusTrast-**M** clearly underperformed compared with Trinity with regard to CRBB precision (0.33–0.68 fold change); the assembler with highest precision in a dataset was always TransLiG or RNA-Bloom.

#### 3.1.3 SQANTI and CRBB evaluation metrics were correlated; true positive sets still differed

The number of transcripts that were considered as true positives by both SQANTI and CRBB or exclusively by only one of them varied between datasets and between assemblers (Supplementary Tables S.18 and S.19). ClusTrast, Trans-ABySS, Oases-**M**, and (with one exception) Shannon and SOAP-denovo-Trans consistently predicted more transcripts that were considered as true positives exclusively by SQANTI and not by CRBB, while TransLiG was the only assembler that consistently predicted more transcripts that were considered true positives exclusively by CRBB. We used SQANTI categories to classify the ClusTrast true positives that were exclusively detected by either SQANTI or CRBB (Supplementary Tables S.23 and S.24, respectively). We observed that the largest category of true positives according to SQANTI but not CRBB was the ISM mono-exon class. The largest category of true positives according to CRBB but not SQANTI was novel not in catalog (NNC) with novel splice sites.

SQANTI and CRBB recall measurements were highly correlated across all assemblies and datasets (*ρ* = 0.93; Supplementary Figure S.4) while SQANTI and CRBB precisions were less correlated (*ρ* = 0.75; Supplementary Figure S.5). We calculated the correlation of precision measurements for each assembler individually: ClusTrast-**M** obtained *ρ* = 0.54 while for all other assemblers *ρ* ≥ 0.82. Next, we excluded the ISM mono-exon class from the set of true positives and recalculated the precision correlation for ClusTrast-**M**: it increased to *ρ* = 0.94.

#### 3.1.4 Reference transcripts were often covered to at least 95% by FSMs

We investigated the number of expressed reference transcript isoforms that were reconstructed to at least 50% and 95% of their length by a single FSM according to SQANTI, Supplementary Table S.16. For all assemblies and both length requirements, either TrAB+Sh or ClusTrast reconstructed the most reference transcript isoforms, with small differences (<5%) except for rice where TrAB+Sh did not produce a result. Between 35.1% (arabidopsis) and 68.8% (rice) of the reference transcript isoforms that had an FSM match were reconstructed by the FSM-classified contig from ClusTrast to at least 95% of their length. The corresponding range for reconstruction to at least 50% of the reference transcript length was between 76.7% (human) and 91.0% (arabidopsis).

#### 3.1.5 An appreciable fraction of reference transcripts were reconstructed with polymorphisms by ClusTrast

We used the subset of reference transcripts with FSM or CRBB hits to estimate how often these reference transcripts were reconstructed as polymorphic variants (SNPs, indels) or as alternatively spliced contigs. In Supplementary Tables S.20 and S.21, the sets labeled *A* contain the FSMs, while the sets labeled *B* contain the CRBB hits. By definition, FSM contigs corresponding to a specific reference transcript are not alternatively spliced, since they contain all splice junctions of their reference transcript. Two (or more) FSM contigs matching one and the same reference transcript are thus polymorphic variants of each other. This is the *A ∖ B* and *A ⋂ B* sets in Supplementary Tables S.20 and S.21. On the other hand, two (or more) contigs that are not FSMs but considered as CRBB hits to one and the same reference transcript, are potentially splice variants of that reference transcript. This is the *B ∖ A* sets. We estimated that 58–81% of the reference transcripts reconstructed by ClusTrast were reconstructed with polymorphic variants, Supplementary Table S.22. Conversely, we estimated that 47–78% of ClusTrast assembled contigs contained polymorphic variants, Supplementary Table S.22.

#### 3.1.6 Recall varied over expression levels and number of exons in isoforms

To determine if the assemblers differed in how well they recovered isoforms of genes with more than one annotated isoform, we calculated SQANTI recall of isoforms binned by genes according to the number of isoforms these genes expressed (Figure 6). In most cases the ranking of assemblers by recall did not change with increasing number of expressed isoforms per gene. ClusTrast-**M** came out on top over almost the entire range for 5 out of 6 datasets, although for mouse it was tied with TrAB+Sh and for arabidopsis it was tied with Oases-**M** and TrAB+Sh. Next, we binned reference transcripts by expression quantiles as measured by RSEM. SQANTI recall increased with increased expression level, for all assemblers and for all data sets except that some assemblers levelled off in the range 80-100%. We observed that recall was higher for ClusTrast-**M** than all other assemblers in the lower end of expression levels (expression quantile <15%) and across the entire range of expression levels for all datasets except arabidopsis and zebrafish where ClusTrast-**M** was tied with TrAB+Sh (Figure 7).

**Figure 6:**
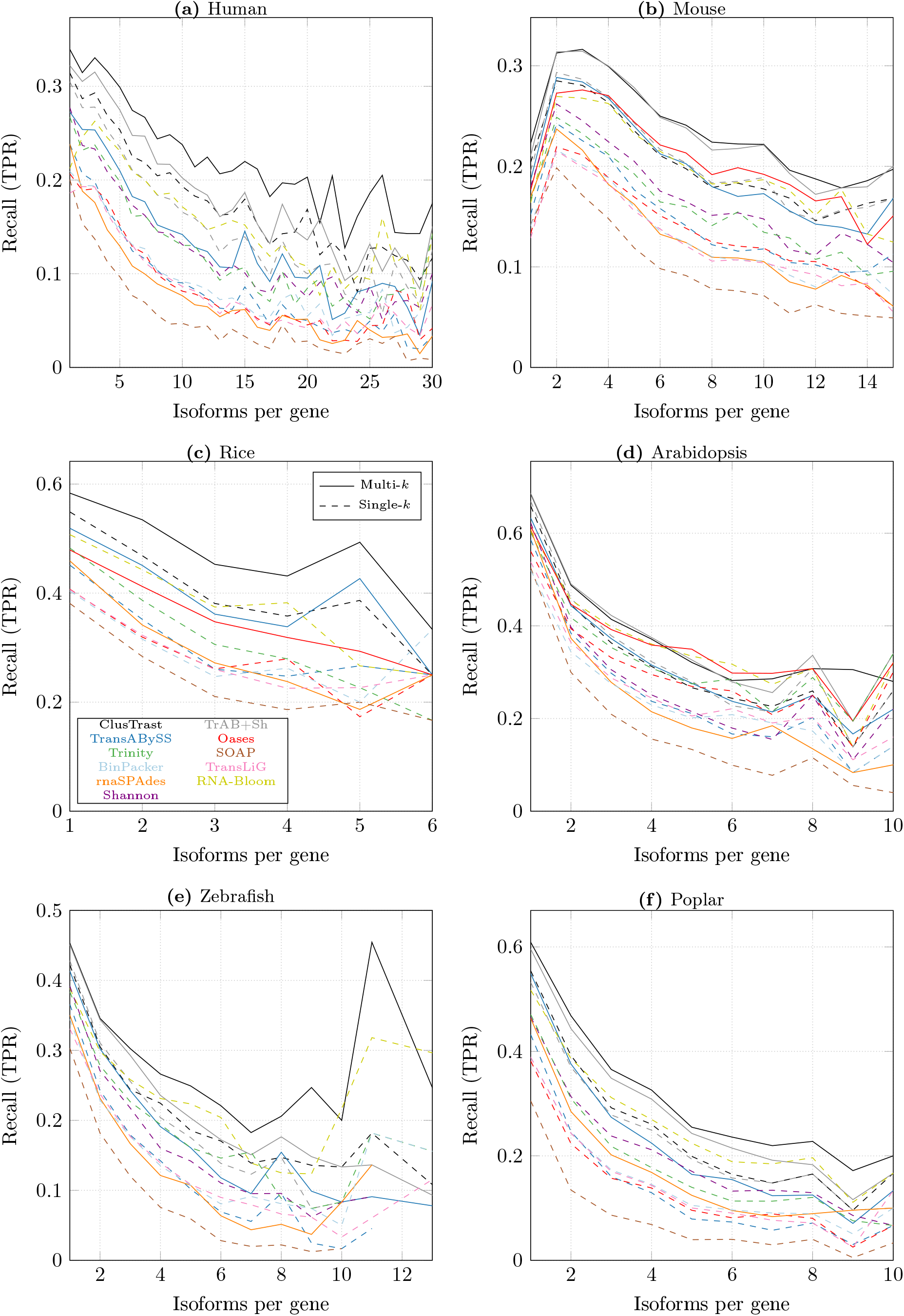
SQANTI recall of reference isoforms (FSM) binned by number of expressed isoforms per gene.

**Figure 7:**
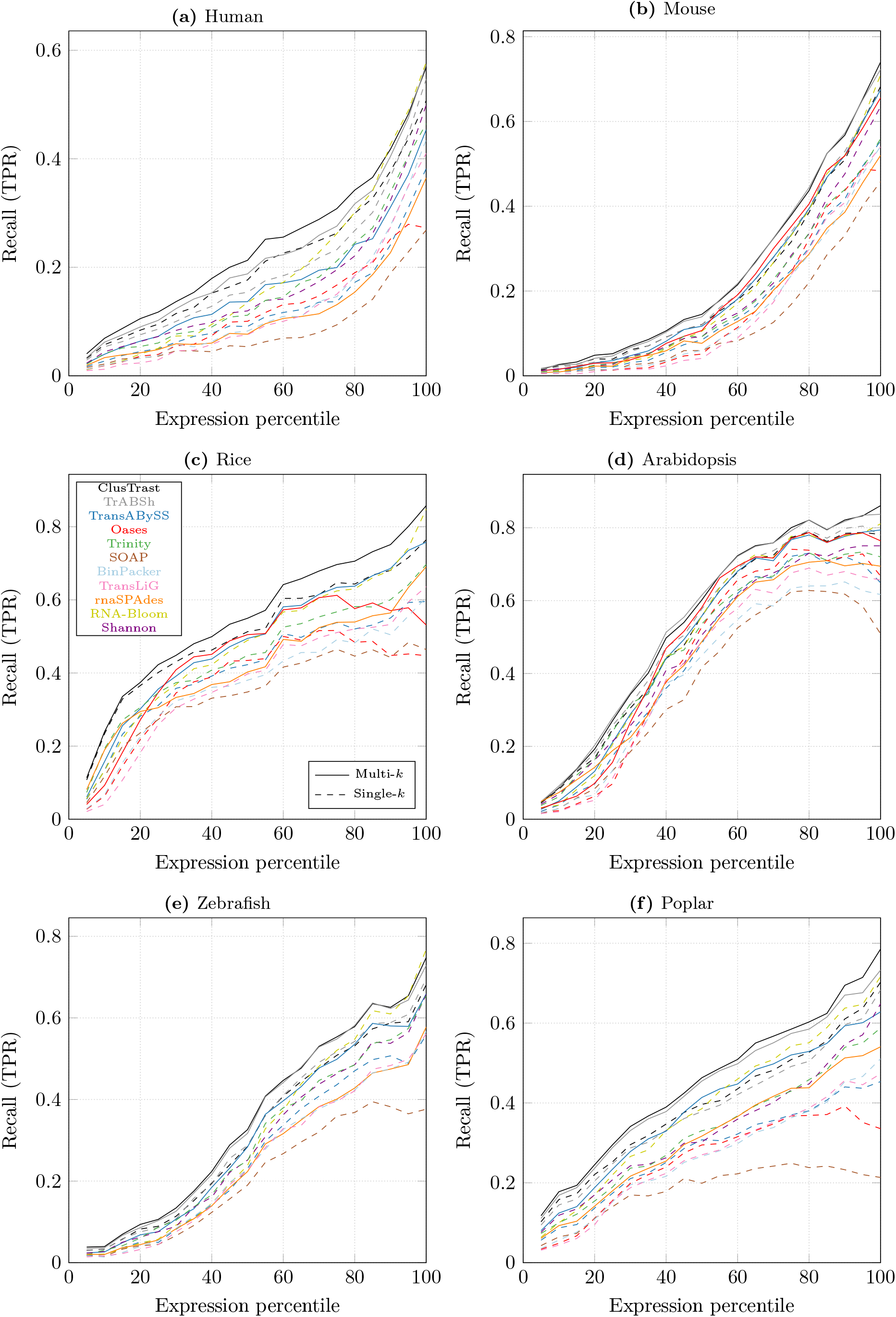
SQANTI recall of expressed transcript isoforms stratified according to RSEM expression. Recall within each bin (5 percentiles) is defined as the proportion of transcript isoforms that have an FSM match to an assembled contig that is ≥ 200bp.

We observed cases where ClusTrast detected highly expressed isoforms missed by other methods. We illustrate this with examples of genes where the highest expressed isoform (according to RSEM) was reconstructed only by ClusTrast and not by any other method. Supplementary Figures S.26 to S.30 contain Sahimi plots for these example genes, one for each of the six datasets (see Supplementary Section C.4 for more details).

#### 3.1.7 Simulated datasets

We evaluated ClusTrast on two simulated datasets: one human dataset from Hölzer and Marz (2019) and one mouse dataset from Hayer et al. (2015). The results (Figures 8 and 9) were analogous to the ones from the real datasets: On the simulated human dataset, ClusTrast was the leading method for recall (according to both SQANTI and CRBB) and the worst for precision (according to SQANTI). On the simulated mouse dataset, it was even between ClusTrast and TrAB+Sh on recall, and a mediocre to low precision for ClusTrast.

**Figure 8:**
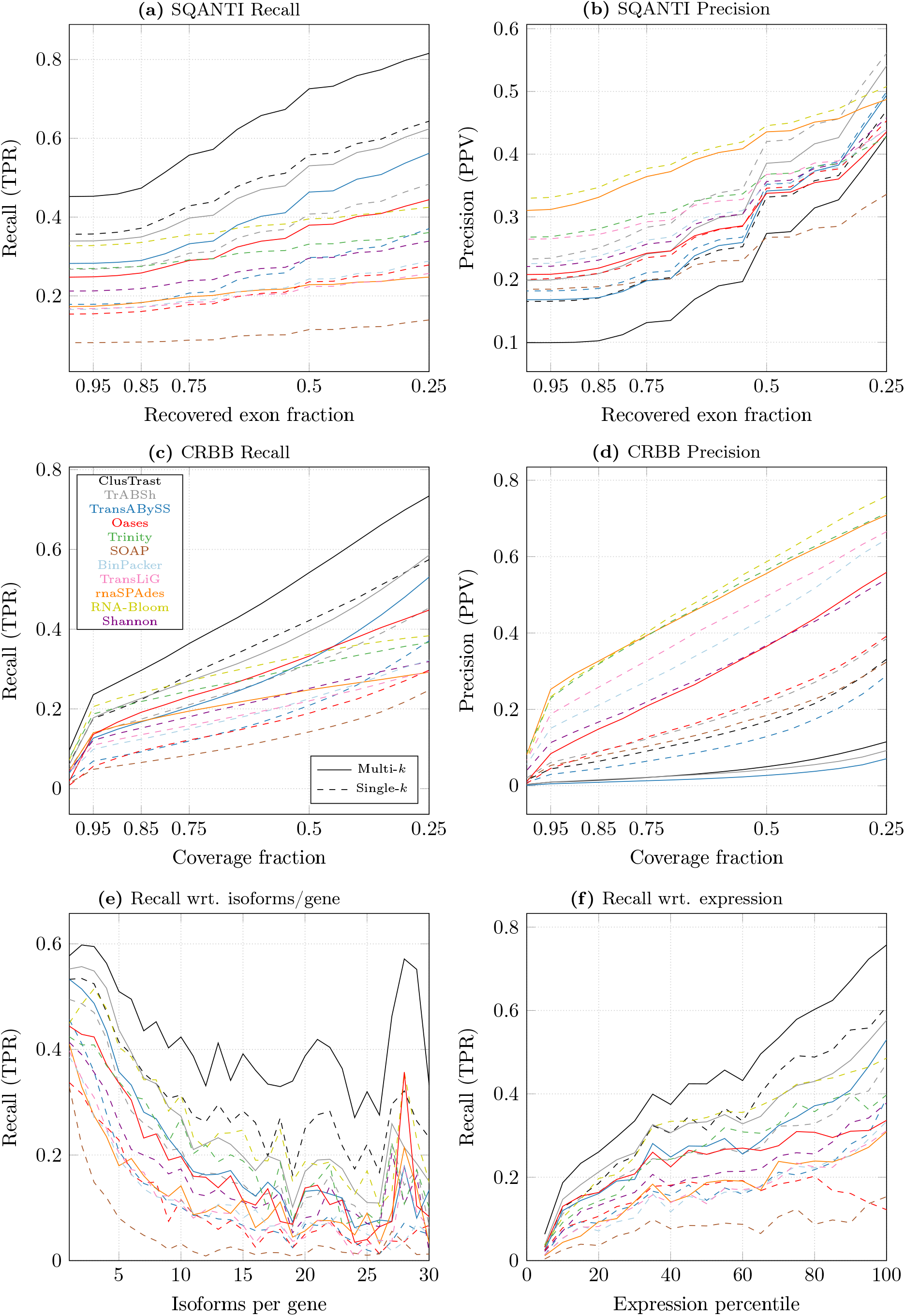
Results for a simulated human dataset from Hölzer and Marz (2019).

**Figure 9:**
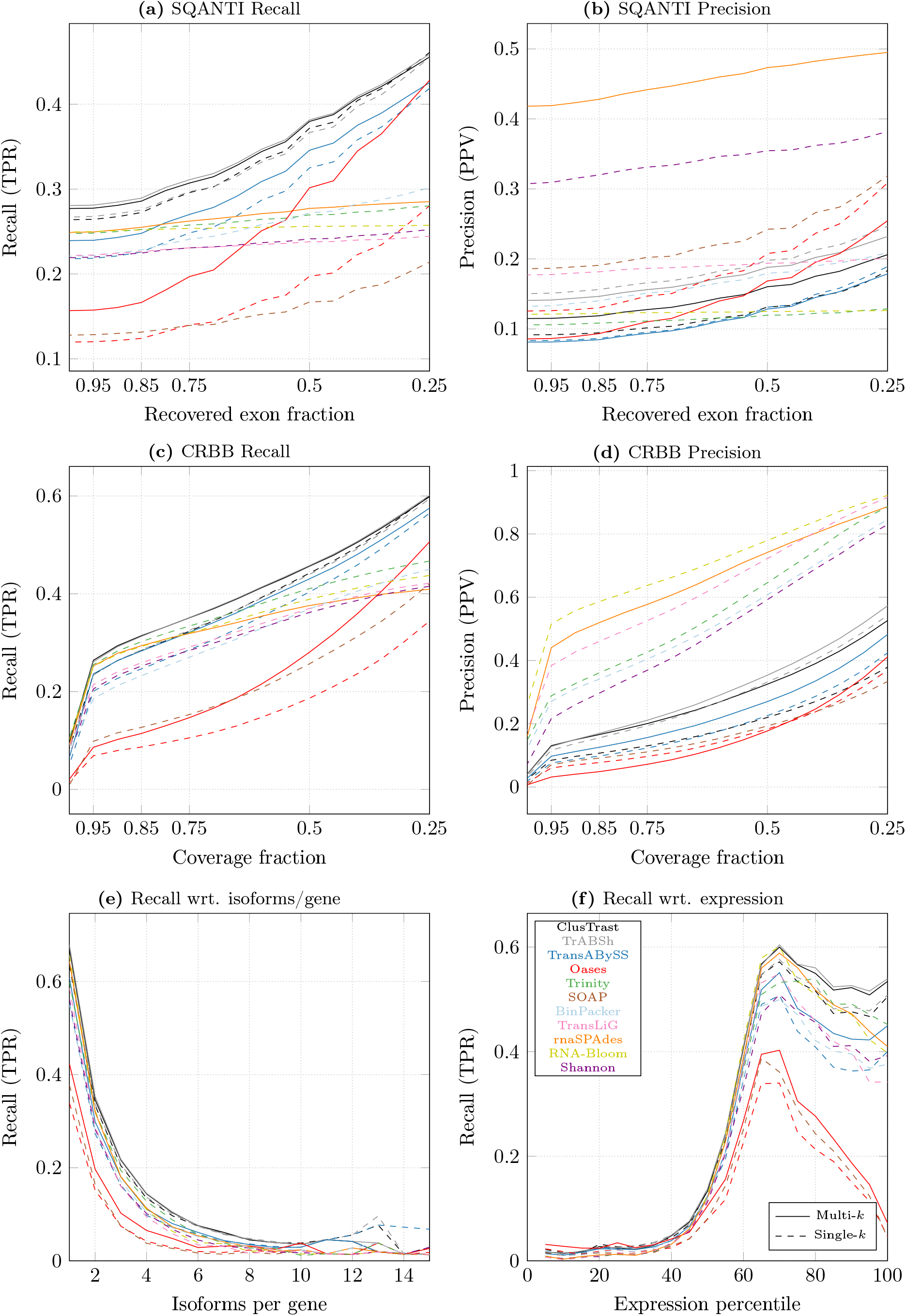
Results for a simulated mouse dataset from Hayer et al. (2015).

### 3.2 Run time and memory usage

Across all real datasets, all assemblers except Oases-**M** completed within 48 hours and required less than 300 GB of memory (Supplementary Table S.25). ClusTrast-**M** took between 660 and 2145 minutes to complete, the longest time of all assemblers for mouse and arabidopsis. Oases-**M** took the longest time to complete for one dataset, and did not complete for the remaining three. SOAP-denovo-Trans was the fastest assembler for all datasets. ClusTrast-**M** peak memory use was between 57.15 and 267.2 GB for the six datasets, highest of all assemblers for one dataset, while Trinity and Oases-**M** had the highest peak memory use for two datasets each. RNA-Bloom had the lowest peak memory usage. Our computational setup is described in Supplementary Table S.26.

### 3.3 Primary assembly alternatives

We assessed alternatives to Trans-ABySS-**M** for primary assembly in ClusTrast: the widely used Trinity, and the recent RNA-Bloom (both of which faring well individually in our own evaluation), and a “meta assembler”, that we call META, constructed by taking the union of the individual assemblies from Trans-ABySS-**M**, Trinity, and RNA-Bloom (Figures 10 - 13). ClusTrast was tested in four versions, each with a different primary assembly, Trans-ABySS-**M**, Trinity, RNA-Bloom, and META (solid lines). Recall was improved as compared to the four individual primary assemblies (dashed lines), according to both SQANTI, Figure 10, and CRBB, Figure 12, while precision was lower for 4/6 datasets according to SQANTI, Figure 11, and overall according to CRBB, Figure 13. We also concatenated the individual assembly from Trans-ABySS-**M**, Trinity, and RNA-Bloom, respectively, to the individual assembly from Shannon (dotted lines). These concatenated assemblies (where the concatenation of Trans-ABySS-**M** and Shannon, TrAB+SH, is included also in section 3.1) yielded higher recall for RNA-Bloom and Trinity as compared to what ClusTrast reached, while ClusTrast demonstrated higher recall for Trans-ABySS-**M** and META.

**Figure 10:**
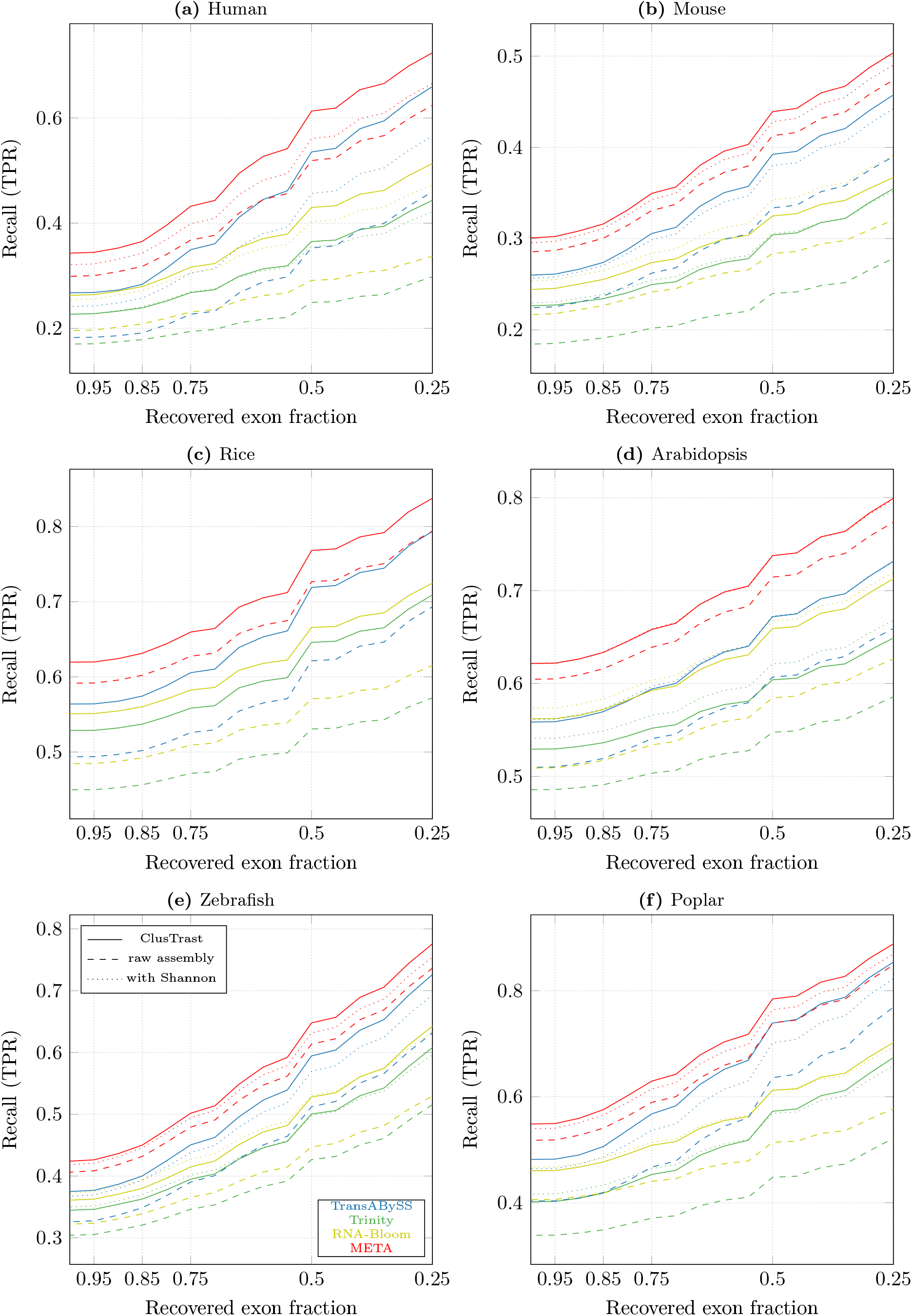
Proportion of reference isoforms with at least one SQANTI classification of FSM or ISM vs. the cumulative proportion of exons recovered by the assembly, for ClusTrast when ran with different tools for primary assembly (solid lines), compared with these tools on their own (dashed lines) and concatenated with Shannon (dotted lines).

**Figure 11:**
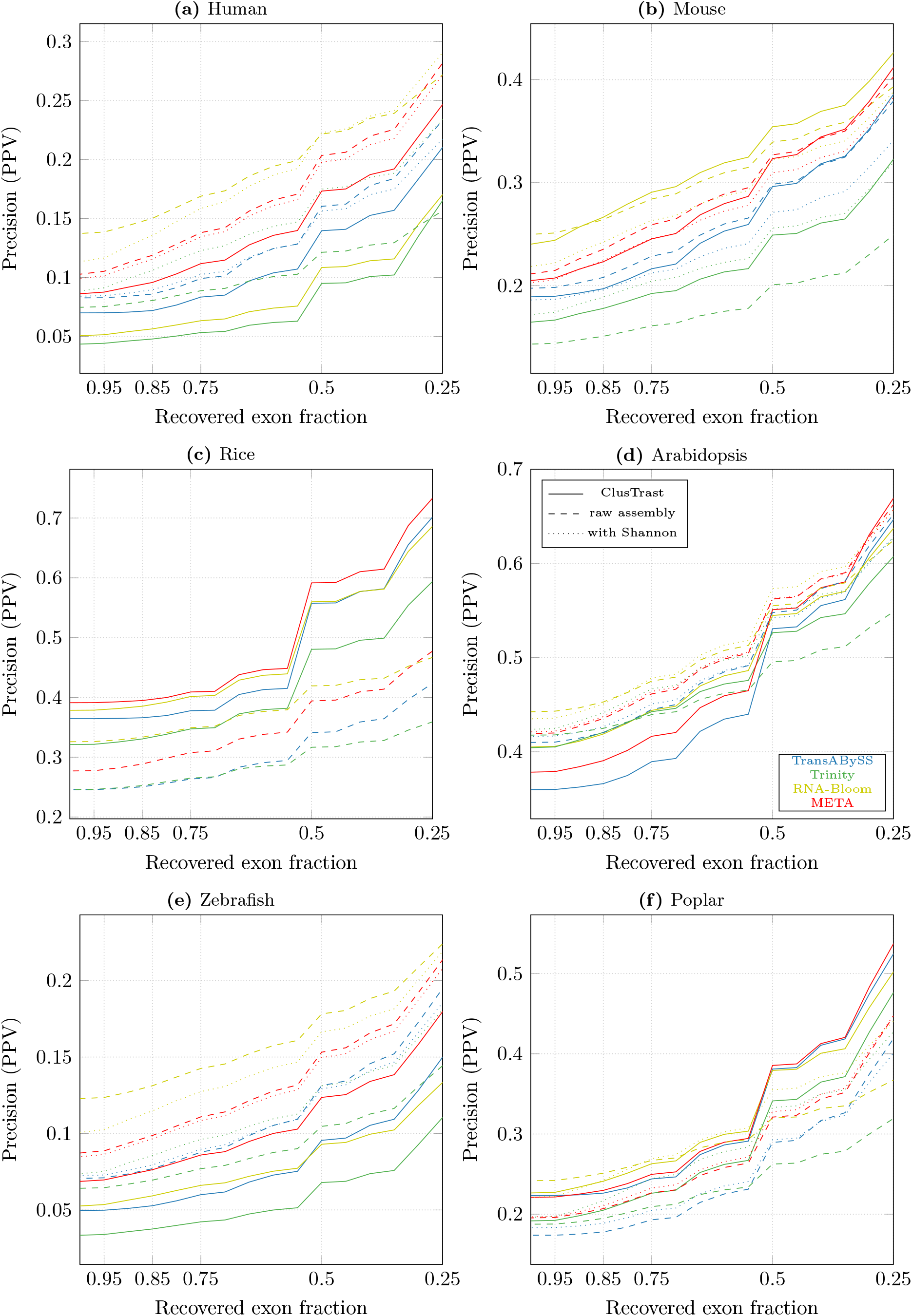
Proportion of reconstructed isoforms classified by SQANTI as FSM or ISM vs. the cumulative proportion of recovered exons from the reference, for ClusTrast when ran with different tools for primary assembly (solid lines), compared with these tools on their own (dashed lines) and concatenated with Shannon (dotted lines).

**Figure 12:**
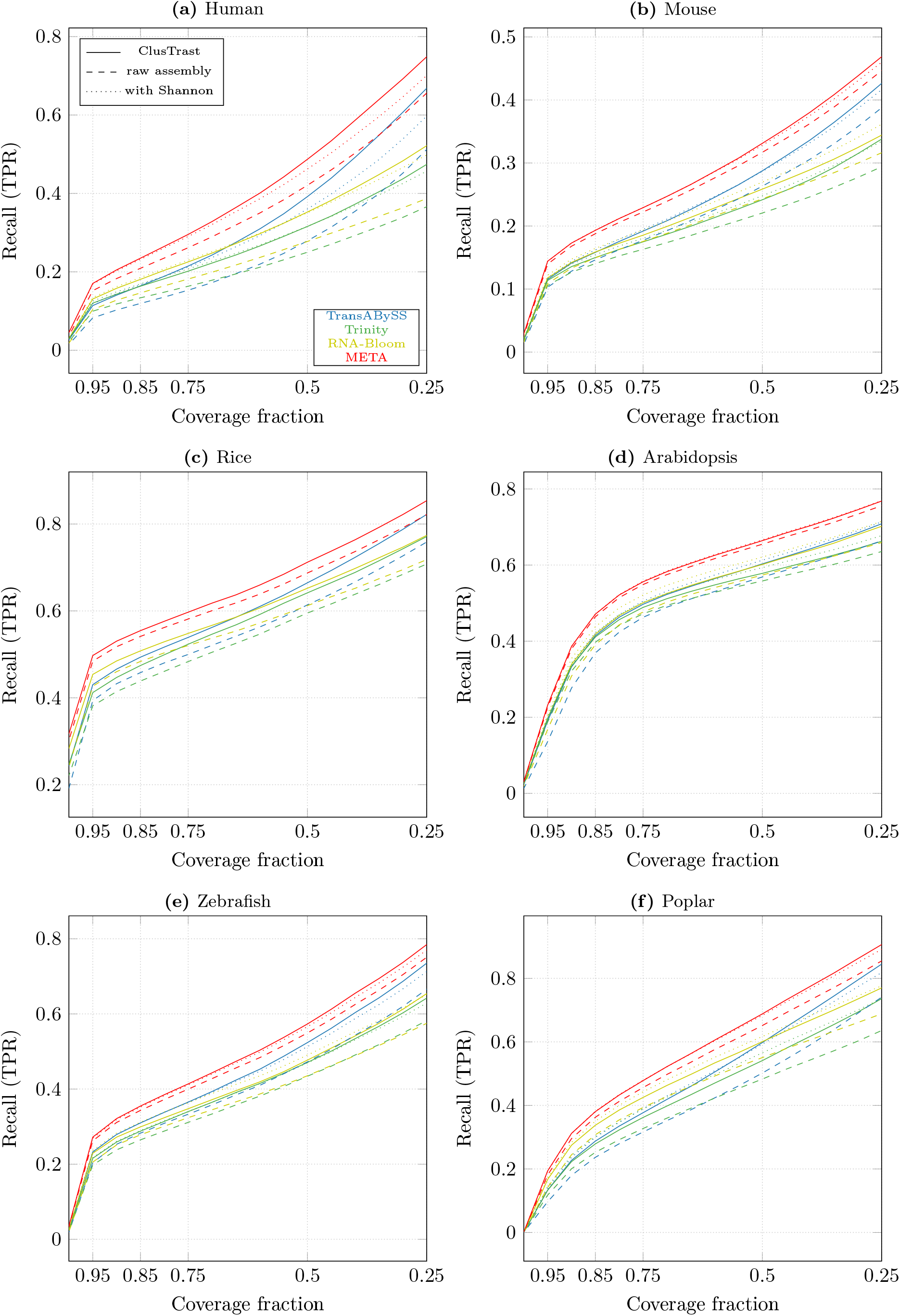
Proportion of references with a CRBB hit vs. the cumulative proportion of recovered reference length, for ClusTrast when ran with different tools for primary assembly (solid lines), compared with these tools on their own (dashed lines) and concatenated with Shannon (dotted lines).

**Figure 13:**
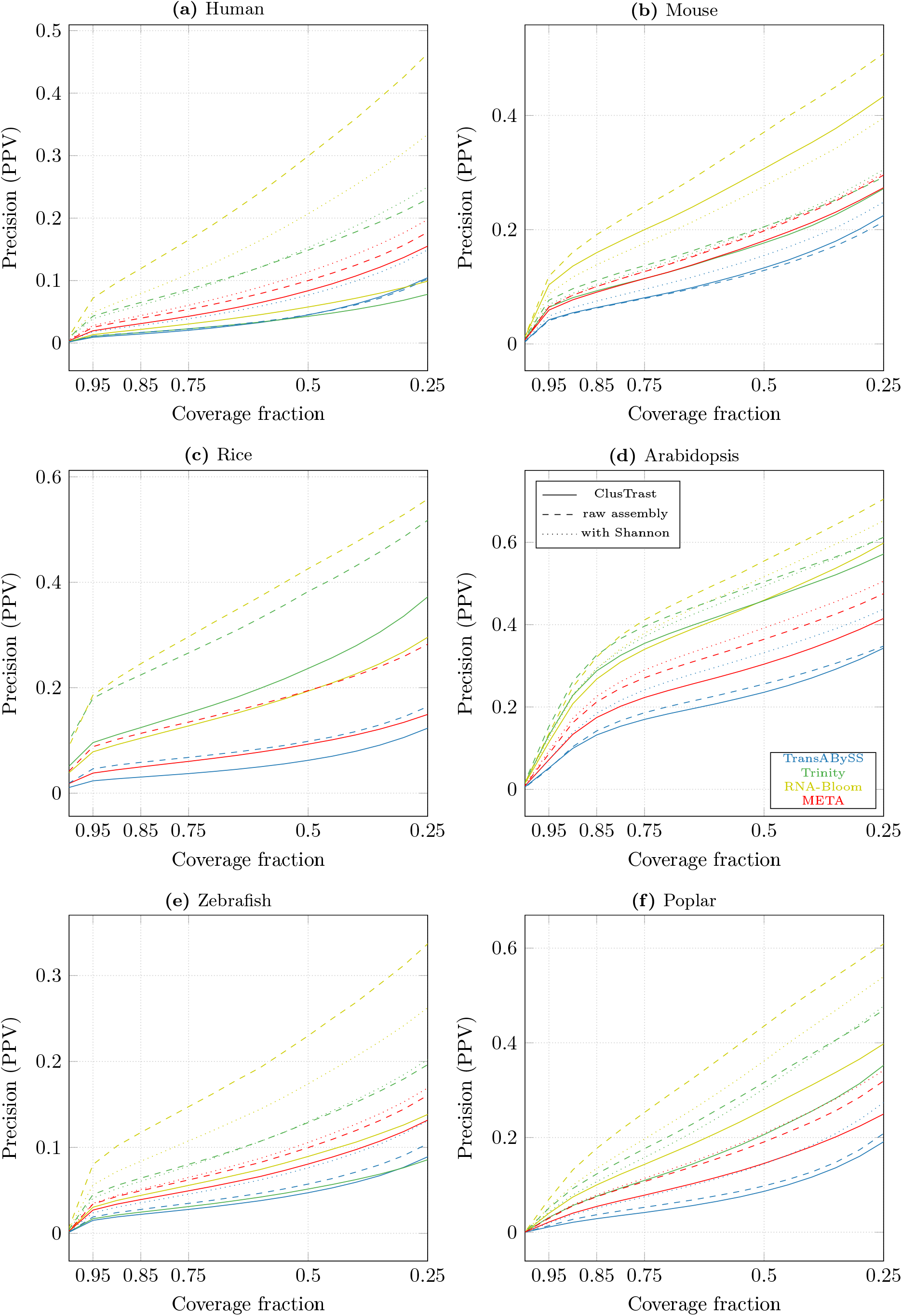
Proportion of reconstructed isoforms with a CRBB hit vs. the cumulative proportion of recovered reference length, for ClusTrast when ran with different tools for primary assembly (solid lines), compared with these tools on their own (dashed lines) and concatenated with Shannon (dotted lines).

The META assembly in itself yielded higher recall than ClusTrast when ran with any other primary assembly (Trans-ABySS-**M**, Trinity, or RNA-Bloom). But ClusTrast with META as primary assembly yielded higher recall than both META on its own and META with Shannon concatenated.

In Figure 14, we show the number of annotated and expressed isoforms that, of all the methods tested in our study, only ClusTrast managed to reconstruct, and with which tool for primary assembly (RNA-Bloom, Trinity, Trans-ABySS-**M**, and META). ClusTrast with Trans-ABySS-**M** for primary assembly (the default version of ClusTrast) managed to reconstruct most such isoforms in all instances except rice, where it placed second after ClusTrast with META as primary assembly.

**Figure 14:**
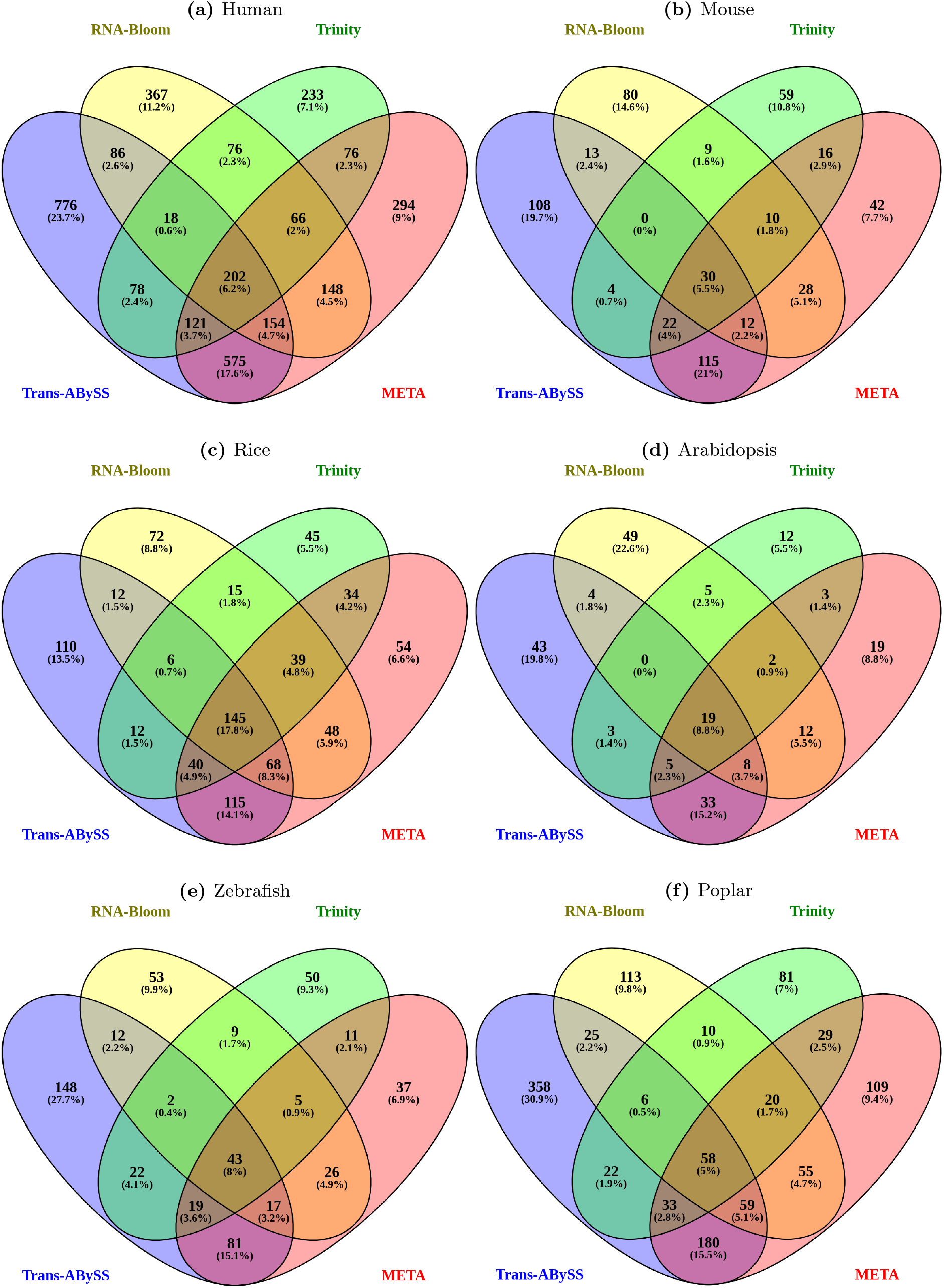
ClusTrast results using four different tools - RNA-Bloom, Trinity, Trans-ABySS-**M**, and META - for the primary assembly. The Venn diagrams show the number of annotated and expressed transcript isoforms reconstructed solely by ClusTrast and no other tested method. ClusTrast with primary assembly from Trans-ABySS-**M** generates the highest number of unique transcript isoforms for all datasets (**a**) - (**f**).

The runtimes for ClusTrast with Trinity as primary assembly were comparable to using Trans-ABySS-**M** for primary assembly. With RNA-Bloom (and thus also META) as primary assembly, however, ClusTrast required more computational resources: the clustering step took 68 hours to complete, and required memory of over 4TB.

Thus, all taken together, we believe that Trans-ABySS-**M** is a reasonable choice for primary assembly in ClusTrast.

## 4 Discussion

We described and assessed the *de novo* transcriptome assembler ClusTrast. Across all six tested datasets, ClusTrast-**M** and ClusTrast-**S** created assemblies that had the highest or among the highest recall as measured by SQANTI and CRBB.

ClusTrast consists of several steps (Figure 1). In the primary assembly step, we have chosen Trans-ABySS as the primary assembler included in the distributed version of ClusTrast. In section 3.3 we show that it is possible to use other assemblers as well, but with lower performance. Using the union of assembled transcripts from more than one assembly tool as the primary assembly improves the recall of ClusTrast. This might be suitable for some users, but requires larger computational resources. The clustering of the guiding contigs is a way to conceptually focus the assembly to genes, and their corresponding isoforms. The most common case is that a reference gene is represented in (i.e., is aligned to) only one cluster, although it is quite common that a reference gene, and its corresponding isoforms, is represented in more than one cluster (Figure S.19). In fact, ClusTrast was used in our recent study (Akhter et al., 2022) to detect isoforms in individual gene families of specific biological interest. The core step of ClusTrast is the clusterwise assembly, which is followed by concatenation of the clusterwise assemblies, and finally by merging with the primary assembly (and removal of duplicates). A comparison of the recall and precision using (i) only the primary assembly, (ii) only the concatenated clusterwise assemblies, and (iii) the final, merged assembly is in Supplementary Tables S.28-S.31. The recall increases substantially using the merged assembly as opposed to only primary or clusterwise assemblies. The fact that the recall increases also when using a union of assembled transcripts from more than one assembly tool as primary assembly (discussed above) further supports that the clusterwise assembly is a helpful approach. We suggest that a wise choice of method for primary assembly coupled with the power of the clusterwise assembly are the key components of the high recall performance of ClusTrast.

In our evaluation of ClusTrast and the other transcriptome assemblers, we have emphasized metrics that do not penalize the reconstruction of different isoforms of a gene. A compact transcriptome assembly is, for instance, preferable when the reconstructed transcripts are intended to be used for aligning reads to a “reference” for, e.g., differential gene expression analysis. In such a situation, our approach would not be helpful. However, when the goal is to find as many supported transcript isoforms as possible, compactness is not in itself desirable and could in fact be counter productive. In a recent review by Thind et al. (2021), the authors point out that there is a need for metrics that better capture the performance with regards to transcript isoforms. Our use of SQANTI’s approach to evaluation is an attempt to address this. SQANTI (Tardaguila et al., 2018) was originally designed to evaluate long reads (e.g., PacBio CCS) but the categories it defined and its aim to classify each non-redundant transcript individually are useful also in evaluation of assembled transcripts. We have been conservative in that we included only the FSM and ISM categories as true positives. Also the NIC category, which encompasses reconstructed transcripts that contain annotated reference exons only, but in novel combinations, could have been included. We compared the results from SQANTI and CRBB (which has been used for transcriptome assembly benchmarking before), and detected a correlation between SQANTI and CRBB scores, particularly strong for recall. We noted that most of the contigs where SQANTI and CRBB classifications of true positives disagreed, belong to the SQANTI class ISM mono-exon. Excluding these from the set of true positives increased the correlation between SQANTI and CRBB precision measurements for ClusTrast (Section 3.1.3). We believe this supports the notion that SQANTI is possible to use for transcriptome assembly evaluation.

Comparing ClusTrast-**M** to one of the most popular transcriptome assemblers, Trinity, revealed that ClusTrast-**M** detected more transcript isoforms than Trinity, and also had a higher precision for isoforms as measured by SQANTI, but clearly underperformed according to CRBB precision. The difference in relative performance of ClusTrast and Trinity according to CRBB and SQANTI precision may be explained by how CRBB handles assembled transcripts with high similarity: If two or more highly similar transcripts exist in the assembly, and some of them have a lower E-value than others, then only the transcripts with E-values below the limit will be considered CRBB hits and thus true positives for precision. SQANTI, in contrast, annotates each transcript independently and therefore calls all assembled transcripts that are similar to a reference transcript as a true positive. We assessed this by recalculating SQANTI precision while only counting one transcript match for every reference transcript (Supplementary Figure S.6), and we observed a marked reduction in precision of Trans-ABySS-**M** and ClusTrast across all assemblies (compare Supplementary Figures S.2 and S.6). We also tested ClusTrast with secondary alignments switched off, and observed a slight improvement for CRBB precision, but at the cost of a reduction in both CRBB and SQANTI recall for most datasets (Supplementary Table S.27).

We observed that ClusTrast generally recovered as many or more known isoforms as TrAB+Sh as measured by SQANTI (Supplementary Figure S.2) while suffering only a small reduction in CRBB precision (Supplementary Figure S.3) and that ClusTrast finished successfully on the rice dataset, where Shannon (and thus TrAB+Sh) failed to create an assembly. A possible explanation for both observations is that the clustering performed in ClusTrast may simplify sub-graphs enough to allow better handling by the Shannon heuristic (Section 2.1.4) and thereby increase sensitivity. ClusTrast-**M** and TrAB**-M**+Sh were also the best in reconstructing isoforms to their full length according to SQANTI (Section 3.1.4).

In general the performance of the assemblers was rather consistent over species, regardless of evaluation approach (SQANTI or CRBB). We tested two additional human datasets (Supplementary Table S.1), for a total of four datasets, one of which simulated, and observed that ClusTrast showed the highest SQANTI and CRBB recall over the range of transcript coverage as well as expression levels for all four human datasets (Supplementary Figures S.7–S.9). However, the precision performance of ClusTrast was mixed. ClusTrast performance was not correlated to the size of the datasets (Supplementary Figure S.17). For all results on all additional datasets, see Supplementary Section C.3.

We used RSEM to estimate the number of expressed isoforms, and in our recall calculations we used only transcripts with TPM>0. Using all reference transcripts in our evaluation, instead of only those that have a TPM>0, would mean a larger denominator when calculating recall, which would lower the recall of all compared assemblers alike. If RSEM makes a mistake and assigns TPM=0 to a reference transcript that in fact is transcribed, then the recall will be underestimated. If RSEM assigns a TPM>0 to a reference transcript which in fact is not transcribed, recall will be overestimated. Similarly, there might be assembled contigs that correspond to real isoforms that are not present in the reference. These contigs are counted as false negatives while they should be true positives, thus precision performance is likely underestimated. For the SQANTI evaluation, it is possible that many of these would be considered as true positives if we had included the NIC category among the true positives.

Within the scope of this paper, we have shown that the current *de novo* transcriptome assembly strategy of ClusTrast is successful in finding the most comprehensive set of isoforms from short read RNA-seq datasets, as it outperformed all other tested methods. We do not claim, however, that we have found the optimal version. Future improvements might come, e.g., from testing other assemblers for the clusterwise assembly, or, provided suitable datasets are available, from including long reads as guiding contigs. The use of long reads in transcriptome analysis is not a focus of this study, but the accuracy of long reads has indeed reached a level that enables the use of long read sequencing to reliably address questions not easily amenable with short read sequencing. For instance, long reads can provide direct information about transcript isoforms present in a sample or the methylation status of DNA or RNA molecules (Kovaka et al., 2023). The most telling example is perhaps the telomere-to-telomere (T2T) sequencing of the human genome (Nurk et al., 2022): long reads revealed approximately 200 Mbases of genomic DNA sequence hitherto undetermined, containing 99 protein coding genes not present in the human reference genome GRCh38. There are emerging methods for analysis of long read transcriptome data, e.g. StringTie2 which can accommodate both short and long reads for a reference-based transcriptome assembly (Kovaka et al., 2019) and IsoQuant that enables transcript discovery in long read data sets (Prjibelski et al., 2023). Finally, we note that a short read *de novo* transcriptome assembly approach, such as ClusTrast or any of the ones included in our study, would potentially be able to find the likes of the missing 99 human genes in other species which not yet have had their full T2T genomes sequenced. This attests to the lingering usefulness of short read sequencing and also to the advantages of *de novo* transcriptome assembly, as a reference-based method by definition would miss any genes or transcripts not present in the reference.

## 5 Conclusion

In our tests of model organisms, ClusTrast consistently detected the most transcript isoforms, not the least for the isoforms in the lower end of the expression range (Section 3.1.6), but at a cost of lower precision. This agrees with our intention of ClusTrast – to provide a comprehensive but non-redundant list of contigs. Therefore, we believe researchers interested in a more complete representation of transcript isoforms from eukaryotic organisms may wish to use ClusTrast. The resulting list of contigs is amenable for further processing and analysis tailored according to the research question at hand.

## Availability and requirements

**Project name**: ClusTrast

**Project home page**: https://github.com/karljohanw/clustrast

**Operating systems**: Linux and MacOS

**Programming language**: Bashscript

**Requirements**: transabyss, shannon_cpp, isONclust, minimap2, awk.

**License**: GPLv3

**Restrictions to use by non-academics**: None

## Authors’ contributions

KJW implemented the method, ran all the experiments and wrote the original version of the manuscript. KJW and OE wrote the final manuscript, with contributions from WWK. KJW and OE analyzed the results. OE and WWK supervised the project. All authors read and approved the final version of the manuscript.

## Acknowledgements

We wish to thank Pelin Akan Sahlén at KTH Royal Institute of Technology for sharing access to the server SAGA.

