## Supplemental material for "ClusTrast: a short read *de novo* transcript isoform assembler guided by clustered contigs"

v2023-02-17

#### Appendix
## &
#### Supplementary material

Karl Johan Westrin,[1]
Warren W. Kretzschmar,[1,2]
Olof Emanuelsson,[1]

[1] Science for Life Laboratory, Department of Gene Technology, KTH Royal Institute of Technology, SE-171 65, Solna, Sweden
[2] Center for Hematology and Regenerative Medicine (HERM), Department of Medicine Huddinge, Karolinska Institute, SE-141 52, Flemingsberg, Sweden

### A Tools and versions

#### A.1 Pre-processing

- **fastp** 0.20.1 (Chen et al., 2018)

#### A.2 Assembly

- **Trinity** 2.10.0 (Grabherr et al., 2011)
  - **Jellyfish** 2.2.7 (Marçais and Kingsford, 2011)
  - **Bowtie** 2.3.4.1 (Langmead and Salzberg, 2012)
  - **Salmon** 1.0.0 (Patro et al., 2017)
- **BinPacker** 1.1 (Liu et al., 2016)
- **TransLiG** 1.3 (Liu et al., 2019)
- **Trans-ABySS** 2.0.1 (Robertson et al., 2010)
  - **ABySS** 2.1.5 (Simpson et al., 2009; Jackman et al., 2017)
- **Shannon** 0.5.0 (Kannan et al., 2016)
  - **Jellyfish** 2.3.0 (Marçais and Kingsford, 2011)
- **SOAPdenovo-Trans** 1.03 (Xie et al., 2014)
- **Oases** 0.2.09 (Schulz et al., 2012)
  - **Velvet** 1.2.10 (Zerbino and Birney, 2008)
- **rnaSPAdes** 3.14.1 (Bushmanova et al., 2019; Bankevich et al., 2012)
- **RNA-Bloom** 1.3.1 (Nip et al., 2020)
  - **minimap2** 2.17 (Li, 2018)
  - **ntCard** 1.2.1 (Mohamadi et al., 2017)
- **ClusTrast** 0.0.0
  - **isONclust** 0.0.6 (Sahlin and Medvedev, 2020)
  - **minimap2** 2.17 (Li, 2018)

#### A.3 Quality

- **RSEM** 1.3.3 (Li and Dewey, 2011)
  - **HISAT2** 2.2.1 (Kim et al., 2019)
- **SQANTI3** 1.4.9 (Tardaguila et al., 2018)
  - **minimap2** 2.17 (Li, 2018)
- **TransRate** 1.0.1 (Smith-Unna et al., 2016)
  - **BLAST+** 2.2.29 (Camacho et al., 2009)

### B Commands

#### B.1 Pre-processing

##### B.1.1 fastp

```
fastp --in1 ${SAMPLE}_1.fastq --in2 ${SAMPLE}_2.fastq \
  --out1 ${SAMPLE}_fastp_1.fq.gz --out2 ${SAMPLE}_fastp_2.fq.gz --thread ${CPU}
```

#### B.2 Assembly

##### B.2.1 Trans-ABySS

```
transabyss --pe ${INPUT_FILE_1} ${INPUT_FILE_2} --threads ${CPU} --outdir ${OUTPUT_DIR}
```

##### B.2.2 Trans-ABySS – multi-$k$

```
for K in `seq 25 7 53`; do
  transabyss -k ${K} --pe ${INPUT_FILE_1} ${INPUT_FILE_2} \
    --threads ${CPU} --outdir ${OUTPUT_DIR}/${K}
done
transabyss-merge --mink 25 --maxk 53 ${OUTPUT_DIR}/*/transabyss-final.fa \
  --threads ${CPU} --abyssmap --out ${OUTPUT_DIR}/${DATASET}.fa
```

##### B.2.3 Trinity

```
Trinity --seqType fq --left ${INPUT_FILE_1} --right ${INPUT_FILE_1} \
  --max_memory ${RAM}G --CPU ${CPU} --output ${OUTPUT_DIR}
```

##### B.2.4 BinPacker

```
BinPacker -s fq -p pair -l ${INPUT_FILE_1} -r ${INPUT_FILE_2} -o ${OUTPUT_DIR}
```

##### B.2.5 TransLiG

```
TransLiG -s fq -p pair -l ${INPUT_FILE_1} -r ${INPUT_FILE_2} -o ${OUTPUT_DIR}
```

##### B.2.6 Oases

```
velveth ${OUTPUT} 19,33,2 -shortPaired -fastq.gz -separate ${INPUT_FILE_1} ${INPUT_FILE_2}
for K in `seq 19 2 31`; do velvetg ${OUTPUT}_${K} -read_trkg yes ; done
for K in `seq 31 -2 19`; do oases ${OUTPUT}_${K} -ins_length 500 & ; done
wait
velveth ${OUTPUT}Merged 27 -long ${OUTPUT}_[1-3][0-9]/transcripts.fa
velvetg ${OUTPUT}Merged -conserveLong yes -read_trkg yes
oases ${OUTPUT}Merged -merge yes
```

##### B.2.7 Shannon

```
RDLN1=`zcat ${INPUT_FILE_1} | awk '(NR==2){ print length($0); exit 0}'`
RDLN2=`zcat ${INPUT_FILE_2} | awk '(NR==2){ print length($0); exit 0}'`
shannon_cpp shannon --output_dir ${OUTPUT_DIR} --pair_read_length ${RDLN1} ${RDLN2} \
  --PE_read_path ${INPUT_FILE_1} ${INPUT_FILE_2} --num_process=${CPU} \
  --avail_mem=${RAM}G --bypass_pre_correct_read
```

##### B.2.8 RNA-Bloom

```
rnabloom -left ${INPUT_FILE_1} -right ${INPUT_FILE_2} \
  -revcomp-right -ntcard -t ${CPU} -outdir ${OUTPUT_DIR}
```

##### B.2.9 rnaSPAdes

```
rnaspades.py -o ${OUTPUT_DIR} -1 ${INPUT_FILE_1} -2 ${INPUT_FILE_2} -t ${CPU}
```

##### B.2.10 SOAPdenovo-Trans

```
RDLEN=`zcat ${INPUT_FILE_1} | awk '(NR==2){ print length($0); exit 0}'`
cat <<EOF > soap.config
> max_rd_len=${RDLEN}
> [LIB]
> avg_ins=200
> q1=${INPUT_FILE_1}
> q2=${INPUT_FILE_2}
> EOF
SOAPdenovo-Trans-31mer all -s soap.config -p ${CPU} -L 200 -o ${OUTPUT_DIR}
```

##### B.2.11 ClusTrast

```
clustrast -u -1 ${INPUT_FILE_1} -2 ${INPUT_FILE_2} -p ${CPU} -m ${MEM_LIM} \
  -t /tmp/clustrast_${DATASET} -o ${OUTPUT}
```

#### B.3 Quality

##### B.3.1 RSEM

```
rsem-prepare-reference --hisat2-hca --num-threads ${CPU} \
  --gtf ${REF_ANNOT} ${REF_GEOME} ${SPECIES}_ref
rsem-calculate-expression --paired-end <(zcat ${INPUT_FILE_1}) <(zcat ${INPUT_FILE_2}) \
  --num-threads ${CPU} --hisat2-hca ${SPECIES}_ref ${SAMPLE}_expr
```

##### B.3.2 SQANTI3

```
sqanti3_qc ${ASSEMBLY} ${REF_ANNOTATION} ${REF_GENOME} --cpus ${CPU} \
  --output ${ASSEMBLER}_${SAMPLE}_sqanti3 --force_id_ignore --skipORF --skip_report

# generate reverse complements for antisense classifications
seqtk subseq ${ASSEMBLY} <(awk '($6=="antisense"){sub(/_dup[0-9]*$/,"",$1); \
  if (!_[$1]++){ print $1 };}' ${SQANTI_FILE}) | seqtk seq -Ar - > antisense.fa
sqanti3_qc antisense.fa ${REF_ANNOTATION} ${REF_GENOME} --cpus ${CPU} \
  --output ${ASSEMBLER}_${SAMPLE}_AS_sqanti3 --force_id_ignore --skipORF --skip_report
rm antisense.fa AS_contigs.txt
```

##### B.3.3 TransRate

```
# the ref-based score from TransRate is actually calculated by CRBB.
transrate --threads=${CPU} --output=${ASSEMBLER}_${SAMPLE} \
  --assembly=${ASSEMBLY} --reference=${REF_TRANSCR}
```

#### C Tables and Results

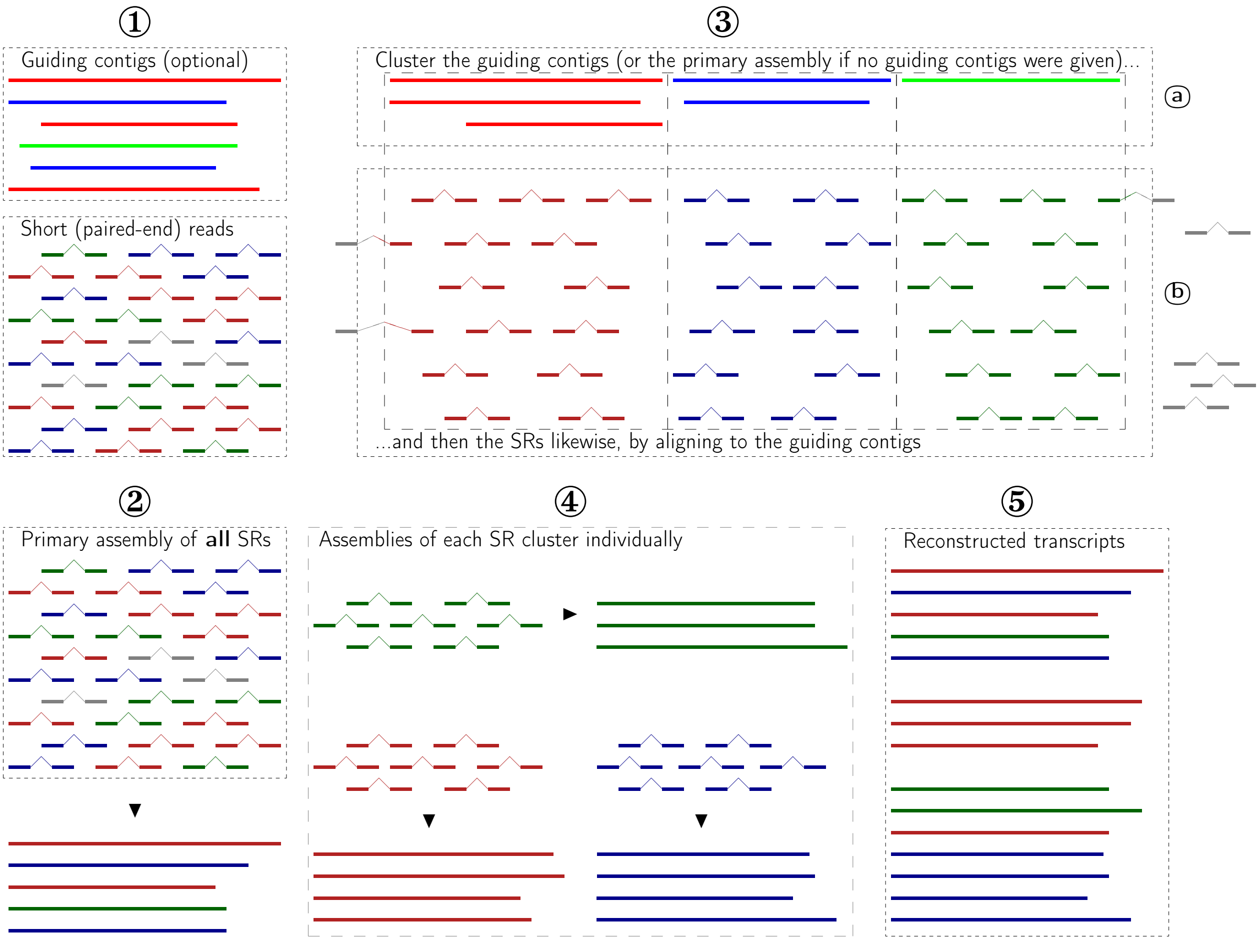


**Figure S.1:** An illustration of how ClusTrast works: Given (1) a set of short RNA-seq reads (SRs) and, optionally, a set of guiding contigs (GCs), (2) assemble the SRs to a primary assembly and (3a) cluster the GCs. If no GCs are given, the primary assembly will be used as GCs. (3b) Align the SRs to the GCs, and assign (based on the alignment) the SRs to the GC clusters (i.e., SRs are now also clustered). (4) Assemble each SR cluster individually. (5) Merge all cluster assemblies and the primary assembly and, by default, remove identical transcripts.

**Table S.1:** Short read RNA-seq datasets used in supplementary. For datasets (also) used in the main paper, see Table 1. RL=read length in bases. SS=strand specificity (assembled with such flags when possible). RPs=million read pairs, before pre-processing (on the left) and after pre-processing (on the right).

| SRA ID | RL | SS | Species | *Notes* | RPs | |
|---|---|---|---|---|---|---|
| SRR1153470 | 2 × 101 | RF | **Human** | Tilgner et al. (2014) | 115 | 103 |
| SRR5344669.1 | 2 × 130 | RF | **Arabidopsis** | Zhang et al. (2017) | 61 | 59 |
| SRR10853135 | 2 × 150 | RF | **Poplar** | inoculated w. *S. musiva* | 31.9 | 31.7 |
| SRR8594084 | 2 × 76 | n/a | **Human** | | 12.4 | 12.3 |
| n/a *(simulated)* | 2 × 100 | n/a | **Human** | Hölzer and Marz (2019) | 57 | n/a |
| n/a *(simulated)* | 2 × 100 | n/a | **Mouse** | Hayer et al. (2015) | 50 | n/a |

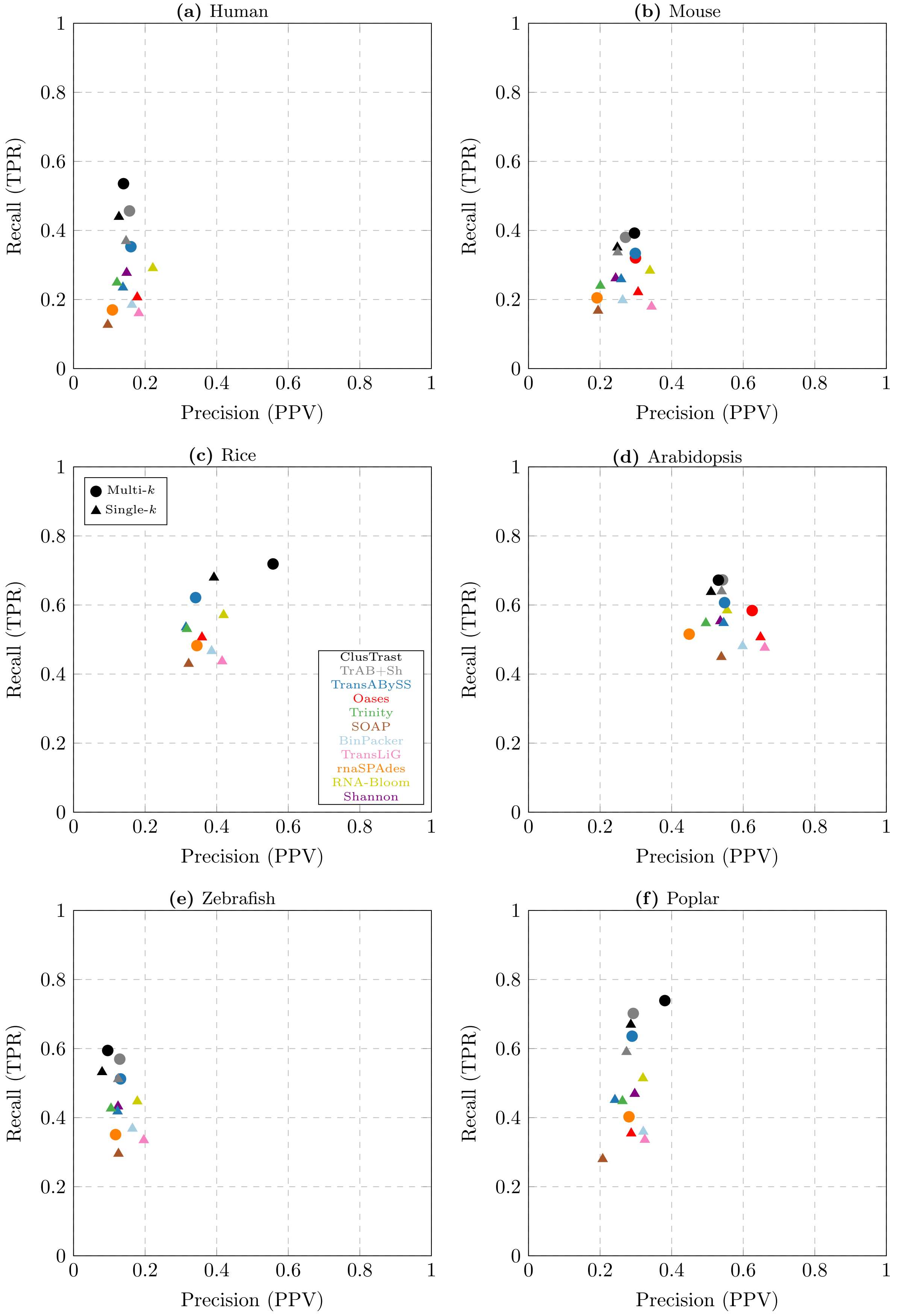


**Figure S.2:** SQANTI precision versus recall. Each FSM and ISM with at least half of the reference isoform's exons is counted as a true positive for precision. Each reference isoform with at least half of its exons covered by a FSM or ISM is counted as a true positive for recall.

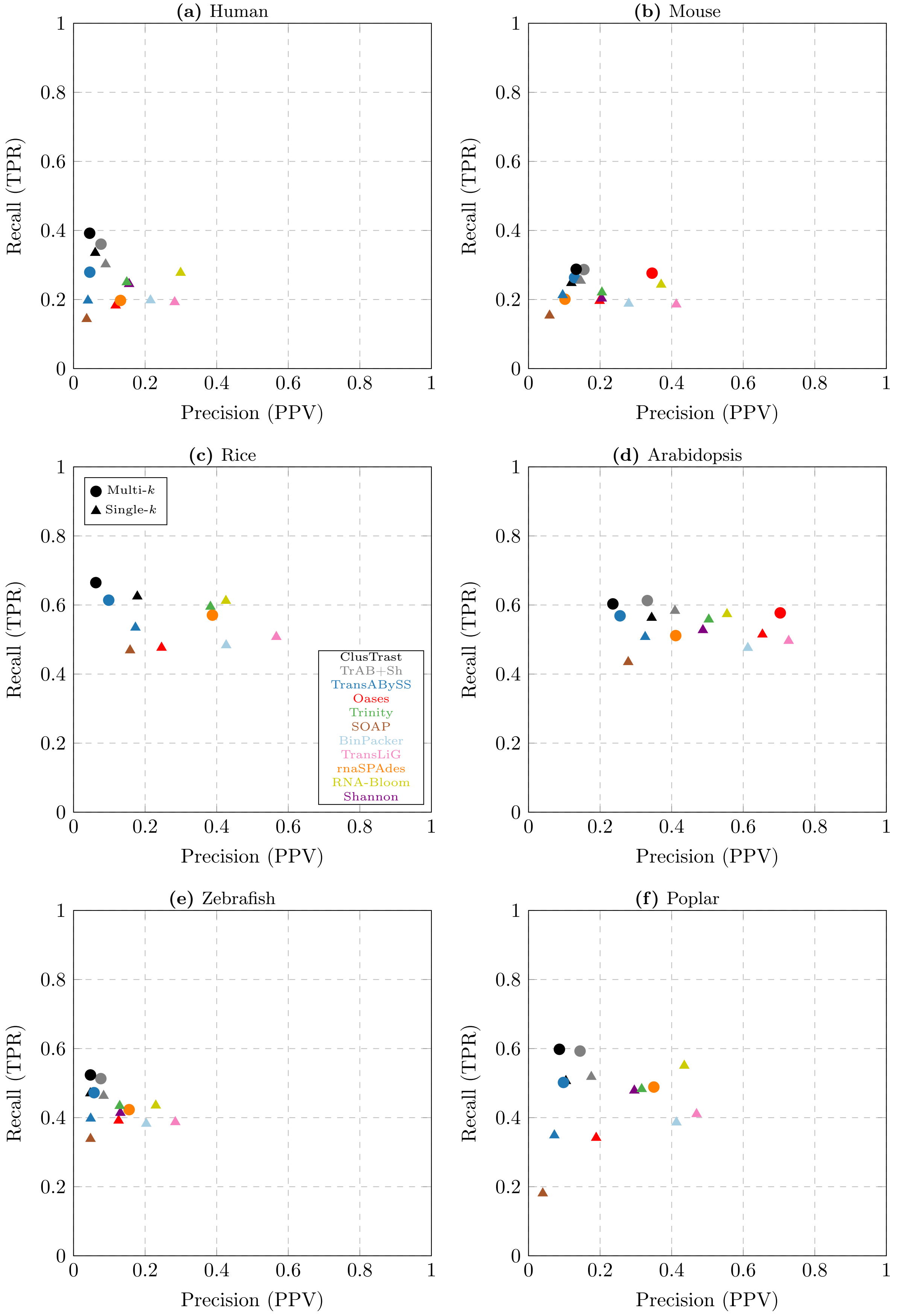


**Figure S.3:** CRBB Precision vs. Recall. Precision is the proportion of assembled contigs that are CRBB hits and cover at least 50% of a reference isoform. Recall is the proportion of reference isoforms covered to at least 50% by an assembled contig that is a CRBB hit.

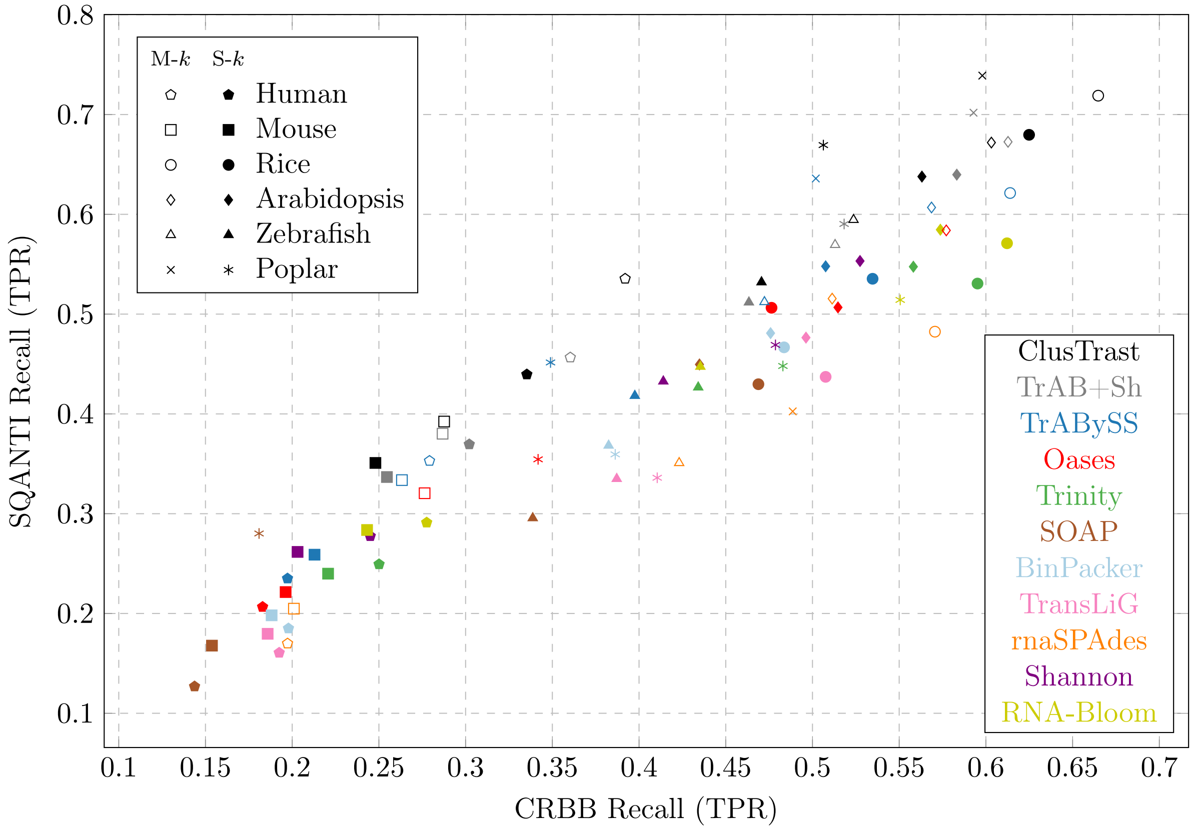


**Figure S.4:** Recall (TPR) – SQANTI vs. CRBB. $\rho = 0.93398049$

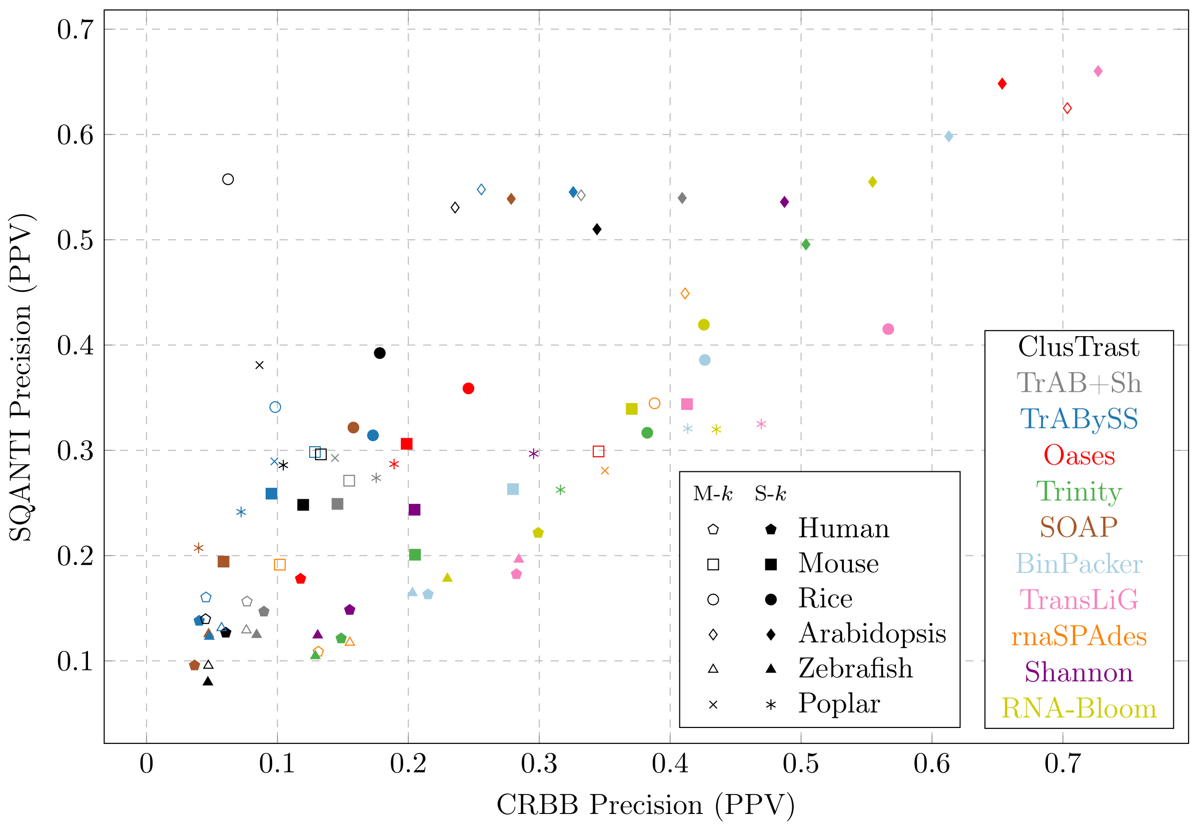


**Figure S.5:** Precision (PPV) – SQANTI vs. CRBB. $\rho = 0.74571316$

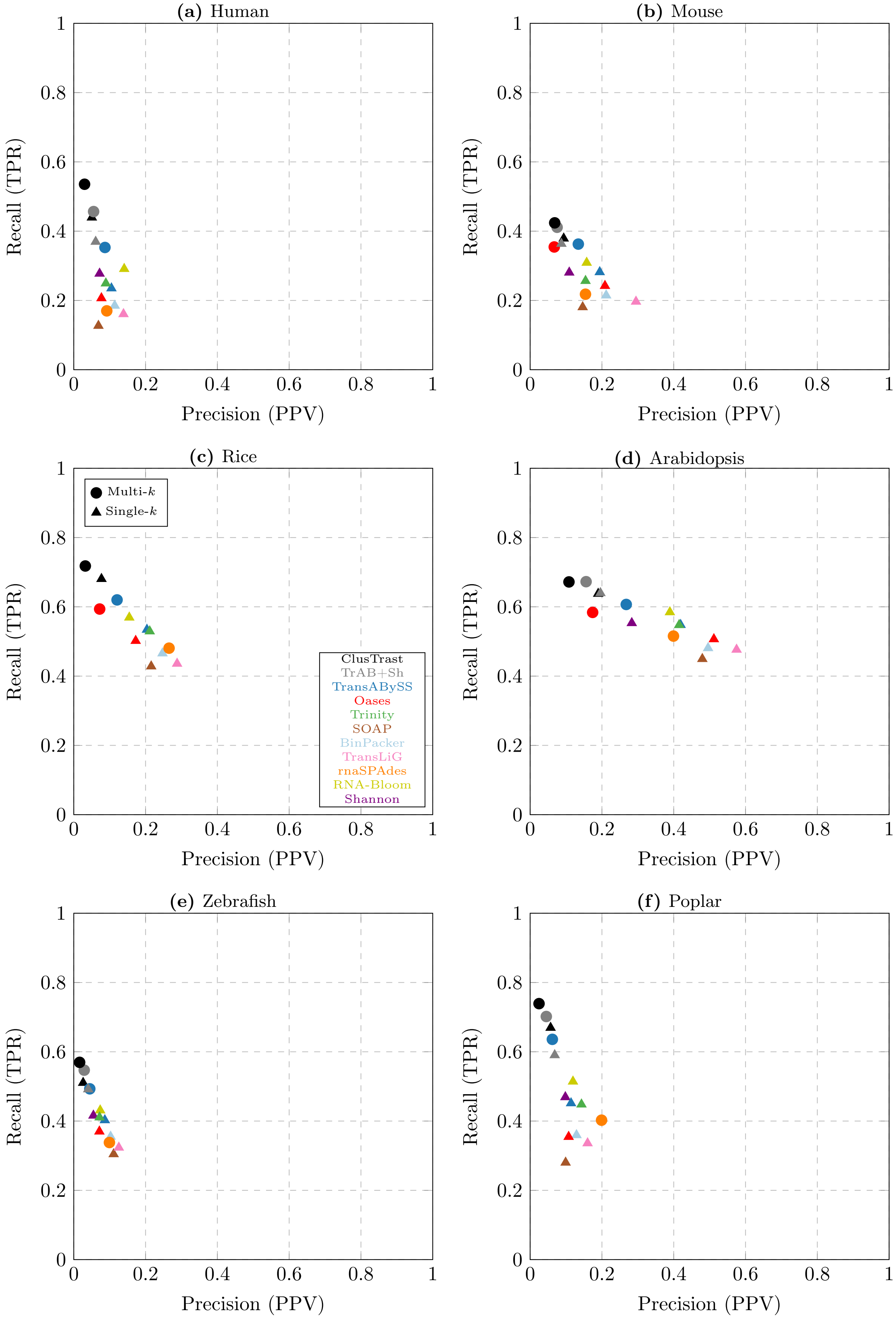


**Figure S.6:** SQANTI precision versus recall. Each *first* FSM or ISM with at least half of a reference isoform's exons is counted as a true positive for precision. Each reference isoform with at least half of its exons covered by a FSM or ISM is counted as a true positive for recall.

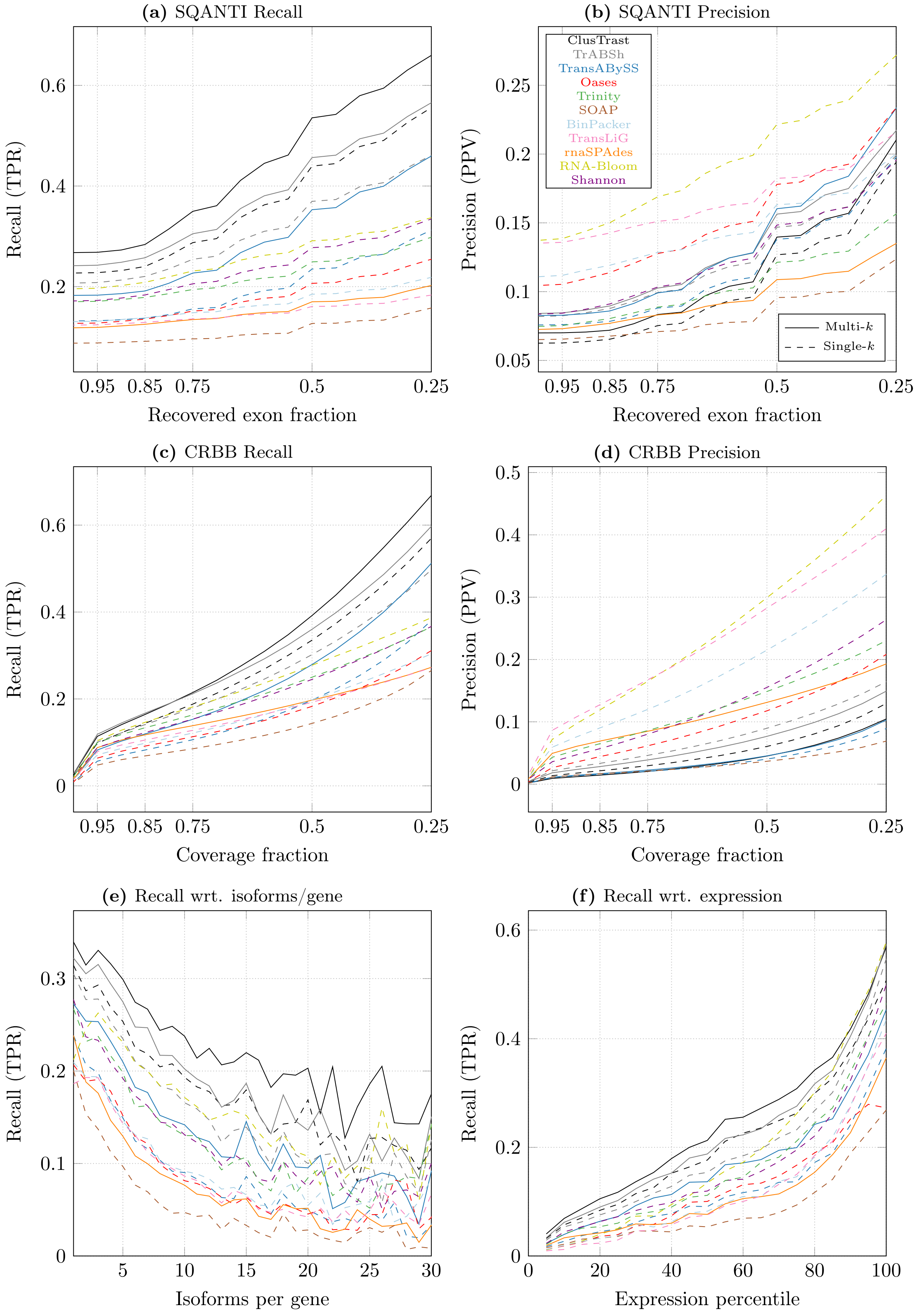


**Figure S.7:** Results for the human dataset SRR5133163.1.

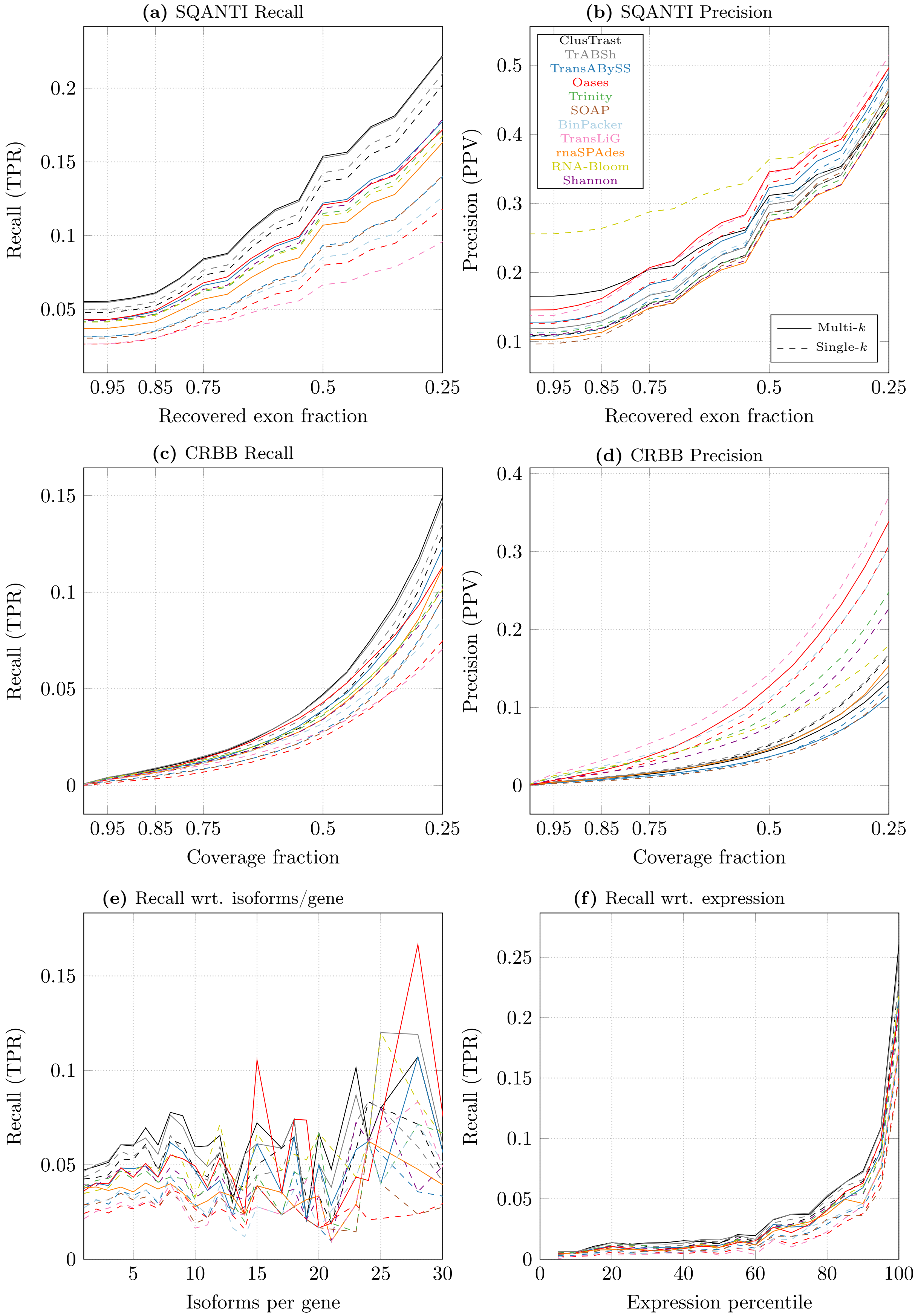


**Figure S.8:** Results for the extra human dataset SRR8594084.

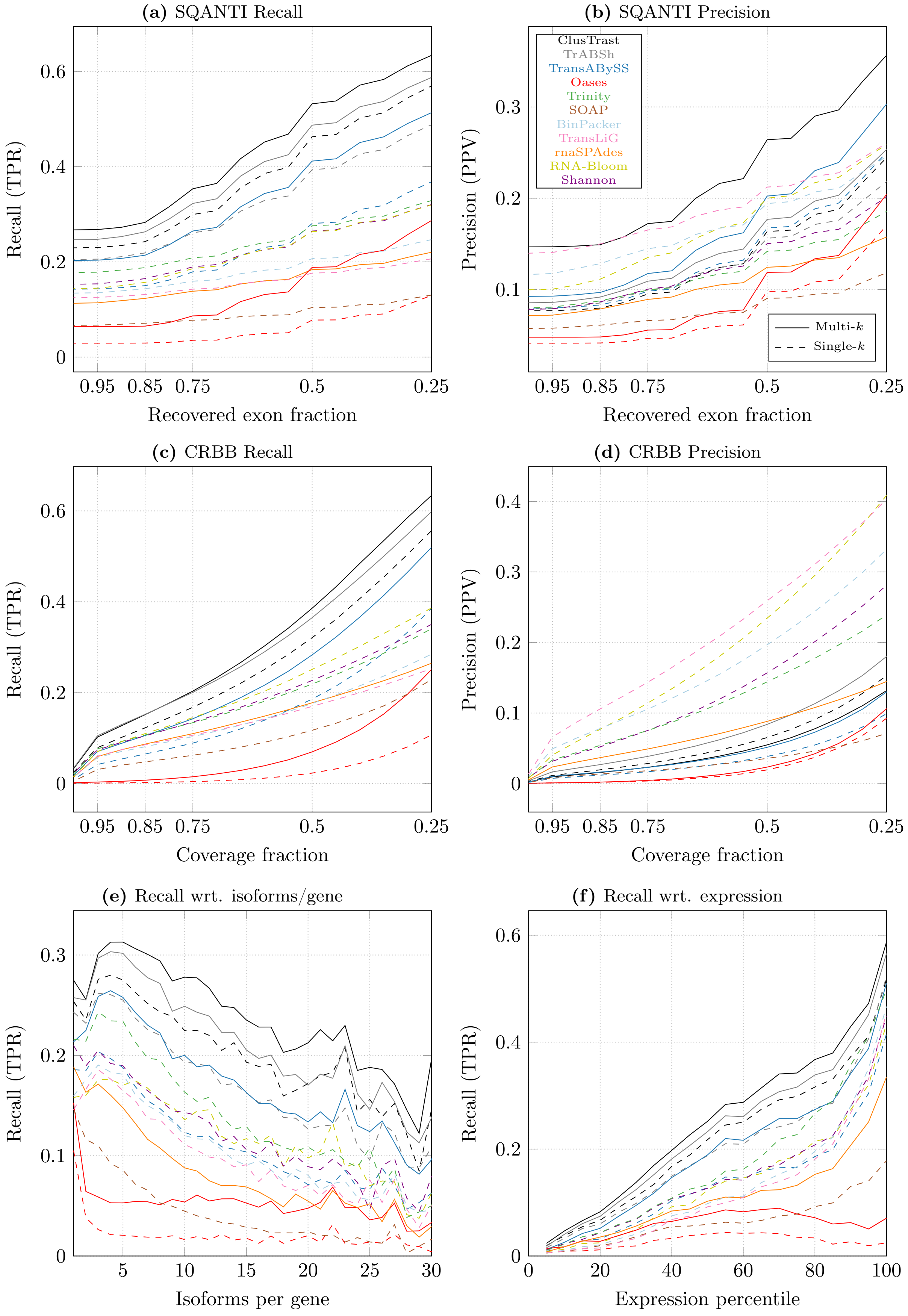


**Figure S.9:** Results for the stranded human dataset SRR1153470.

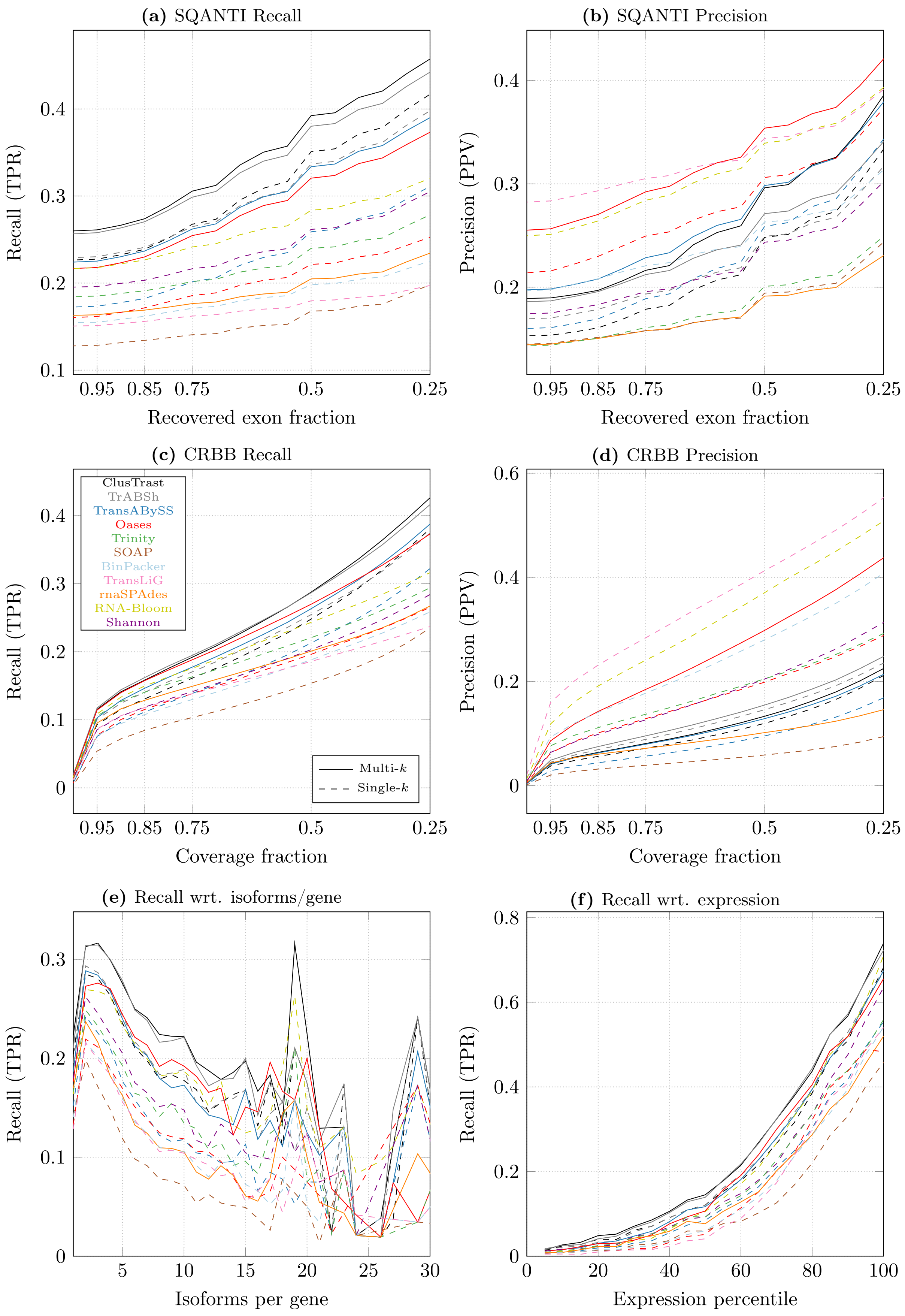


**Figure S.10:** Results for the mouse dataset SRR8632985.

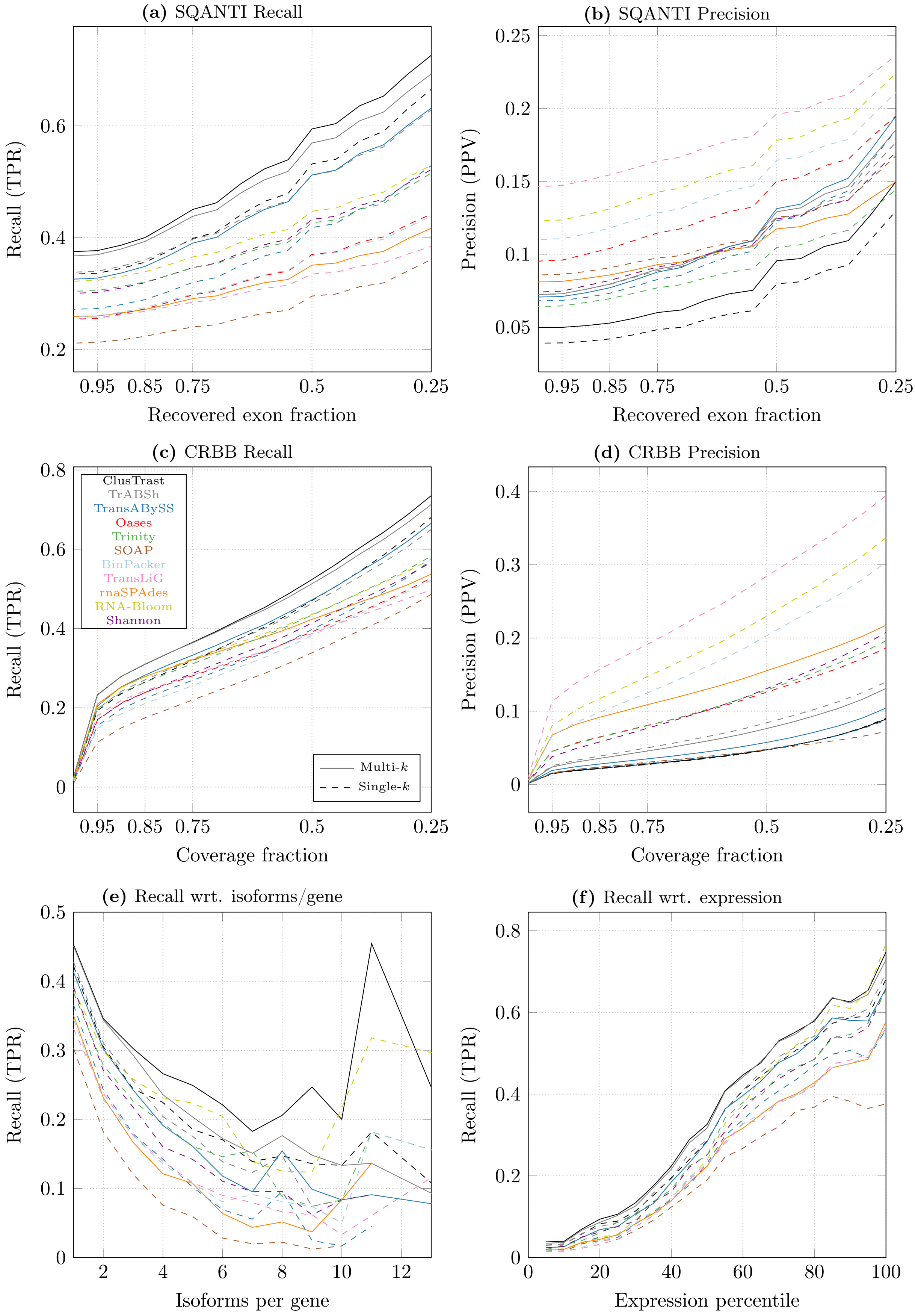


**Figure S.11:** Results for the zebrafish dataset SRR10728575.

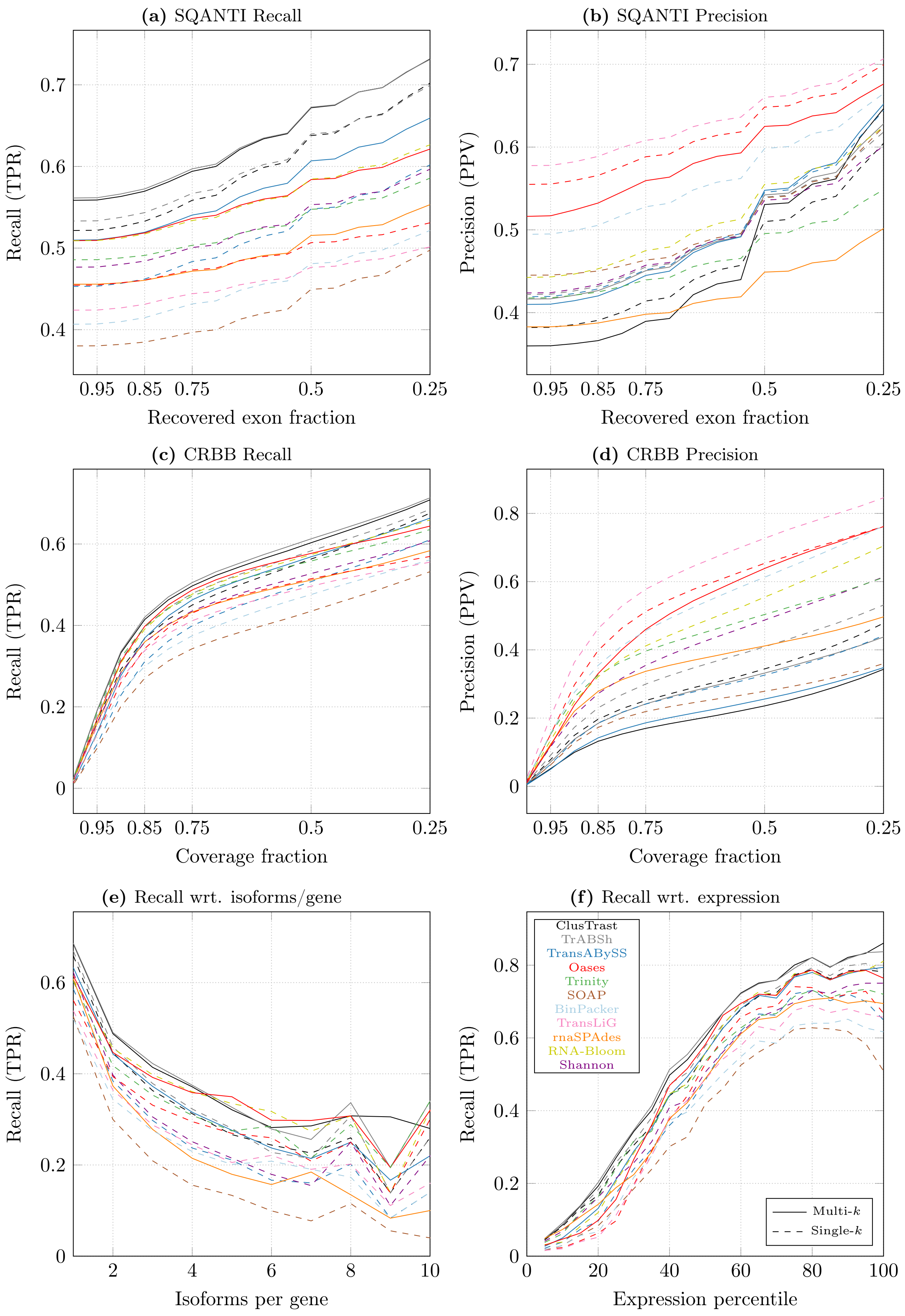


**Figure S.12:** Results for the arabidopsis dataset SRR11278019.

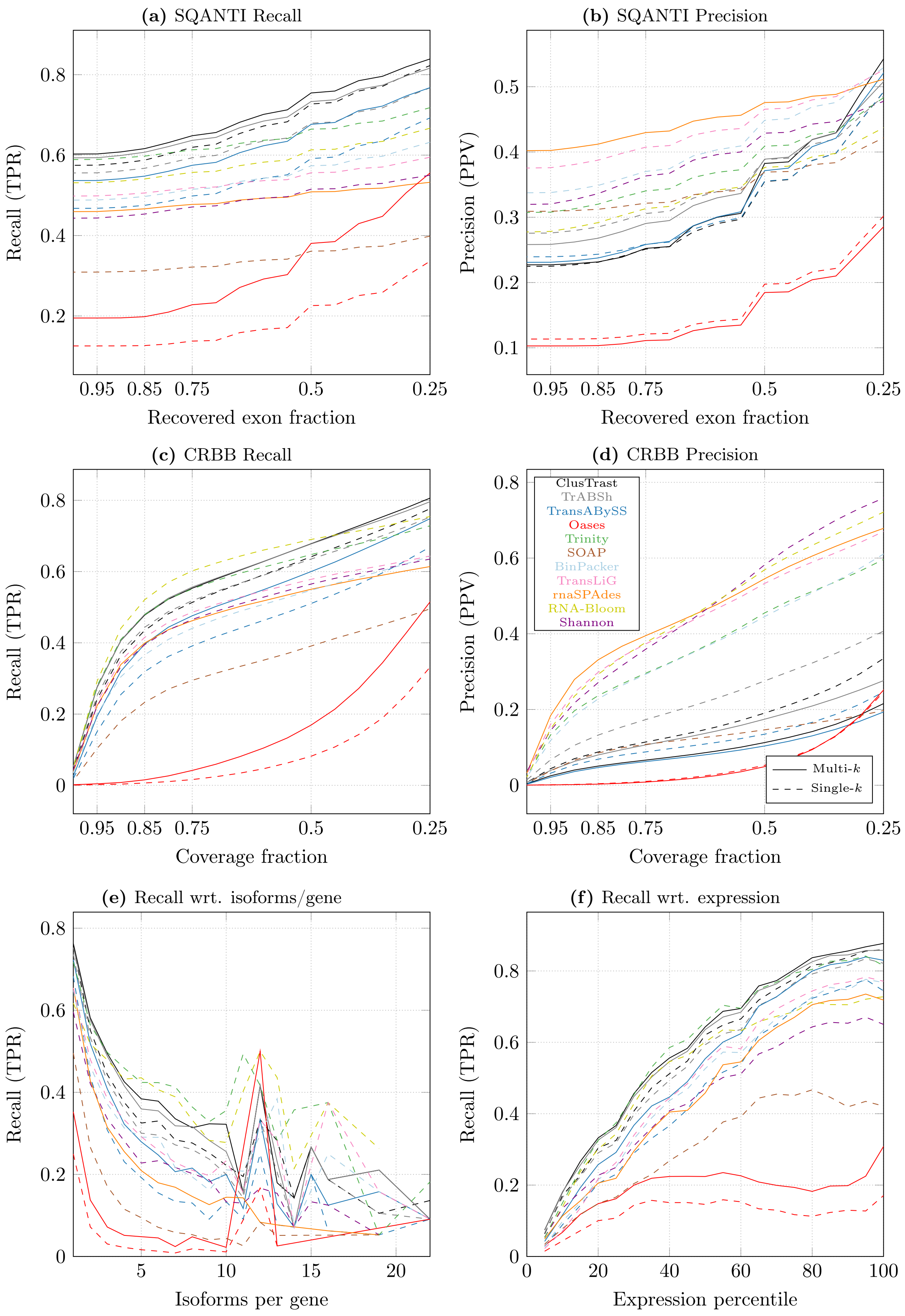


**Figure S.13:** Results for the stranded arabidopsis dataset SRR5344669.1.

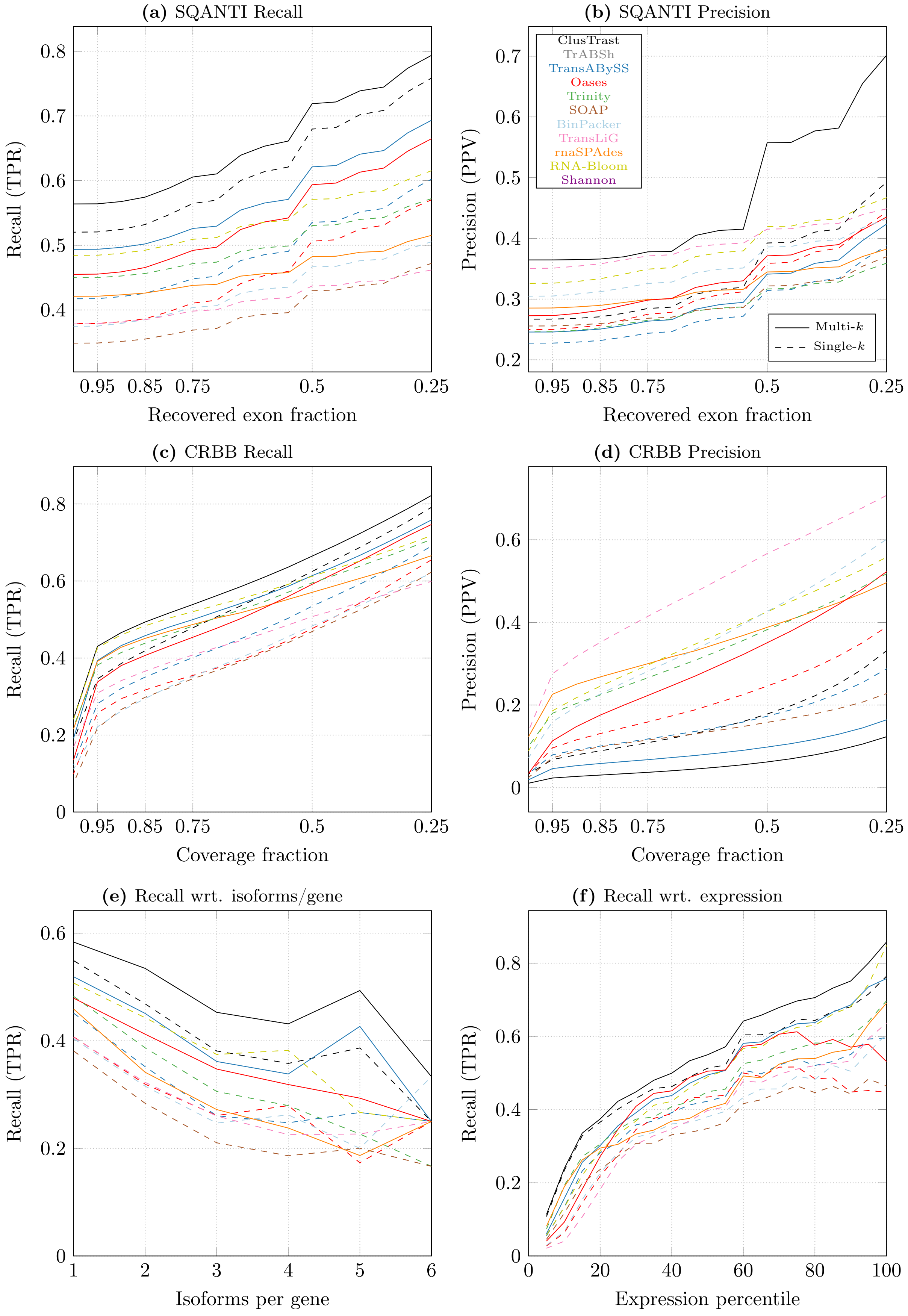


**Figure S.14:** Results for the rice dataset SRR11341576.

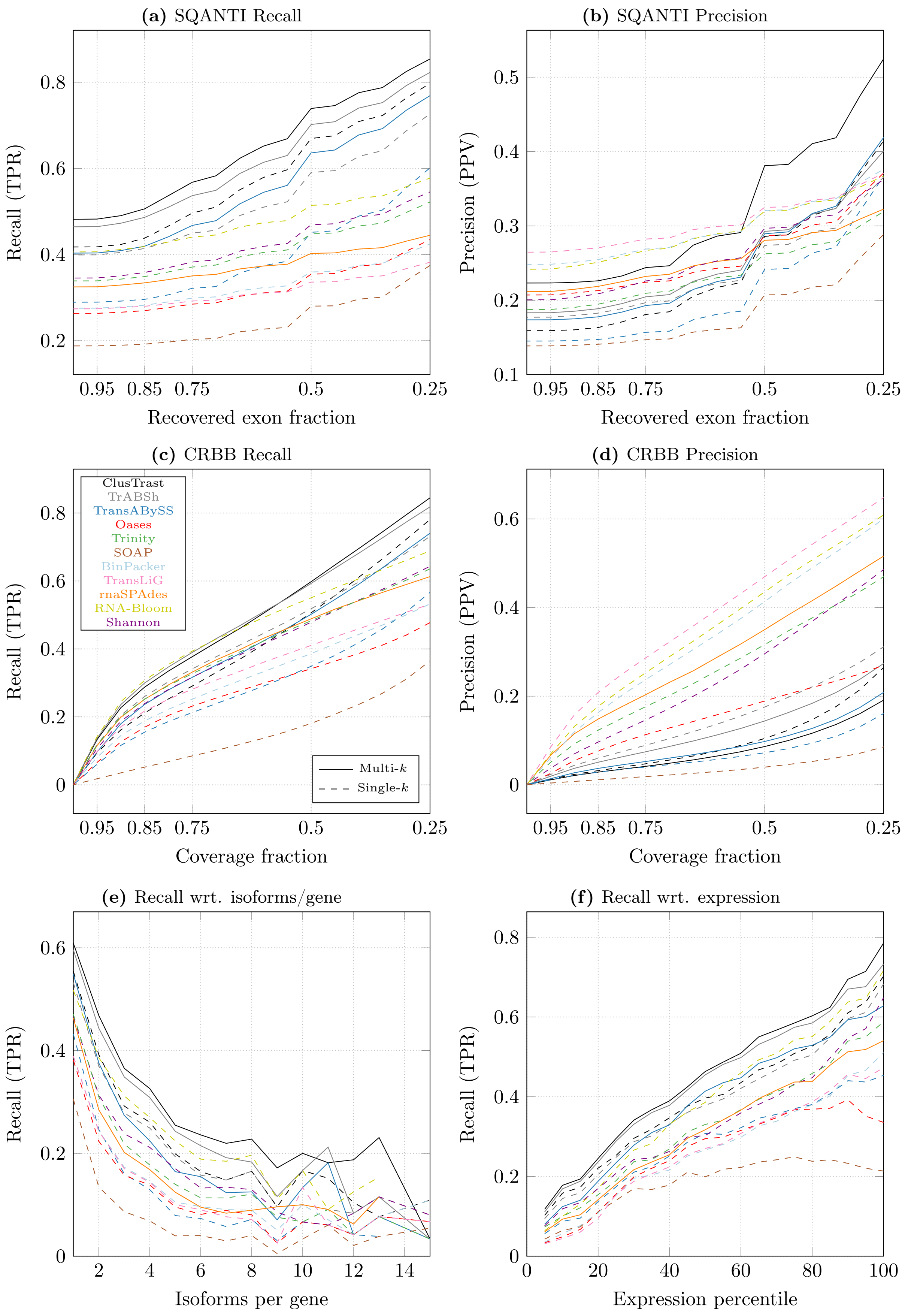


**Figure S.15:** Results for the poplar dataset SRR5986240.1.

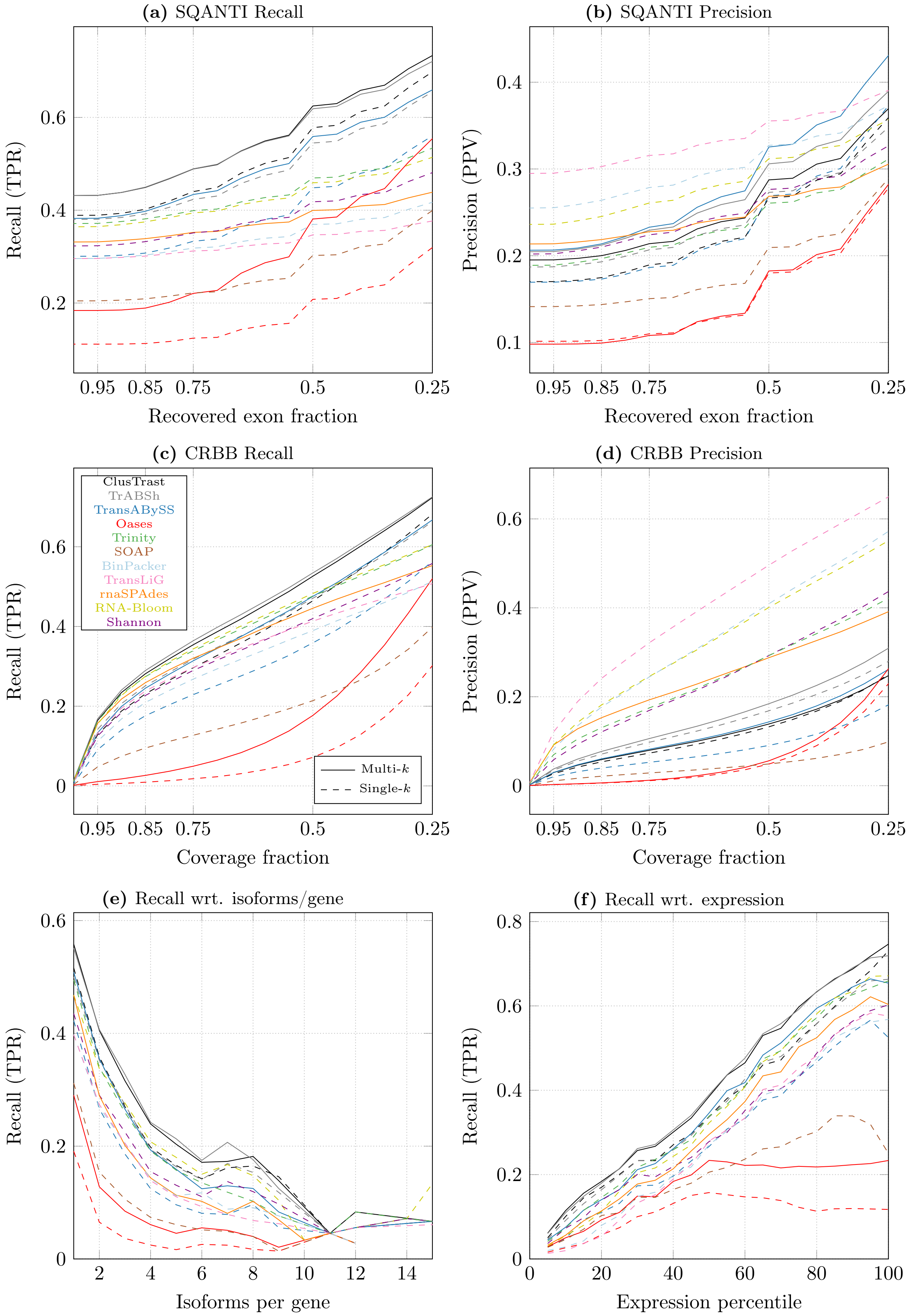


**Figure S.16:** Results for the stranded poplar (metatranscriptomic) dataset SRR10853135.

**Table S.2:** Contig metrics and statistics for the human dataset SRR5133163.1.

| *Contig stats* | Lengths | | Contigs ≥ 0bp | | | Contigs ≥ 500bp | | |
|---|---|---|---|---|---|---|---|---|
| Human | Min | Max | \|seqs\| | Mean | Median | \|seqs\| | Mean | Median |
| ClusTrast-M | 100 | 26345 | 2163519 | 435.95 | 253 | 408592 | 1282.29 | 824 |
| ClusTrast | 100 | 22107 | 1064694 | 527.65 | 282 | 276540 | 1325.19 | 857 |
| TrABySS-M | 100 | 26345 | 898260 | 414.14 | 150 | 161550 | 1500.00 | 975 |
| TrABySS | 100 | 21754 | 473524 | 388.15 | 150 | 78801 | 1372.13 | 899 |
| Shannon | 201 | 34864 | 348526 | 1203.10 | 492 | 172464 | 2117.95 | 1375 |
| Oases-31 | 81 | 31430 | 356142 | 857.39 | 310 | 130944 | 1947.05 | 1250 |
| Trinity | 184 | 28231 | 250254 | 1032.44 | 418 | 106466 | 2013.49 | 1226 |
| SOAP | 100 | 25410 | 380509 | 407.79 | 150 | 53099 | 1782.18 | 1003 |
| BinPacker | 200 | 27157 | 143970 | 1500.42 | 683 | 92340 | 2159.68 | 1294 |
| TransLiG | 200 | 49419 | 101000 | 2094.36 | 958 | 79587 | 2569.26 | 1433 |
| rnaSPAdes | 73 | 35733 | 166659 | 1046.41 | 421 | 69336 | 2059.59 | 1156 |
| RNA-Bloom | 218 | 25109 | 183761 | 1636.42 | 889 | 130272 | 2155.04 | 1465 |

**Table S.3:** Contig metrics and statistics for the mouse dataset SRR8632985.

| *Contig stats* | Lengths | | Contigs ≥ 0bp | | | Contigs ≥ 500bp | | |
|---|---|---|---|---|---|---|---|---|
| Mouse | Min | Max | \|seqs\| | Mean | Median | \|seqs\| | Mean | Median |
| ClusTrast-M | 100 | 19508 | 480816 | 667.62 | 280 | 145852 | 1673.32 | 1192 |
| ClusTrast | 100 | 19508 | 311823 | 631.01 | 273 | 90510 | 1611.45 | 1145 |
| TrABySS-M | 100 | 14372 | 278764 | 614.03 | 204 | 77504 | 1718.83 | 1268 |
| TrABySS | 100 | 14372 | 167676 | 510.76 | 184 | 38143 | 1611.95 | 1161 |
| Shannon | 201 | 15524 | 144107 | 1082.99 | 466 | 69121 | 1938.45 | 1444 |
| Oases | 100 | 46184 | 347823 | 1263.96 | 718 | 206121 | 1963.77 | 1517 |
| Trinity | 185 | 17458 | 94085 | 1032.58 | 404 | 40572 | 2008.12 | 1471 |
| SOAP | 100 | 14062 | 162120 | 403.01 | 164 | 22687 | 1780.21 | 1229 |
| BinPacker | 201 | 13889 | 56051 | 1373.49 | 723 | 35955 | 1961.96 | 1410 |
| TransLiG | 200 | 32572 | 35752 | 1920.33 | 1233 | 30054 | 2221.41 | 1590 |
| rnaSPAdes | 37 | 17841 | 135571 | 621.16 | 220 | 32150 | 1920.00 | 1391 |
| RNA-Bloom | 200 | 14188 | 108737 | 1463.97 | 872 | 70342 | 2093.94 | 1640 |

**Table S.4:** Contig metrics and statistics for the rice dataset SRR11341576.

| *Contig stats* | Lengths | | Contigs ≥ 0bp | | | Contigs ≥ 500bp | | |
|---|---|---|---|---|---|---|---|---|
| Rice | Min | Max | \|seqs\| | Mean | Median | \|seqs\| | Mean | Median |
| ClusTrast-M | 100 | 29078 | 1122834 | 463.15 | 293 | 272436 | 1146.31 | 792 |
| ClusTrast | 100 | 29078 | 358145 | 663.55 | 407 | 147588 | 1212.57 | 901 |
| TrABySS-M | 100 | 17606 | 477189 | 379.26 | 120 | 86400 | 1391.90 | 1076 |
| TrABySS | 100 | 17599 | 127940 | 586.36 | 296 | 42257 | 1310.26 | 1009 |
| Oases-31 | 100 | 24950 | 134077 | 749.89 | 383 | 56126 | 1459.95 | 1152 |
| Trinity | 189 | 17472 | 88548 | 1131.43 | 728 | 54927 | 1624.20 | 1323 |
| SOAP | 100 | 14594 | 106326 | 519.12 | 212 | 27251 | 1444.17 | 1161 |
| BinPacker | 200 | 13932 | 63615 | 1348.21 | 945 | 46588 | 1720.11 | 1367 |
| TransLiG | 201 | 18323 | 49012 | 1782.24 | 1362 | 41579 | 2039.71 | 1619 |
| rnaSPAdes | 73 | 16443 | 63480 | 1132.98 | 686 | 36872 | 1719.26 | 1434 |
| RNA-Bloom | 200 | 16091 | 131428 | 1109.99 | 822 | 89115 | 1487.86 | 1206 |

**Table S.5:** Contig metrics and statistics for arabidopsis dataset SRR11278019.

| *Contig stats* | Lengths | | Contigs ≥ 0bp | | | Contigs ≥ 500bp | | |
|---|---|---|---|---|---|---|---|---|
| Arabidopsis | Min | Max | \|seqs\| | Mean | Median | \|seqs\| | Mean | Median |
| ClusTrast-M | 100 | 14948 | 259170 | 688.17 | 364 | 96269 | 1424.84 | 1107 |
| ClusTrast | 100 | 15012 | 126848 | 865.34 | 493 | 62917 | 1470.88 | 1193 |
| TrABySS-M | 100 | 14875 | 127032 | 665.58 | 248 | 45430 | 1531.74 | 1249 |
| TrABySS | 100 | 14861 | 58153 | 795.61 | 410 | 26378 | 1490.24 | 1228 |
| Shannon | 201 | 17397 | 62537 | 1246.16 | 876 | 41246 | 1728.92 | 1412 |
| Oases-31 | 100 | 15561 | 33912 | 1429.94 | 1195 | 26761 | 1735.48 | 1468 |
| Trinity | 201 | 15561 | 42986 | 1202.41 | 863 | 28165 | 1669.87 | 1394 |
| SOAP | 100 | 15540 | 50426 | 682.34 | 260 | 18571 | 1516.95 | 1276 |
| BinPacker | 201 | 15561 | 30770 | 1442.29 | 1159 | 25431 | 1673.36 | 1382 |
| TransLiG | 200 | 21504 | 25919 | 1671.46 | 1372 | 23801 | 1788.48 | 1475 |
| rnaSPAdes | 61 | 11095 | 42138 | 978.84 | 556 | 22003 | 1619.82 | 1371 |
| RNA-Bloom | 200 | 11931 | 48819 | 1211.58 | 938 | 35909 | 1527.76 | 1272 |

**Table S.6:** Contig metrics and statistics for zebrafish dataset SRR10728575.

| *Contig stats* | Lengths | | Contigs ≥ 0bp | | | Contigs ≥ 500bp | | |
|---|---|---|---|---|---|---|---|---|
| Zebrafish | Min | Max | \|seqs\| | Mean | Median | \|seqs\| | Mean | Median |
| ClusTrast-M | 100 | 19658 | 1355491 | 561.86 | 277 | 368804 | 1437.73 | 946 |
| ClusTrast | 100 | 16700 | 921194 | 525.59 | 311 | 266718 | 1180.85 | 819 |
| TrABySS-M | 100 | 19658 | 801185 | 506.77 | 187 | 197286 | 1474.22 | 1011 |
| TrABySS | 100 | 16632 | 360424 | 422.38 | 176 | 71458 | 1342.09 | 944 |
| Shannon | 201 | 26136 | 273909 | 1007.32 | 493 | 135636 | 1719.60 | 1175 |
| Oases-31 | 83 | 20823 | 272208 | 910.34 | 342 | 109292 | 1932.74 | 1352 |
| Trinity | 185 | 17314 | 207245 | 910.42 | 418 | 88454 | 1720.73 | 1171 |
| SOAP | 100 | 16517 | 266654 | 375.56 | 150 | 35919 | 1655.22 | 1138 |
| BinPacker | 200 | 26181 | 123556 | 1394.72 | 807 | 83128 | 1918.81 | 1333 |
| TransLiG | 200 | 31151 | 90382 | 2107.77 | 1235 | 72925 | 2533.82 | 1681 |
| rnaSPAdes | 78 | 30330 | 123430 | 1156.81 | 422 | 54291 | 2231.24 | 1522 |
| RNA-Bloom | 200 | 17368 | 211699 | 1434.28 | 761 | 132802 | 2089.21 | 1495 |

**Table S.7:** Contig metrics and statistics for the poplar dataset SRR5986240.1.

| *Contig stats* | Lengths | | Contigs ≥ 0bp | | | Contigs ≥ 500bp | | |
|---|---|---|---|---|---|---|---|---|
| Poplar | Min | Max | \|seqs\| | Mean | Median | \|seqs\| | Mean | Median |
| ClusTrast-M | 100 | 16720 | 1780433 | 513.02 | 330 | 530680 | 1094.74 | 793 |
| ClusTrast | 100 | 15808 | 673528 | 592.71 | 428 | 283885 | 1018.03 | 764 |
| TrABySS-M | 100 | 16720 | 771869 | 497.12 | 267 | 228815 | 1176.60 | 883 |
| TrABySS | 100 | 15647 | 273753 | 441.27 | 270 | 66189 | 1078.58 | 799 |
| Shannon | 201 | 20432 | 234956 | 1256.75 | 832 | 152309 | 1767.64 | 1445 |
| Oases-31 | 100 | 16219 | 245366 | 706.57 | 266 | 79883 | 1721.63 | 1440 |
| Trinity | 197 | 16730 | 154491 | 1123.79 | 741 | 97762 | 1585.03 | 1294 |
| SOAP | 100 | 19424 | 224198 | 311.83 | 171 | 30207 | 1117.79 | 855 |
| BinPacker | 201 | 17177 | 134758 | 1556.48 | 1137 | 107890 | 1859.90 | 1476 |
| TransLiG | 201 | 26772 | 99237 | 1797.46 | 1351 | 84563 | 2049.44 | 1627 |
| rnaSPAdes | 73 | 21907 | 99092 | 1223.27 | 891 | 63962 | 1719.09 | 1462 |
| RNA-Bloom | 200 | 17930 | 212396 | 1344.48 | 1029 | 153393 | 1739.77 | 1459 |

**Table S.8:** Contig metrics and statistics for the stranded human dataset SRR1153470.

| *Contig stats* | Lengths | | Contigs ≥ 0bp | | | Contigs ≥ 500bp | | |
|---|---|---|---|---|---|---|---|---|
| Human-SS | Min | Max | \|seqs\| | Mean | Median | \|seqs\| | Mean | Median |
| ClusTrast-M | 100 | 26631 | 2767795 | 446.38 | 261 | 575544 | 1234.40 | 801 |
| ClusTrast | 100 | 21827 | 1392565 | 499.09 | 279 | 367171 | 1229.08 | 830 |
| TrABySS-M | 100 | 22360 | 1226624 | 417.19 | 159 | 252745 | 1336.80 | 919 |
| TrABySS | 100 | 21722 | 646852 | 361.25 | 161 | 112016 | 1213.63 | 847 |
| Shannon | 201 | 35933 | 504565 | 1210.14 | 558 | 272083 | 1979.96 | 1323 |
| Oases | 200 | 5649 | 762004 | 515.89 | 421 | 291744 | 809.80 | 708 |
| Trinity | 185 | 26092 | 315871 | 997.46 | 442 | 139917 | 1864.49 | 1162 |
| SOAP | 100 | 24058 | 432213 | 404.49 | 157 | 67640 | 1657.49 | 1007 |
| BinPacker | 200 | 24244 | 179243 | 1386.67 | 765 | 128203 | 1803.56 | 1129 |
| TransLiG | 201 | 38941 | 127732 | 2010.24 | 1028 | 106740 | 2337.41 | 1350 |
| rnaSPAdes | 49 | 42667 | 310313 | 782.97 | 318 | 102507 | 1863.84 | 1163 |
| RNA-Bloom | 205 | 21725 | 295790 | 1189.38 | 680 | 188119 | 1663.59 | 1158 |

**Table S.9:** Contig metrics and statistics for stranded arabidopsis dataset SRR5344669.1.

| *Contig stats* | Lengths | | Contigs ≥ 0bp | | | Contigs ≥ 500bp | | |
|---|---|---|---|---|---|---|---|---|
| Arabid.-SS | Min | Max | \|seqs\| | Mean | Median | \|seqs\| | Mean | Median |
| ClusTrast-M | 100 | 16904 | 889878 | 466.82 | 261 | 207839 | 1282.15 | 962 |
| ClusTrast | 100 | 16904 | 378157 | 648.50 | 354 | 141615 | 1314.03 | 1023 |
| TrABySS-M | 100 | 16654 | 531320 | 394.48 | 135 | 113027 | 1243.86 | 971 |
| TrABySS | 100 | 16645 | 201356 | 485.26 | 236 | 55666 | 1210.09 | 940 |
| Shannon | 201 | 19978 | 92633 | 1821.79 | 1465 | 74694 | 2185.14 | 1783 |
| Oases | 200 | 5061 | 399379 | 694.35 | 596 | 245008 | 908.94 | 803 |
| Trinity | 197 | 18359 | 96292 | 1204.22 | 860 | 61385 | 1715.46 | 1463 |
| SOAP | 100 | 17298 | 111856 | 468.77 | 162 | 23117 | 1608.69 | 1326 |
| BinPacker | 200 | 18767 | 80421 | 1214.91 | 860 | 53802 | 1658.04 | 1372 |
| TransLiG | 200 | 18731 | 65110 | 1383.04 | 1086 | 47177 | 1786.34 | 1514 |
| rnaSPAdes | 63 | 20699 | 50605 | 1355.65 | 1136 | 36260 | 1780.09 | 1540 |
| RNA-Bloom | 200 | 17455 | 135445 | 1424.30 | 1128 | 103194 | 1769.88 | 1475 |

**Table S.10:** Contig metrics and statistics for the stranded poplar dataset SRR10853135.

| *Contig stats* | Lengths | | Contigs ≥ 0bp | | | Contigs ≥ 500bp | | |
|---|---|---|---|---|---|---|---|---|
| Poplar-SS | Min | Max | \|seqs\| | Mean | Median | \|seqs\| | Mean | Median |
| ClusTrast-M | 100 | 23244 | 609568 | 609.13 | 335 | 213379 | 1285.24 | 992 |
| ClusTrast | 100 | 21706 | 404809 | 572.01 | 333 | 136532 | 1201.54 | 914 |
| TrABySS-M | 100 | 23244 | 358182 | 568.49 | 264 | 119130 | 1302.90 | 1012 |
| TrABySS | 100 | 21706 | 215650 | 441.72 | 220 | 52552 | 1163.84 | 887 |
| Shannon | 201 | 26563 | 136509 | 1102.46 | 655 | 78702 | 1683.12 | 1351 |
| Oases | 200 | 6938 | 353091 | 742.19 | 621 | 223903 | 966.26 | 838 |
| Trinity | 199 | 19456 | 125401 | 991.61 | 579 | 68793 | 1547.79 | 1268 |
| SOAP | 100 | 51830 | 206811 | 333.24 | 174 | 30771 | 1145.72 | 868 |
| BinPacker | 200 | 19060 | 89596 | 1354.81 | 1005 | 67798 | 1685.12 | 1372 |
| TransLiG | 201 | 36030 | 65302 | 1572.43 | 1241 | 56048 | 1776.73 | 1448 |
| rnaSPAdes | 197 | 34306 | 81228 | 957.55 | 523 | 41762 | 1558.67 | 1292 |
| RNA-Bloom | 200 | 23191 | 126337 | 1339.72 | 912 | 84808 | 1843.00 | 1443 |

**Table S.11:** Contig metrics and statistics for the simulated dataset from Hölzer and Marz (2019).

| *Contig stats* | Lengths | | Contigs $\geq$ 0bp | | | Contigs $\geq$ 500bp | | |
|---|---|---|---|---|---|---|---|---|
| Human-SIM | Min | Max | \|seqs\| | Mean | Median | \|seqs\| | Mean | Median |
| ClusTrast-M | 100 | 24247 | 960736 | 289.47 | 210 | 112879 | 1049.15 | 742 |
| ClusTrast | 100 | 24247 | 93568 | 703.67 | 399 | 38002 | 1328.41 | 889 |
| TrABySS-M | 100 | 24247 | 539513 | 197.64 | 105 | 30186 | 1333.08 | 861 |
| TrABySS | 100 | 24238 | 40015 | 566.98 | 296 | 11513 | 1382.55 | 900 |
| Shannon | 201 | 32287 | 23789 | 1839.48 | 1111 | 16936 | 2458.51 | 1847 |
| Oases | 200 | 64989 | 75862 | 1895.78 | 1215 | 59716 | 2314.56 | 1677 |
| Trinity | 201 | 24411 | 16747 | 2370.06 | 1754 | 13747 | 2818.73 | 2186 |
| SOAP | 100 | 25255 | 15057 | 1028.18 | 187 | 4866 | 2836.91 | 2136 |
| BinPacker | 200 | 24808 | 13418 | 1935.68 | 1219 | 9861 | 2518.15 | 1885 |
| TransLiG | 201 | 27543 | 11194 | 2368.68 | 1476 | 8260 | 3097.79 | 2271 |
| rnaSPAdes | 131 | 27266 | 9149 | 2246.06 | 1650 | 7846 | 2566.58 | 1962 |
| RNA-Bloom | 201 | 24118 | 19070 | 2478.19 | 1880 | 17404 | 2679.59 | 2069 |

**Table S.12:** Contig metrics and statistics for the simulated dataset from Hayer et al. (2015).

| *Contig stats* | Lengths | | Contigs $\geq$ 0bp | | | Contigs $\geq$ 500bp | | |
|---|---|---|---|---|---|---|---|---|
| Mouse-SIM | Min | Max | \|seqs\| | Mean | Median | \|seqs\| | Mean | Median |
| ClusTrast-M | 100 | 44464 | 136626 | 1582.95 | 855 | 102240 | 2000.90 | 1292 |
| ClusTrast | 100 | 43822 | 163735 | 1190.35 | 621 | 103350 | 1694.79 | 984 |
| TrABySS-M | 100 | 40516 | 104320 | 1348.51 | 749 | 76079 | 1722.58 | 1058 |
| TrABySS | 100 | 40516 | 89376 | 1150.18 | 638 | 56073 | 1639.21 | 1055 |
| Shannon | 201 | 58212 | 36219 | 3457.95 | 2840 | 34998 | 3566.41 | 2929 |
| Oases | 200 | 26379 | 113514 | 1477.16 | 991 | 86696 | 1826.47 | 1348 |
| Trinity | 201 | 63858 | 79323 | 3315.41 | 2737 | 76987 | 3405.38 | 2804 |
| SOAP | 100 | 76809 | 44772 | 1029.94 | 199 | 16321 | 2528.34 | 1986 |
| BinPacker | 201 | 43805 | 57655 | 3202.76 | 2549 | 55597 | 3308.30 | 2630 |
| TransLiG | 201 | 76192 | 42034 | 3479.04 | 2789 | 41538 | 3516.34 | 2818 |
| rnaSPAdes | 132 | 24271 | 19267 | 2973.48 | 2498 | 19052 | 3002.71 | 2521 |
| RNA-Bloom | 222 | 43819 | 71086 | 3943.94 | 3179 | 70838 | 3956.25 | 3190 |

**Table S.13:** Contig metrics and statistics for the extra human dataset SRR8594084.

| *Contig stats* | Lengths | | Contigs $\geq$ 0bp | | | Contigs $\geq$ 500bp | | |
|---|---|---|---|---|---|---|---|---|
| Human-Extra | Min | Max | \|seqs\| | Mean | Median | \|seqs\| | Mean | Median |
| ClusTrast-M | 100 | 7782 | 122192 | 346.75 | 239 | 23700 | 870.96 | 723 |
| ClusTrast | 100 | 7782 | 69625 | 350.26 | 246 | 12673 | 880.55 | 720 |
| TrABySS-M | 100 | 7782 | 73692 | 267.16 | 149 | 9051 | 912.71 | 747 |
| TrABySS | 100 | 7782 | 39555 | 276.29 | 169 | 4934 | 881.44 | 728 |
| Shannon | 201 | 12151 | 29250 | 504.36 | 349 | 9039 | 956.66 | 767 |
| Oases | 200 | 8308 | 43463 | 654.07 | 490 | 21206 | 989.78 | 803 |
| Trinity | 195 | 7982 | 22984 | 508.45 | 355 | 7218 | 954.93 | 755 |
| SOAP | 100 | 7118 | 39802 | 269.20 | 170 | 4588 | 861.94 | 720 |
| BinPacker | 201 | 8995 | 14539 | 588.62 | 443 | 6223 | 937.83 | 742 |
| TransLiG | 200 | 41998 | 9729 | 752.49 | 538 | 5455 | 1072.60 | 745 |
| rnaSPAdes | 37 | 7893 | 36283 | 346.51 | 223 | 6191 | 913.62 | 744 |
| RNA-Bloom | 200 | 17177 | 44206 | 965.93 | 660 | 26328 | 1406.83 | 1101 |

**Table S.14:** Minimum and maximum fold change of SQANTI recall and precision for ClusTrast-**S** and ClusTrast-**M** across datasets (Figure S.2).

| | | Recall (TPR) | | Precision (PPV) | |
|---|---|---|---|---|---|
| | | min | max | min | max |
| Assembler | Comparison assembler | | | | |
| clustrast-m | oases-31 | 1.33 | 2.59 | 0.78 | 1.55 |
| | oases-m | 1.15 | 1.22 | 0.85 | 0.99 |
| | transabyss-m | 1.11 | 1.52 | 0.73 | 1.63 |
| | transabyss-s | 1.23 | 2.28 | 0.78 | 1.77 |
| | trinity | 1.23 | 2.15 | 0.91 | 1.76 |
| clustrast-s | oases-31 | 1.26 | 2.13 | 0.71 | 1.09 |
| | oases-m | 1.09 | 1.09 | 0.82 | 0.83 |
| | transabyss-m | 1.04 | 1.25 | 0.61 | 1.15 |
| | transabyss-s | 1.16 | 1.87 | 0.65 | 1.25 |
| | trinity | 1.16 | 1.76 | 0.76 | 1.24 |

**Table S.15:** Minimum and maximum fold change of CRBB recall and precision for ClusTrast-**S** and ClusTrast-**M** across datasets (Figure S.3).

| | | Recall (TPR) | | Precision (PPV) | |
|---|---|---|---|---|---|
| | | min | max | min | max |
| Assembler | Comparison assembler | | | | |
| clustrast-m | oases-31 | 1.17 | 2.14 | 0.25 | 0.67 |
| | oases-m | 1.04 | 1.05 | 0.34 | 0.39 |
| | transabyss-m | 1.06 | 1.40 | 0.63 | 1.04 |
| | transabyss-s | 1.19 | 1.99 | 0.36 | 1.40 |
| | trinity | 1.08 | 1.57 | 0.16 | 0.65 |
| clustrast-s | oases-31 | 1.09 | 1.83 | 0.37 | 0.72 |
| | oases-m | 0.90 | 0.98 | 0.35 | 0.49 |
| | transabyss-m | 0.94 | 1.20 | 0.82 | 1.81 |
| | transabyss-s | 1.11 | 1.70 | 0.98 | 1.50 |
| | trinity | 1.01 | 1.34 | 0.33 | 0.68 |

**Table S.16:** Number of expressed reference transcript isoforms reconstructed to at least 50% or 95% of their length by a single FSM, according to SQANTI.

| | Human | | | Arabidopsis | | | Mouse | | | Rice | | | Zebrafish | | | Poplar | | |
|---|---|---|---|---|---|---|---|---|---|---|---|---|---|---|---|---|---|---|
| | Total | 50% | 95% | Total | 50% | 95% | Total | 50% | 95% | Total | 50% | 95% | Total | 50% | 95% | Total | 50% | 95% |
| ClusTrast-M | **15385** | **11797** | 8317 | **16623** | 15143 | 5829 | 11391 | **9019** | 5920 | **19005** | **16013** | **13069** | **11568** | 10204 | **7484** | **20742** | **17754** | 10054 |
| TrAB-M+Sh | 15173 | 11575 | **8319** | 16612 | **15223** | **5836** | **11392** | 9000 | **5925** | n/a | n/a | n/a | 11565 | **10252** | 7475 | 20383 | 17539 | **10189** |
| TrABySS-M | 13804 | 9826 | 6607 | 15556 | 14062 | 3851 | 10744 | 8337 | 5267 | 17294 | 14335 | 11579 | 10685 | 9375 | 6629 | 18840 | 15419 | 7762 |
| ClusTrast | 13951 | 10810 | 7237 | 15943 | 14362 | 5131 | 10601 | 8309 | 5096 | 17807 | 14745 | 10871 | 10720 | 9298 | 6328 | 18679 | 15069 | 7373 |
| TrAB+Sh | 13856 | 10654 | 7241 | 16170 | 14762 | 5503 | 10689 | 8472 | 5311 | n/a | n/a | n/a | 10933 | 9544 | 6430 | 18212 | 15161 | 8775 |
| TrABySS | 11344 | 7870 | 4653 | 14375 | 12743 | 3016 | 9286 | 7020 | 4009 | 15210 | 11885 | 8413 | 9325 | 7946 | 5004 | 14655 | 10630 | 4822 |
| Shannon | 11721 | 9338 | 6234 | 14837 | 13588 | 4830 | 9523 | 7736 | 4600 | n/a | n/a | n/a | 9861 | 8641 | 5546 | 15713 | 13569 | 7910 |
| Oases-M | n/a | n/a | n/a | 14922 | 13926 | 4760 | 9839 | 8065 | 5221 | 15520 | 13070 | 10058 | n/a | n/a | n/a | n/a | n/a | n/a |
| Oases-S | 9341 | 6869 | 4432 | 13638 | 12743 | 3677 | 8076 | 6438 | 3698 | 13397 | 10673 | 7783 | 8639 | 7686 | 5306 | 12759 | 10070 | 5137 |
| Trinity | 10659 | 8688 | 6101 | 14443 | 13312 | 4958 | 8587 | 7080 | 4579 | 15371 | 13301 | 10835 | 9741 | 8689 | 6082 | 15704 | 13649 | 8025 |
| SOAP | 8940 | 5796 | 3601 | 12804 | 11099 | 2727 | 8171 | 5878 | 3229 | 13608 | 9837 | 6901 | 7562 | 6303 | 3827 | 10831 | 5995 | 2069 |
| BinPacker | 9123 | 7310 | 4724 | 12420 | 11347 | 3573 | 7845 | 6504 | 3760 | 12979 | 10758 | 7068 | 8560 | 7543 | 4714 | 12917 | 10996 | 5973 |
| TransLiG | 8665 | 7353 | 5130 | 12928 | 12255 | 4118 | 7674 | 6666 | 4139 | 13023 | 11805 | 9414 | 8425 | 7705 | 5419 | 12839 | 11532 | 6921 |
| rnaSPAdes | 9339 | 7830 | 5767 | 14537 | 13281 | 4473 | 9005 | 7166 | 4675 | 14593 | 12840 | 11053 | 8801 | 8145 | 6130 | 15480 | 13947 | 7941 |
| RNA-Bloom | 11068 | 9582 | 6741 | 14672 | 13630 | 4657 | 9211 | 7911 | 5156 | 16206 | 14273 | 12138 | 10038 | 9087 | 6408 | 17775 | 15973 | 9452 |

**Table S.17:** Reconstructed isoforms per gene according to SQANTI.

| | Human | Arabid. | Mouse | Rice | Zebrafish | Poplar | Human-SS | Arabid.-SS | Poplar-SS | Human-SIM | Mouse-SIM | Human-Extra |
|---|---|---|---|---|---|---|---|---|---|---|---|---|
| ClusTrast-M | **1.71654** | 1.11707 | **1.47432** | **1.09582** | **1.15309** | **1.17313** | **1.97096** | 1.20135 | 1.11921 | **2.47249** | 1.02572 | 1.22222 |
| TrAB-M+Sh | 1.62519 | 1.12425 | 1.46778 | n/a | 1.13722 | 1.15832 | 1.88281 | 1.19454 | **1.1244** | 1.81989 | 1.03022 | 1.21051 |
| TrABySS-M | 1.4411 | 1.09938 | 1.39324 | 1.07211 | 1.10126 | 1.1052 | 1.70497 | 1.12941 | 1.08605 | 1.82055 | 1.01594 | 1.17687 |
| ClusTrast | 1.54663 | 1.08637 | 1.36798 | 1.07385 | 1.10765 | 1.13223 | 1.80876 | 1.16908 | 1.0966 | 2.08074 | 1.02776 | 1.17339 |
| TrAB+Sh | 1.4502 | 1.09672 | 1.37431 | n/a | 1.09952 | 1.12256 | 1.67476 | 1.15684 | 1.09709 | 1.3323 | 1.03041 | 1.16404 |
| TrABySS | 1.19808 | 1.05857 | 1.24532 | 1.04313 | 1.04751 | 1.04661 | 1.38378 | 1.06933 | 1.04605 | 1.3323 | 1.01589 | 1.09815 |
| Shannon | 1.34519 | 1.05783 | 1.25087 | n/a | 1.06399 | 1.10043 | 1.45644 | 1.13512 | 1.06899 | 1.56339 | 1.00956 | 1.09475 |
| Oases-M | n/a | 1.12786 | 1.4659 | 1.07848 | n/a | n/a | 1.23877 | 1.05549 | 1.0571 | 1.77696 | **1.04676** | **1.23455** |
| Oases-S | 1.32416 | 1.09371 | 1.26672 | 1.04606 | 1.07431 | 1.06521 | 1.0645 | 1.01575 | 1.02616 | 1.51605 | 1.00689 | 1.09785 |
| Trinity | 1.44301 | 1.10434 | 1.30069 | 1.04535 | 1.0889 | 1.07196 | 1.57482 | 1.25831 | 1.07021 | 1.92659 | 1.01519 | 1.11906 |
| SOAP | 1.08434 | 1.0125 | 1.09791 | 1.02881 | 1.01375 | 1.03536 | 1.09465 | 1.0231 | 1.03024 | 1.13964 | 1.0107 | 1.04794 |
| BinPacker | 1.29124 | 1.07166 | 1.19477 | 1.03614 | 1.05152 | 1.05636 | 1.3799 | 1.15635 | 1.05029 | 1.41625 | 1.02376 | 1.0585 |
| TransLiG | 1.28182 | 1.0714 | 1.18647 | 1.03294 | 1.05496 | 1.04884 | 1.35804 | 1.16138 | 1.04062 | 1.41372 | 1.0102 | 1.06358 |
| rnaSPAdes | 1.15344 | 1.03075 | 1.16602 | 1.02844 | 1.02659 | 1.03986 | 1.21715 | 1.07721 | 1.02438 | 1.41211 | 1.00414 | 1.05627 |
| RNA-Bloom | 1.57418 | **1.13318** | 1.422 | 1.06794 | 1.11795 | 1.13058 | 1.68069 | **1.31187** | 1.09699 | 2.08498 | 1.00729 | 1.18883 |

**Table S.18:** Number of precision true positives detected by SQANTI and CRBB, respectively. $A$=True positives detected by SQANTI. $B$=True positives detected by CRBB.

| | Human | | | Mouse | | | Rice | | | Arabidopsis | | | Zebrafish | | | Poplar | | |
|---|---|---|---|---|---|---|---|---|---|---|---|---|---|---|---|---|---|---|
| | $A \setminus B$ | $A \cap B$ | $B \setminus A$ | $A \setminus B$ | $A \cap B$ | $B \setminus A$ | $A \setminus B$ | $A \cap B$ | $B \setminus A$ | $A \setminus B$ | $A \cap B$ | $B \setminus A$ | $A \setminus B$ | $A \cap B$ | $B \setminus A$ | $A \setminus B$ | $A \cap B$ | $B \setminus A$ |
| ClusTrast-M | 264662 | 52032 | 45473 | 74023 | 44215 | 20174 | 932669 | 67164 | 37412 | 89616 | 48541 | 12552 | 99577 | 49327 | 30381 | 594508 | 98120 | 55535 |
| TrAB-M+Sh | 142336 | 43119 | 51457 | 50326 | 42902 | 22437 | n/a | n/a | n/a | 47157 | 49121 | 13768 | 59323 | 48928 | 32469 | 216363 | 83889 | 60734 |
| TrABySS-M | 111063 | 20665 | 20129 | 32148 | 24026 | 11848 | 148831 | 28294 | 18635 | 32809 | 25492 | 7001 | 44682 | 28726 | 17206 | 176076 | 48751 | 26664 |
| ClusTrast | 79924 | 31415 | 33038 | 37053 | 26132 | 11760 | 101965 | 39422 | 22823 | 26035 | 35528 | 8121 | 33666 | 26919 | 15748 | 140923 | 45026 | 25354 |
| TrAB+Sh | 58731 | 32448 | 40232 | 34275 | 29463 | 15964 | n/a | n/a | n/a | 24717 | 38952 | 10388 | 26801 | 31613 | 21186 | 85207 | 47934 | 40964 |
| TrABySS | 27300 | 9999 | 9052 | 16052 | 10574 | 5429 | 20863 | 13812 | 8321 | 10342 | 15317 | 3638 | 12060 | 11383 | 5898 | 44862 | 12805 | 6973 |
| Shannon | 31826 | 22638 | 31465 | 18224 | 18936 | 10571 | n/a | n/a | n/a | 14371 | 23699 | 6777 | 14926 | 20452 | 15368 | 40351 | 35288 | 34194 |
| Oases-M | n/a | n/a | n/a | 43674 | 64886 | 38827 | 61668 | 55010 | 53088 | 24947 | 53925 | 21596 | n/a | n/a | n/a | n/a | n/a | n/a |
| Oases-S | 29566 | 19582 | 22324 | 10407 | 12893 | 7355 | 25894 | 18378 | 14683 | 4712 | 17171 | 4995 | 10256 | 19259 | 15032 | 33359 | 24257 | 22145 |
| Trinity | 15610 | 15103 | 22105 | 8355 | 10737 | 8527 | 11602 | 17910 | 15963 | 5574 | 16387 | 5266 | 7359 | 14535 | 12190 | 15973 | 25347 | 23506 |
| SOAP | 17486 | 5856 | 8109 | 12206 | 6330 | 3148 | 17085 | 10309 | 6433 | 7340 | 11569 | 2479 | 6991 | 7092 | 5333 | 33503 | 5626 | 3297 |
| BinPacker | 12725 | 11845 | 19116 | 6363 | 8919 | 6774 | 14591 | 13947 | 13184 | 5392 | 14205 | 4653 | 7538 | 13507 | 11594 | 19669 | 27074 | 28640 |
| TransLiG | 9089 | 10291 | 18247 | 4351 | 8484 | 6276 | 9792 | 13769 | 14002 | 4018 | 14400 | 4441 | 6025 | 12477 | 13224 | 13228 | 21693 | 24909 |
| rnaSPAdes | 8254 | 9983 | 11887 | 8898 | 8835 | 4973 | 9190 | 14351 | 10285 | 5164 | 14055 | 3280 | 3908 | 10648 | 8520 | 9245 | 19207 | 15493 |
| RNA-Bloom | 17476 | 23373 | 31608 | 13290 | 23910 | 16394 | 26882 | 31249 | 24708 | 6821 | 20533 | 6544 | 10763 | 27114 | 21541 | 21333 | 47562 | 44896 |



**Table S.19:** Number of recall true positives detected by SQANTI and CRBB, respectively. $A$=True positives detected by SQANTI. $B$=True positives detected by CRBB.

| | Human | | | Mouse | | | Rice | | | Arabidopsis | | | Zebrafish | | | Poplar | | |
|---|---|---|---|---|---|---|---|---|---|---|---|---|---|---|---|---|---|---|
| | $A \setminus B$ | $A \cap B$ | $B \setminus A$ | $A \setminus B$ | $A \cap B$ | $B \setminus A$ | $A \setminus B$ | $A \cap B$ | $B \setminus A$ | $A \setminus B$ | $A \cap B$ | $B \setminus A$ | $A \setminus B$ | $A \cap B$ | $B \setminus A$ | $A \setminus B$ | $A \cap B$ | $B \setminus A$ |
| ClusTrast-M | 29917 | 23205 | 11197 | 12336 | 13430 | 3892 | 7683 | 19347 | 3857 | 3019 | 15386 | 1144 | 7994 | 13741 | 4426 | 14972 | 22304 | 6464 |
| TrAB-M+Sh | 27859 | 20191 | 11436 | 11932 | 13338 | 3930 | n/a | n/a | n/a | 2935 | 15447 | 1177 | 7647 | 13491 | 4307 | 14065 | 21693 | 6831 |
| TrABySS-M | 25709 | 14554 | 9948 | 11161 | 11710 | 4146 | 7322 | 16495 | 4936 | 2984 | 14498 | 1562 | 7388 | 11985 | 4400 | 14883 | 18138 | 6011 |
| ClusTrast | 24164 | 18290 | 11137 | 11444 | 11441 | 3494 | 7756 | 17538 | 4275 | 3237 | 14821 | 1251 | 7402 | 11904 | 4422 | 15494 | 18322 | 6036 |
| TrAB+Sh | 21358 | 15698 | 10816 | 10463 | 11650 | 3685 | n/a | n/a | n/a | 2985 | 15061 | 1287 | 6952 | 11819 | 4255 | 12706 | 17808 | 7126 |
| TrABySS | 17032 | 8868 | 8448 | 9168 | 8853 | 3968 | 7538 | 13209 | 5454 | 3359 | 13312 | 1803 | 6325 | 9395 | 4397 | 13083 | 11382 | 5407 |
| Shannon | 13557 | 11829 | 9674 | 6892 | 9122 | 3110 | n/a | n/a | n/a | 2677 | 13870 | 1843 | 5206 | 9979 | 4383 | 8691 | 14734 | 8295 |
| Oases-M | n/a | n/a | n/a | 9411 | 11788 | 4464 | 6431 | 15674 | 4973 | 2010 | 14352 | 1622 | n/a | n/a | n/a | n/a | n/a | n/a |
| Oases-S | 13050 | 8247 | 7808 | 6366 | 8134 | 3683 | 7377 | 11655 | 4974 | 1922 | 13070 | 1978 | 4828 | 9020 | 4569 | 8899 | 9986 | 6458 |
| Trinity | 11429 | 11013 | 10936 | 5934 | 8610 | 4682 | 4714 | 14294 | 6481 | 2389 | 13544 | 2346 | 4821 | 10090 | 4970 | 7904 | 14300 | 8936 |
| SOAP | 11321 | 5111 | 7501 | 7165 | 5849 | 3410 | 7547 | 10405 | 5960 | 3501 | 11634 | 2331 | 4924 | 6632 | 5118 | 12817 | 5357 | 3346 |
| BinPacker | 8666 | 8005 | 9359 | 4800 | 7252 | 4079 | 5696 | 11120 | 5761 | 2528 | 11816 | 2284 | 4275 | 8586 | 4682 | 6806 | 11009 | 7575 |
| TransLiG | 6960 | 7465 | 9430 | 3667 | 7197 | 3996 | 4161 | 11418 | 6300 | 1949 | 12344 | 2337 | 3543 | 8128 | 5302 | 5661 | 10870 | 8879 |
| rnaSPAdes | 7494 | 7863 | 9455 | 5887 | 7805 | 4299 | 4342 | 12899 | 7019 | 2763 | 13366 | 2406 | 3512 | 8754 | 5925 | 6060 | 13873 | 9635 |
| RNA-Bloom | 12119 | 13721 | 10638 | 6712 | 10449 | 4197 | 4856 | 15694 | 5675 | 2192 | 14039 | 1824 | 4846 | 10758 | 4345 | 8102 | 17345 | 9145 |

**Table S.20:** Number of reconstructed reference transcripts, following at least one of two criteria: $A$=FSMs detected by SQANTI, with max 50 bases difference to annotated TSS. $B$=CRBB hits with coverage ≥0.95. The columns can be interpreted as follows: $B \setminus A$: most likely reconstructed only with alternative splicing involed. $A \setminus B$ and $A \cap B$: certainly reconstructed without alternative splicing involed.

| | Human | | | Mouse | | | Rice | | | Arabidopsis | | | Zebrafish | | | Poplar | | |
|---|---|---|---|---|---|---|---|---|---|---|---|---|---|---|---|---|---|---|
| | $A \setminus B$ | $A \cap B$ | $B \setminus A$ | $A \setminus B$ | $A \cap B$ | $B \setminus A$ | $A \setminus B$ | $A \cap B$ | $B \setminus A$ | $A \setminus B$ | $A \cap B$ | $B \setminus A$ | $A \setminus B$ | $A \cap B$ | $B \setminus A$ | $A \setminus B$ | $A \cap B$ | $B \setminus A$ |
| ClusTrast-M | 2729 | 5274 | 4726 | 1277 | 4717 | 2233 | 1226 | 11063 | 3958 | 879 | 2945 | 2764 | 1180 | 5326 | 2743 | 3643 | 4515 | 1900 |
| TrAB-M+Sh | 2912 | 5458 | 4965 | 1359 | 4774 | 2301 | n/a | n/a | n/a | 898 | 2906 | 2796 | 1194 | 5283 | 2754 | 3879 | 4524 | 2072 |
| TrABySS-M | 1779 | 3485 | 3766 | 1101 | 3947 | 2256 | 1054 | 9525 | 4215 | 708 | 1733 | 2400 | 1009 | 4390 | 2691 | 2826 | 3151 | 1507 |
| ClusTrast | 2308 | 4469 | 4136 | 1008 | 3777 | 1827 | 1043 | 8604 | 3463 | 787 | 2593 | 2508 | 964 | 4388 | 2298 | 2640 | 3107 | 1436 |
| TrAB+Sh | 2412 | 4656 | 4181 | 1129 | 4099 | 1980 | n/a | n/a | n/a | 835 | 2733 | 2714 | 1015 | 4506 | 2319 | 3308 | 3802 | 1904 |
| TrABySS | 1063 | 2342 | 2641 | 749 | 2745 | 1811 | 822 | 6386 | 3429 | 553 | 1334 | 2050 | 721 | 3249 | 2111 | 1724 | 1845 | 1076 |
| Shannon | 2003 | 3931 | 3799 | 882 | 3441 | 1707 | n/a | n/a | n/a | 729 | 2370 | 2613 | 894 | 3820 | 2079 | 2994 | 3347 | 1813 |
| Oases-M | n/a | n/a | n/a | 1316 | 4179 | 2706 | 1134 | 7913 | 3870 | 753 | 2270 | 2692 | n/a | n/a | n/a | n/a | n/a | n/a |
| Oases-S | 1441 | 2540 | 3065 | 761 | 2543 | 2097 | 807 | 5934 | 3048 | 560 | 1703 | 2257 | 821 | 3448 | 2399 | 1895 | 1987 | 1185 |
| Trinity | 2222 | 4403 | 4350 | 955 | 3666 | 2696 | 839 | 9164 | 4160 | 687 | 2570 | 2967 | 968 | 4371 | 2492 | 2903 | 3654 | 2007 |
| SOAP | 760 | 1757 | 2440 | 647 | 1798 | 1455 | 783 | 4575 | 3040 | 498 | 1212 | 1946 | 781 | 2128 | 1791 | 839 | 530 | 337 |
| BinPacker | 1451 | 3057 | 3210 | 768 | 2729 | 1889 | 765 | 4786 | 2936 | 592 | 1781 | 2211 | 745 | 3181 | 1944 | 2180 | 2460 | 1439 |
| TransLiG | 1700 | 3309 | 3637 | 846 | 3049 | 2166 | 1226 | 7173 | 3608 | 653 | 2039 | 2526 | 927 | 3704 | 2533 | 2573 | 2935 | 1773 |
| rnaSPAdes | 1484 | 3444 | 4280 | 760 | 3447 | 2293 | 993 | 9213 | 4455 | 633 | 2212 | 2767 | 884 | 4227 | 3028 | 2650 | 3647 | 1923 |
| RNA-Bloom | 2510 | 4635 | 4488 | 1278 | 4144 | 2401 | 1050 | 10592 | 4303 | 748 | 2366 | 2611 | 1076 | 4578 | 2548 | 3540 | 4545 | 2290 |

**Table S.21:** Number of assembled contigs, following at least one of two criteria: $A$=FSMs detected by SQANTI, with max 50 bases difference to annotated TSS. $B$=CRBB hits with coverage ≥0.95. The columns can be interpreted as follows: $B \setminus A$: most likely alternative splicing involved. $A \setminus B$ and $A \cap B$: certainly no alternative splicing involved.

| | Human | | | Mouse | | | Rice | | | Arabidopsis | | | Zebrafish | | | Poplar | | |
|---|---|---|---|---|---|---|---|---|---|---|---|---|---|---|---|---|---|---|
| | $A \setminus B$ | $A \cap B$ | $B \setminus A$ | $A \setminus B$ | $A \cap B$ | $B \setminus A$ | $A \setminus B$ | $A \cap B$ | $B \setminus A$ | $A \setminus B$ | $A \cap B$ | $B \setminus A$ | $A \setminus B$ | $A \cap B$ | $B \setminus A$ | $A \setminus B$ | $A \cap B$ | $B \setminus A$ |
| ClusTrast-M | 6525 | 9414 | 10504 | 3252 | 12662 | 7838 | 6109 | 25376 | 13952 | 2018 | 5358 | 8063 | 3224 | 14736 | 10736 | 13428 | 12620 | 7105 |
| TrAB-M+Sh | 4819 | 9762 | 12681 | 3000 | 12502 | 8096 | n/a | n/a | n/a | 2094 | 5228 | 8337 | 3353 | 14624 | 10787 | 11693 | 12148 | 7249 |
| TrABySS-M | 2062 | 4763 | 5330 | 1561 | 7017 | 4412 | 1990 | 14172 | 7858 | 1016 | 2243 | 4078 | 1812 | 8795 | 6434 | 6221 | 6926 | 3677 |
| ClusTrast | 3086 | 6773 | 7723 | 1737 | 7578 | 4337 | 2046 | 15519 | 8278 | 1435 | 4242 | 5698 | 1667 | 8111 | 5163 | 5037 | 5727 | 2971 |
| TrAB+Sh | 3683 | 7550 | 9865 | 2081 | 8500 | 5503 | n/a | n/a | n/a | 1651 | 4419 | 6555 | 2245 | 9387 | 6576 | 7215 | 7272 | 4577 |
| TrABySS | 918 | 2525 | 2525 | 642 | 2997 | 1826 | 726 | 6613 | 3545 | 569 | 1432 | 2301 | 696 | 3519 | 2202 | 1736 | 2041 | 1024 |
| Shannon | 2778 | 5116 | 7408 | 1437 | 5510 | 3686 | n/a | n/a | n/a | 1084 | 2995 | 4263 | 1563 | 5961 | 4409 | 5472 | 5249 | 3566 |
| Oases-M | n/a | n/a | n/a | 5146 | 15932 | 13998 | 4356 | 18133 | 16744 | 2532 | 4771 | 9306 | n/a | n/a | n/a | n/a | n/a | n/a |
| Oases-S | 2155 | 3783 | 5648 | 858 | 3445 | 2945 | 1164 | 7405 | 5520 | 681 | 1993 | 3172 | 1319 | 6493 | 5770 | 3811 | 3644 | 2654 |
| Trinity | 1985 | 4842 | 5951 | 841 | 3993 | 3199 | 1125 | 10088 | 5858 | 775 | 2863 | 3695 | 1053 | 5433 | 3924 | 4240 | 4912 | 2932 |
| SOAP | 634 | 1884 | 2468 | 561 | 1884 | 1382 | 705 | 4664 | 2981 | 476 | 1237 | 1929 | 716 | 2199 | 1799 | 821 | 554 | 323 |
| BinPacker | 1633 | 3672 | 4864 | 715 | 3010 | 2230 | 1357 | 5591 | 4544 | 673 | 2021 | 2760 | 1091 | 4681 | 3526 | 5339 | 4876 | 3267 |
| TransLiG | 1834 | 3837 | 4885 | 767 | 3304 | 2408 | 2235 | 7910 | 5594 | 731 | 2281 | 3019 | 1555 | 5342 | 4887 | 5121 | 5062 | 3358 |
| rnaSPAdes | 1231 | 3777 | 4451 | 596 | 3639 | 2245 | 1149 | 9602 | 4745 | 628 | 2303 | 2883 | 842 | 4785 | 3531 | 2952 | 4392 | 2204 |
| RNA-Bloom | 3336 | 5690 | 7495 | 2737 | 7012 | 5873 | 2980 | 14861 | 9581 | 882 | 2748 | 3850 | 2090 | 9566 | 7377 | 6770 | 8866 | 5894 |

**Table S.22:** Estimated percentage of polymorphic variants, in reconstructed transcripts (using values from Table S.20) and assembled contigs (Table S.21) for ClusTrast-**M**.

| Table | Human | Mouse | Rice | Arabidopsis | Zebrafish | Poplar |
|---|---|---|---|---|---|---|
| S.20 | 62.8722 | 72.8577 | 75.6386 | 58.0449 | 70.3427 | 81.1096 |
| S.21 | 60.2768 | 67.0007 | 69.2937 | 47.7751 | 62.5871 | 78.5691 |

**Table S.23:** Number of true positive candidates from ClusTrast-**M** detected by SQANTI (calculated for precision; *full* or *incomplete* splice match, recovered exon fraction≥0.25), but not detected with CRBB. *Category* and *Subcategory* are classifications made by SQANTI. A=aligned with CRBB, *but* coverage<0.25. NA=not aligned w. CRBB at all. FSM=full splice match. ISM=incomplete splice match. IR=intron retention.

| Caterogy | Subcategory | | Human | Mouse | Rice | Arabidopsis | Zebrafish | Poplar |
|---|---|---|---|---|---|---|---|---|
| FSM | 3′ fragment | A | 19688 | 4783 | 12393 | 3440 | 10283 | 22768 |
| FSM | 3′ fragment | NA | 19227 | 7480 | 54296 | 5914 | 9426 | 57865 |
| FSM | 5′ fragment | A | 17119 | 5971 | 18233 | 3455 | 6649 | 21697 |
| FSM | 5′ fragment | NA | 14125 | 4687 | 66761 | 5283 | 4696 | 41274 |
| FSM | internal frag. | A | 25139 | 5364 | 18500 | 3658 | 11841 | 28729 |
| FSM | internal frag. | NA | 19025 | 6219 | 53100 | 5343 | 7905 | 50013 |
| ISM | IR | A | 1012 | 174 | 1018 | 234 | 398 | 3332 |
| ISM | IR | NA | 364 | 69 | 1456 | 222 | 204 | 2809 |
| ISM | mono-exon | A | 34768 | 19001 | 123044 | 16266 | 17822 | 76749 |
| ISM | mono-exon | NA | 173706 | 25069 | 753154 | 49278 | 74106 | 390669 |
| ISM | multi-exon | A | 9215 | 3048 | 6732 | 1725 | 2410 | 7308 |
| ISM | multi-exon | NA | 11307 | 4161 | 59803 | 6652 | 2537 | 39785 |

**Table S.24:** Number of true positive candidates (calculated for precision; cov≥0.25) from ClusTrast-**M** detected by CRBB, but not by SQANTI, and how SQANTI actually classifies them. F=Fusion. GI=genic intron. G=genic. IG=intergenic. NIC=novel in catalog. NNC=novel not in catalog. IR=intron retention.

| Category | Subcategory | Human | Mouse | Rice | Arabidopsis | Zebrafish | Poplar |
|---|---|---|---|---|---|---|---|
| F | IR | 824 | 123 | 395 | 430 | 199 | 1569 |
| F | multi-exon | 1977 | 587 | 3340 | 1834 | 1704 | 5985 |
| GI | mono-exon | 0 | 0 | 3 | 0 | 3 | 0 |
| G | mono-exon | 8167 | 1975 | 12549 | 1284 | 5475 | 10259 |
| G | multi-exon | 3808 | 1921 | 9838 | 1754 | 1388 | 9755 |
| IG | mono-exon | 3808 | 997 | 4442 | 370 | 5798 | 4376 |
| IG | multi-exon | 7113 | 2679 | 2014 | 202 | 4049 | 3819 |
| NIC | known junct. | 5161 | 3519 | 457 | 99 | 2374 | 431 |
| NIC | known sites | 4184 | 1995 | 1679 | 442 | 1477 | 586 |
| NIC | IR | 6556 | 1533 | 4141 | 1730 | 2744 | 10556 |
| NIC | mono-exon | 18867 | 3416 | 5543 | 1414 | 3906 | 9301 |
| NIC | mono-exon+IR | 4439 | 750 | 1958 | 704 | 620 | 5033 |
| NNC | nov. site(s) | 23261 | 10987 | 35861 | 8102 | 24914 | 50403 |
| NNC | IR | 4877 | 1227 | 3576 | 1483 | 2735 | 12251 |

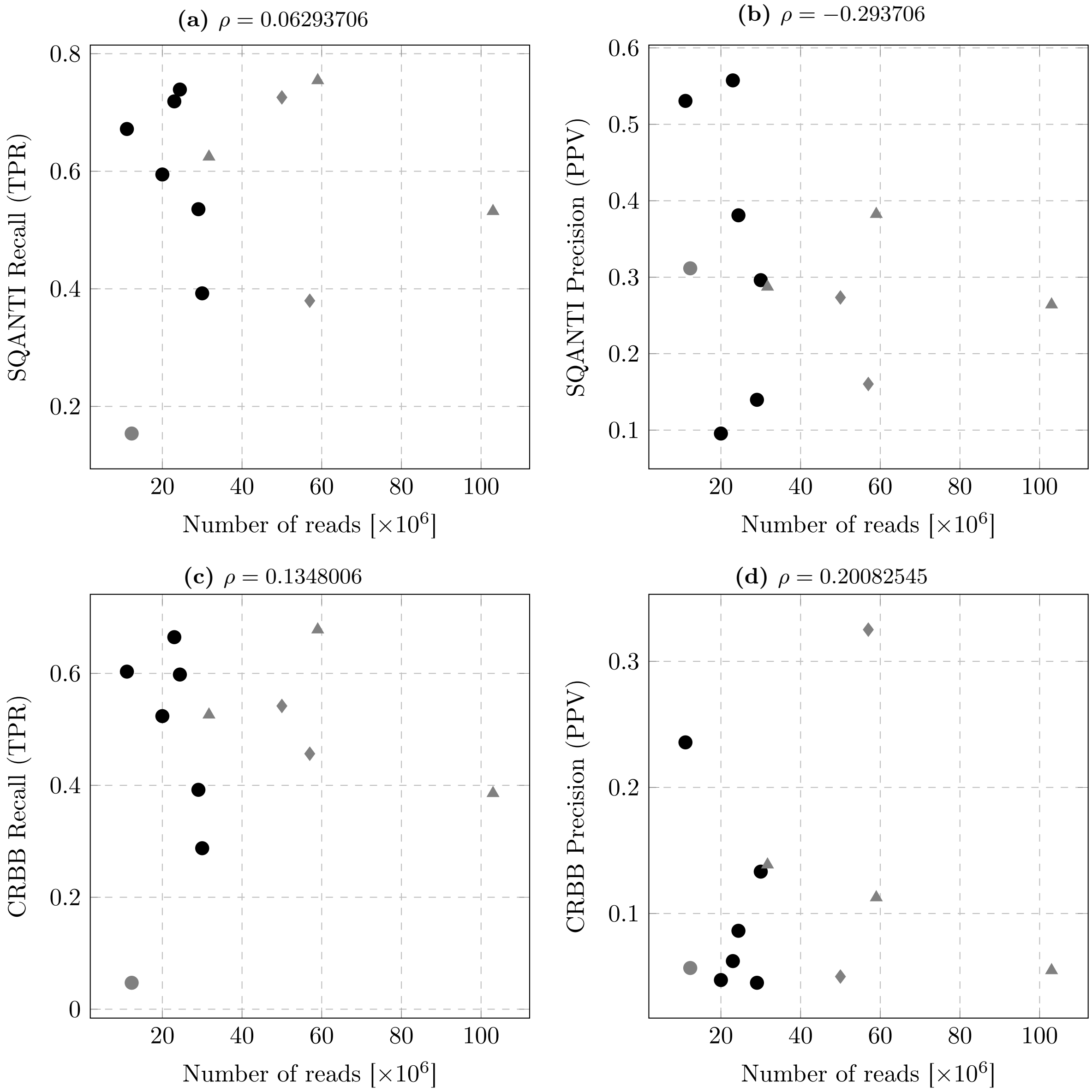


**Figure S.17:** Precision and Recall for ClusTrast-**M** from SQANTI and CRBB w.r.t. sequencing depth, i.e. number of reads (after preprocessing) in each dataset. These plots show low correlation between quality of the result and sequencing depth for ClusTrast-**M**. Bullet=nonstranded dataset. Triangle=stranded dataset. Diamond=simulated dataset. Black=dataset used in main paper. Gray=additional dataset (only used in supplementary).

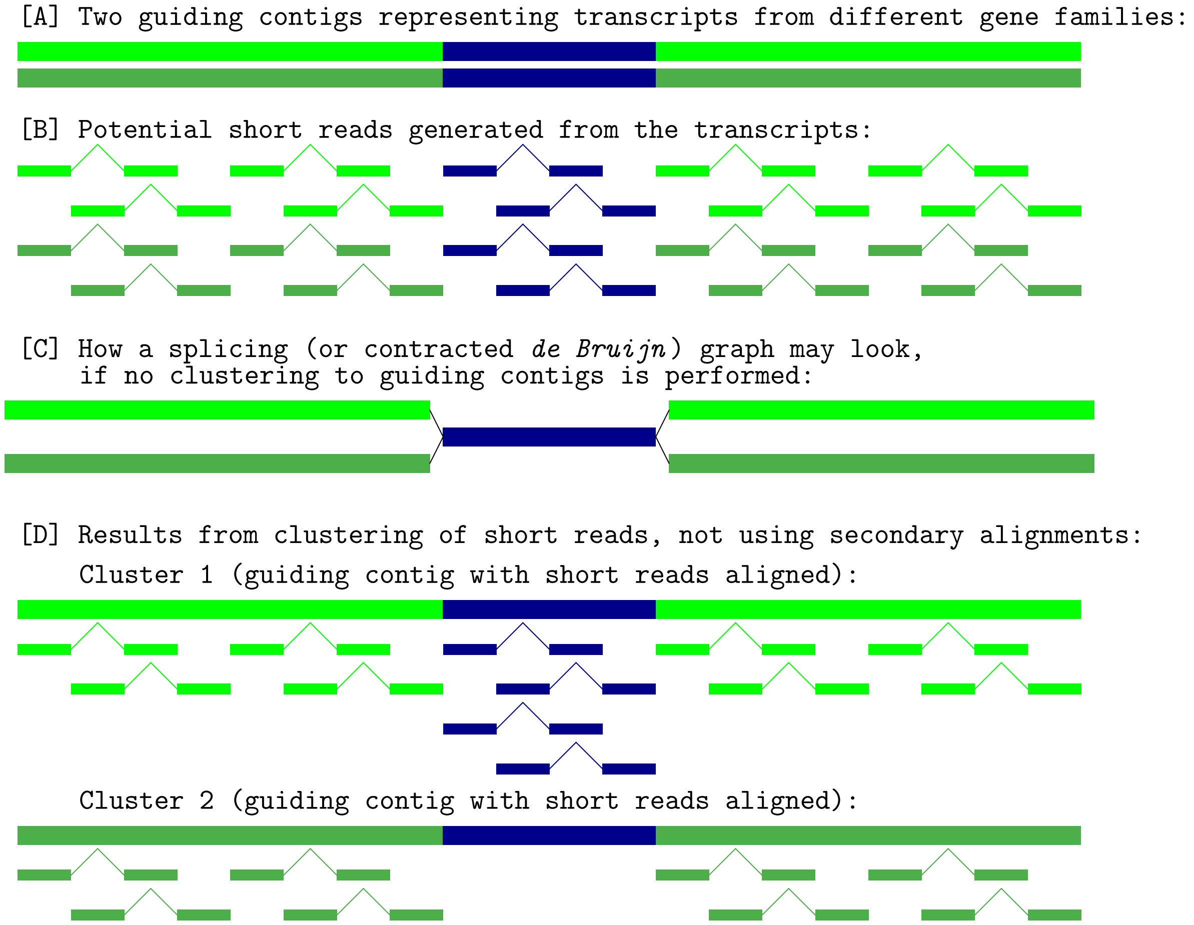


**Figure S.18:** Schematic illustration of the benefits of clustering. The two transcripts in [A] are two different guiding contigs. The blue region is identical between the two, while the dark and light green regions differ. The reads in [B] are coloured according to their origin. In [C] is illustrated what a splicing (or contracted *de Bruijn*) graph could look look like if no clustering to guiding contigs occurred. A traditional assembler might from this generate four contigs of which two are chimeric. In [D] clustering has been performed. Here, the short reads originating from the common region might only be aligned to one of the guiding contigs, if secondary alignments are not used. In that case, the dark green transcript would not be reconstructed. If secondary alignments are allowed, the blue reads may be aligned to cluster 2 as well, enabling reconstruction of the dark green transcript.

#### C.1 Run time and memory use

**Table S.25:** Computational resources used for the different assemblies.

**(a)** Execution time in minutes.

| Runtime [m] | Human | Mouse | Rice | Arabid. | Zebrafish | Poplar |
|---|---|---|---|---|---|---|
| ClusTrast-M | 2145 | 660 | 1164 | 882 | 1511 | 808 |
| ClusTrast | 1369 | 457 | 351 | 226 | 388 | 671 |
| TrABySS-M | 480 | 185 | 234 | 78 | 232 | 268 |
| TrABySS | 203 | 92 | 154 | 50 | 150 | 175 |
| Shannon | 110 | 62 | 2140 | 21 | 82 | 80 |
| Oases-M | 2880 | 586 | 23400 | 152 | 2880 | 2880 |
| Trinity | 672 | 150 | 157 | 109 | 924 | 240 |
| SOAP | 29 | 8 | 20 | 7 | 20 | 25 |
| BinPacker | 960 | 313 | 575 | 108 | 577 | 776 |
| TransLiG | 845 | 249 | 428 | 93 | 668 | 699 |
| rnaSPAdes | 77 | 34 | 56 | 18 | 53 | 78 |
| RNA-Bloom | 94 | 56 | 58 | 18 | 81 | 62 |

**(b)** Maximum memory usage in GB.

| Memory [GB] | Human | Mouse | Rice | Arabid. | Zebrafish | Poplar |
|---|---|---|---|---|---|---|
| ClusTrast-M | 135.0 | 162.6 | 135.7 | 134.9 | 267.2 | 57.15 |
| ClusTrast | 28.2 | 17.7 | 30.1 | 11.1 | 32.1 | 16.75 |
| TrABySS-M | 134.9 | 64.9 | 100.5 | 80.0 | 93.3 | 43.8 |
| TrABySS | 25.7 | 12.9 | 16.9 | 13.3 | 16.7 | 8.5 |
| Shannon | 92.5 | 32.6 | 2000 | 41.8 | 38.9 | 39.45 |
| Oases-M | 264 | 46.6 | 186 | 123 | 120 | 2000 |
| Trinity | 237 | 237 | 237 | 237 | 237 | 237 |
| SOAP | 25.2 | 8.8 | 14.2 | 14.1 | 14.3 | 8.7 |
| BinPacker | 102 | 39.1 | 40.2 | 46.4 | 42.1 | 33.9 |
| TransLiG | 103 | 31.4 | 41.8 | 47.1 | 40.7 | 32.6 |
| rnaSPAdes | 31.2 | 12 | 15.7 | 10.5 | 15.1 | 6.8 |
| RNA-Bloom | 13.4 | 7.8 | 6.27 | 5.0 | 7.7 | 5.1 |

**Table S.26:** Information on the machines used for our benchmarking. SOAP, rnaSPAdes and RNA-Bloom, known to be lightweight, were run on `SAGA`, all other assemblers were run on `PDC`.

| Machine | CPU | Core | Memory | OS |
|---|---|---|---|---|
| `PDC` | Intel Xeon CPU E5-2690 v3 @ 2.60GHz | 48 | 512 GB | CentOS 7.7 |
| `SAGA` | Intel Xeon Silver 4110 CPU @ 2.10GHz | 16 | 92 GB | Scientific Linux 7.9 |

The run time and memory usage of each assembler was measured by the UNIX command `time`. We used two computer setups for testing (see Table S.26). To minimize timing artifacts related to storage access latency, all input, temporary, and output files were stored in shared memory using the dedicated file system mounted under `/dev/shm`. This was particularly important for ClusTrast and Trinity, both relying on access to many small temporary files during parts of their execution.

The fastest assembler was SOAP-denovo-Trans. RNA-Bloom was relatively fast and had the lowest peak memory use. ClusTrast-**S** and Trans-ABySS-**S** were slightly faster than ClusTrast-**M** and Trans-ABySS-**M**. ClusTrast-**S** also had consistently lower peak memory use than Shannon.

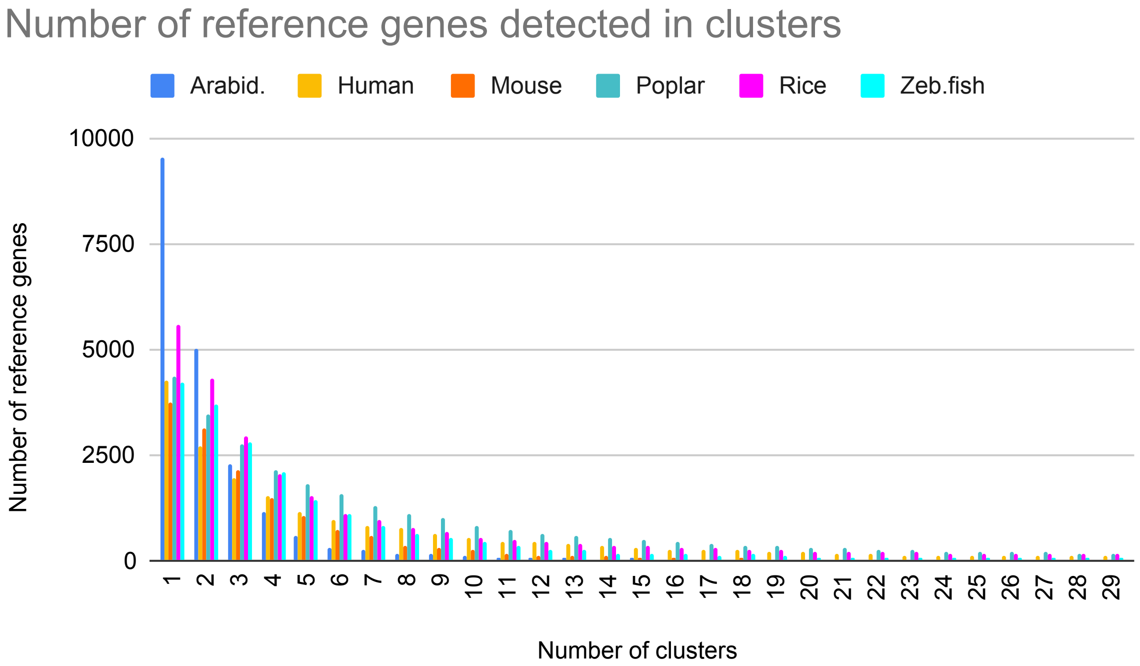


**Figure S.19:** Genes per cluster

#### C.2 Cluster characteristics

The clustering of the guiding contigs is performed by isONclust, which originally intended to cluster PacBio CCS (circular concensus sequence) reads so that one cluster=one gene. In our case, the purpose of the clustering is to reduce the risk of large *de Bruijn*-graph components in order to simplify the clusterwise assembly using Shannon. Such large components may also introduce a larger risk of outputting chimeric transcripts – such as for the example in Figure S.18 panel [C], where the reconstruction step could produce a chimeric transcript consisting of one part from the light green and one part from the dark green transcript, joined through their common blue part. Figure S.19 shows the number of (reference) genes per cluster in the datasets used in the main paper.

The clustering of the (input) short reads is performed by minimap2, which maps the reads to the guiding contigs, to be joined with the output from isONclust. The preset minimap2 option `-x sr` is used, together with flags allowing up to 100000 secondary mappings to be included. However, minimap2 also uses a secondary-to-primary score ratio, which limits the secondary mappings. The `-x sr` option sets this ratio to 0.5. Thus, all secondary mappings (in practice) with $\geq$ 50% of the maximum mapping score is kept to be clustered. Figure S.20 shows how many reads that are mapped to 1,2,3, etc. different clusters for the datasets used in the main paper. A read can only occur once per cluster.

We keep secondary mappings to assure that, in the clustering of short reads to guiding contigs, any read whose origin is ambiguous, will be assigned to all clusters that have a guiding contig to which it might be sufficiently well aligned to (given it fulfills the mapping requirements in the minimap2 mapping, see above). This has the effect of allowing construction of less fragmented *de Bruijn*-graphs for all the individual short read clusters to which the read can be mapped. If secondary mappings were not used, each read would be assigned to only one single short read cluster. As a consequence, the read can be used to construct an assembled transcript in only one single cluster, and this would potentially prevent the construction of an

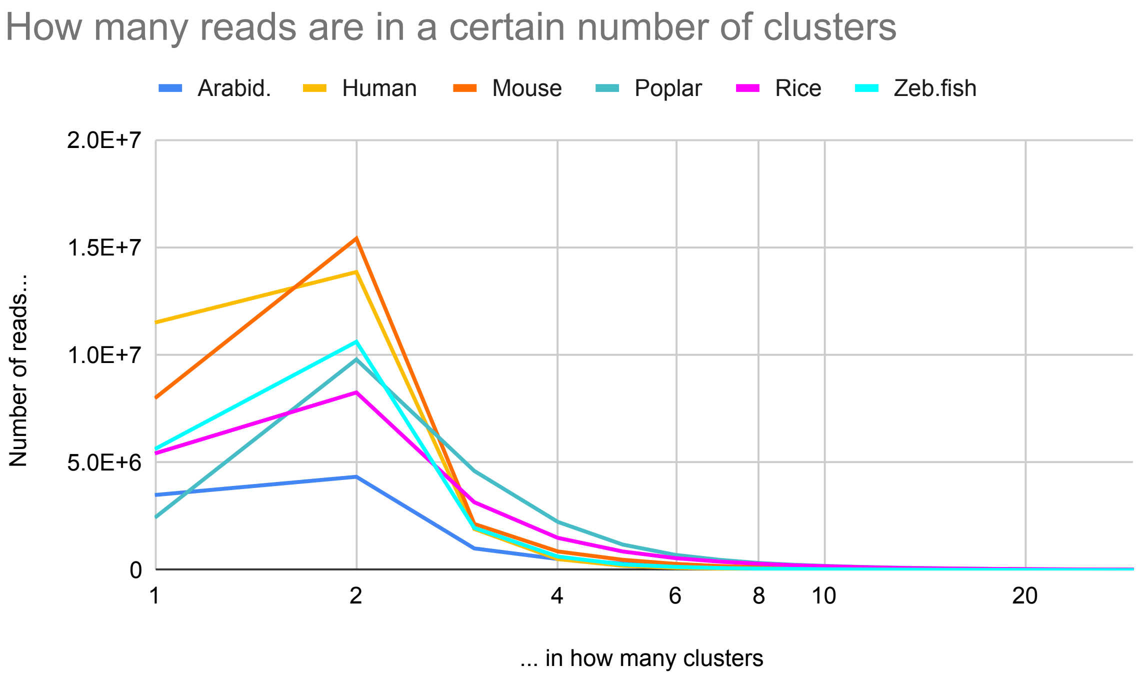


**Figure S.20:** Reads per cluster

**Table S.27:** Difference for ClusTrast-**M** when ran with and without secondary mappings considered. Red (negative number)=better with secondary mappings. Green (positive number)=better without secondary mappings. Black (zero)=indifferent. x=recovered exon percentage for SQANTI, coverage percentage for CRBB.

**(a)** Difference in SQANTI precision (PPV).

| x | human | mouse | rice | arabidopsis | poplar | zebrafish |
|---|---|---|---|---|---|---|
| 50% | $2.47 \cdot 10^{-2}$ | $-2.19 \cdot 10^{-2}$ | $-1.05 \cdot 10^{-1}$ | $-6.67 \cdot 10^{-3}$ | $-5.67 \cdot 10^{-2}$ | $2.72 \cdot 10^{-2}$ |
| 95% | $1.55 \cdot 10^{-2}$ | $-2.65 \cdot 10^{-3}$ | $-6.04 \cdot 10^{-2}$ | $1.16 \cdot 10^{-2}$ | $-3.11 \cdot 10^{-2}$ | $1.69 \cdot 10^{-2}$ |

**(b)** Difference in CRBB precision (PPV).

| x | human | mouse | rice | arabidopsis | poplar | zebrafish |
|---|---|---|---|---|---|---|
| 50% | $1.81 \cdot 10^{-2}$ | $1.62 \cdot 10^{-2}$ | $5.96 \cdot 10^{-2}$ | $4.15 \cdot 10^{-2}$ | $2.68 \cdot 10^{-2}$ | $1.91 \cdot 10^{-2}$ |
| 95% | $6.32 \cdot 10^{-3}$ | $9.55 \cdot 10^{-3}$ | $2.85 \cdot 10^{-2}$ | $9.74 \cdot 10^{-3}$ | $4.52 \cdot 10^{-3}$ | $7.12 \cdot 10^{-3}$ |

**(c)** Difference in SQANTI recall (TPR).

| x | human | mouse | rice | arabidopsis | poplar | zebrafish |
|---|---|---|---|---|---|---|
| 50% | $-3.38 \cdot 10^{-2}$ | $-7.77 \cdot 10^{-3}$ | $-5.45 \cdot 10^{-3}$ | $3.02 \cdot 10^{-3}$ | $-1.35 \cdot 10^{-2}$ | $-7.59 \cdot 10^{-3}$ |
| 95% | $-8.56 \cdot 10^{-3}$ | $-3.71 \cdot 10^{-3}$ | $1.46 \cdot 10^{-3}$ | $2.02 \cdot 10^{-3}$ | $-1.01 \cdot 10^{-2}$ | $-3.84 \cdot 10^{-3}$ |

**(d)** Difference in CRBB recall (TPR).

| x | human | mouse | rice | arabidopsis | poplar | zebrafish |
|---|---|---|---|---|---|---|
| 50% | $-2.16 \cdot 10^{-2}$ | $-4.02 \cdot 10^{-3}$ | $-3.15 \cdot 10^{-4}$ | $4.66 \cdot 10^{-4}$ | $-1.54 \cdot 10^{-2}$ | $-3.69 \cdot 10^{-3}$ |
| 95% | $9.1 \cdot 10^{-5}$ | $3.3 \cdot 10^{-5}$ | $6.39 \cdot 10^{-3}$ | $-2.21 \cdot 10^{-3}$ | $-3.53 \cdot 10^{-3}$ | $-7.49 \cdot 10^{-4}$ |

assembled transcript in a cluster which actually contains a guiding contig that has a region that is compatible with the short read. This could happen, e.g., if two genes share a common stretch of nucleotide sequence, for instance if the two genes contain the same protein domain or an otherwise conserved sequence in the coding, 5', or 3' UTR regions. See Figure S.18 for an illustration. Thus, the fact that we use secondary mappings is a design choice consistent with the aim to produce an as comprehensive set of transcripts as possible.

There is, however, an option to switch off secondary mappings in ClusTrast. We tested this option on the six datasets used in the main paper, see Table S.27. We observed (*i*) overall minor differences in recall but with worse SQANTI and CRBB recall for 5/6 datasets at 50% exon/transcript coverage (and 4 and 3, respectively, with worse performance at 95% coverage ), (*ii*) marginally improved CRBB precision for all 6 datasets, and likewise marginally improved SQANTI precision for 2/6 datasets, and (*iii*) smaller difference between SQANTI and CRBB precision measurements for 4/6 datasets.

#### C.3 Supplemental datasets

Apart from the datasets in the main paper, we also tested ClusTrast on some further datasets, listed in Table S.1, in order to be able to see if the species, strand specificity or number of reads could have any impact on the assembly. Despite the fact that ClusTrast does not have any strand specific option, the results (Figures S.9, S.13 and S.16) seem not to differ in performance compared to the non-stranded ones.

Dataset origin (i.e. species) and size are discussed in the main paper (Discussion).

As shown in Figure S.17, performance measures for ClusTrast-**M** were not correlated with the number of reads in the data sets ($-0.29 < \rho < 0.2$).

#### C.4 Sashimi plots

We observed cases where ClusTrast detected highly expressed isoforms missed by other methods. To illustrate this, we used the aligner STAR (Dobin et al., 2012) together with ggsashimi (Garrido-Martín et al., 2018) to create sashimi plots over the the genes containing the *highest expressed isoform* (according to RSEM) which, out of all the tested assemblers, only ClusTrast managed to reconstruct (as an FSM according to SQANTI), Figures S.26–S.37.

For the human (Figure S.26), mouse (Figure S.27), and rice (Figure S.28) datasets in the main paper, ClusTrast managed to reconstruct the highest expressed isoform of the example gene, while all the other assemblers only reconstructed lesser expressed isoforms. For arabidopsis (Figure S.29), the highest and second highest expressed isoform of the example gene was detected by more than one method (including ClusTrast), but the 3rd and 4th highest expressed isoforms of that gene were detected only by ClusTrast (note that their TPMs still are very high - in the range 1695–2808). For zebrafish (Figure S.30) and poplar (Figure S.31), ClusTrast the highest expressed isoform belonged to a *gene*, fairly highly expressed, which other assemblers had missed completely.

For the supplemental datasets, ClusTrast solely reconstructed a gene's highest expressed isoform in the stranded human (Figure S.32), the stranded poplar (Figure S.34), the extra human (Figure S.35) and the simulated dataset from Hölzer and Marz (2019; Figure S.36). In the stranded arabidopsis (Figure S.33) and in the simulated dataset from Hayer et al. (2015; Figure S.37), ClusTrast solely reconstructed an isoform from a *gene* not detected by any of the other tested assemblers (a single-isoform gene in arabidopsis).

In three of the twelve examples, there are other isoforms from the same gene which *only* some other assembler(s) reconstructed and not ClusTrast, but none of these isoforms were particulary highly expressed, neither when compared to all isoforms in the dataset nor when compared to

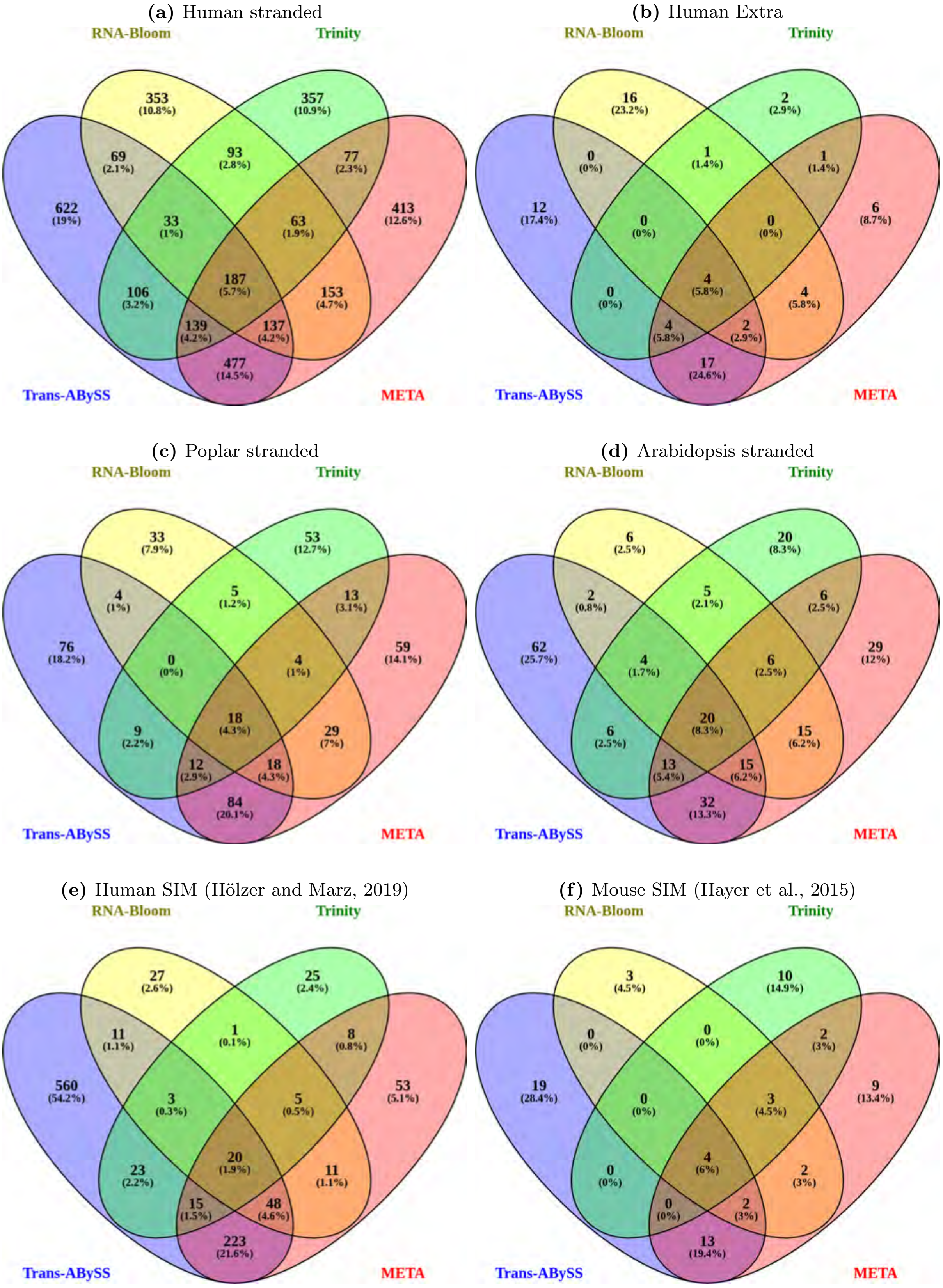


**Figure S.21:** ClusTrast results using four different tools - RNA-Bloom, Trinity, Trans-ABySS-**M**, and META - for the primary assembly. The Venn diagrams show the number of annotated and expressed transcript isoforms reconstructed solely by ClusTrast and no other tested method. ClusTrast with primary assembly from Trans-ABySS-**M** generates the highest number of unique transcript isoforms for all datasets (**a**) - (**f**).

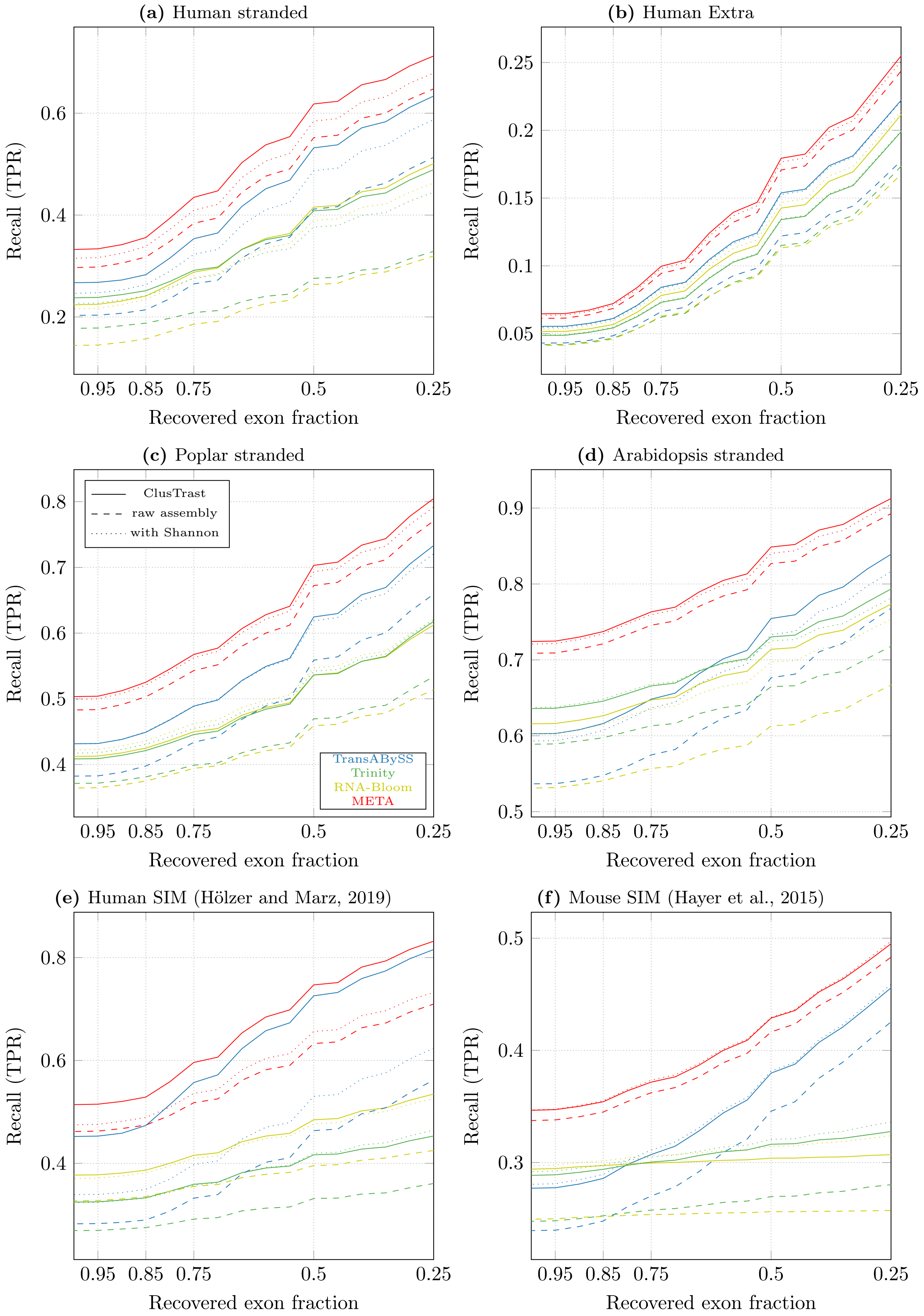


**Figure S.22:** Proportion of reference isoforms with at least one SQANTI classification of FSM or ISM vs. the cumulative proportion of exons recovered by the assembly, for ClusTrast when ran with different tools for primary assembly (solid lines), compared with these tools on their own (dashed lines) and concatenated with Shannon (dotted lines).

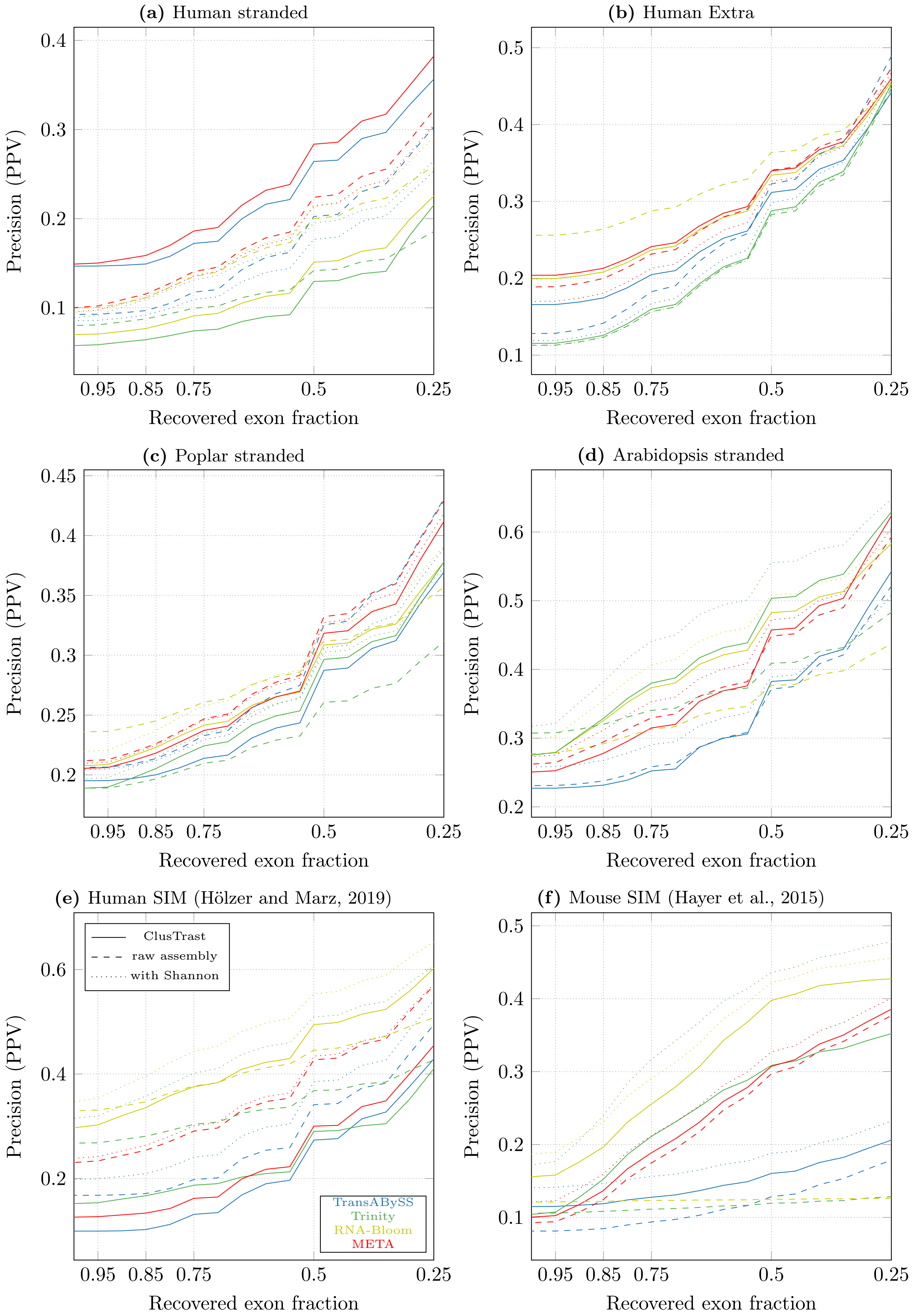


**Figure S.23:** Proportion of reconstructed isoforms classified by SQANTI as FSM or ISM vs. the cumulative proportion of recovered exons from the reference, for ClusTrast when ran with different tools for primary assembly (solid lines), compared with these tools on their own (dashed lines) and concatenated with Shannon (dotted lines).

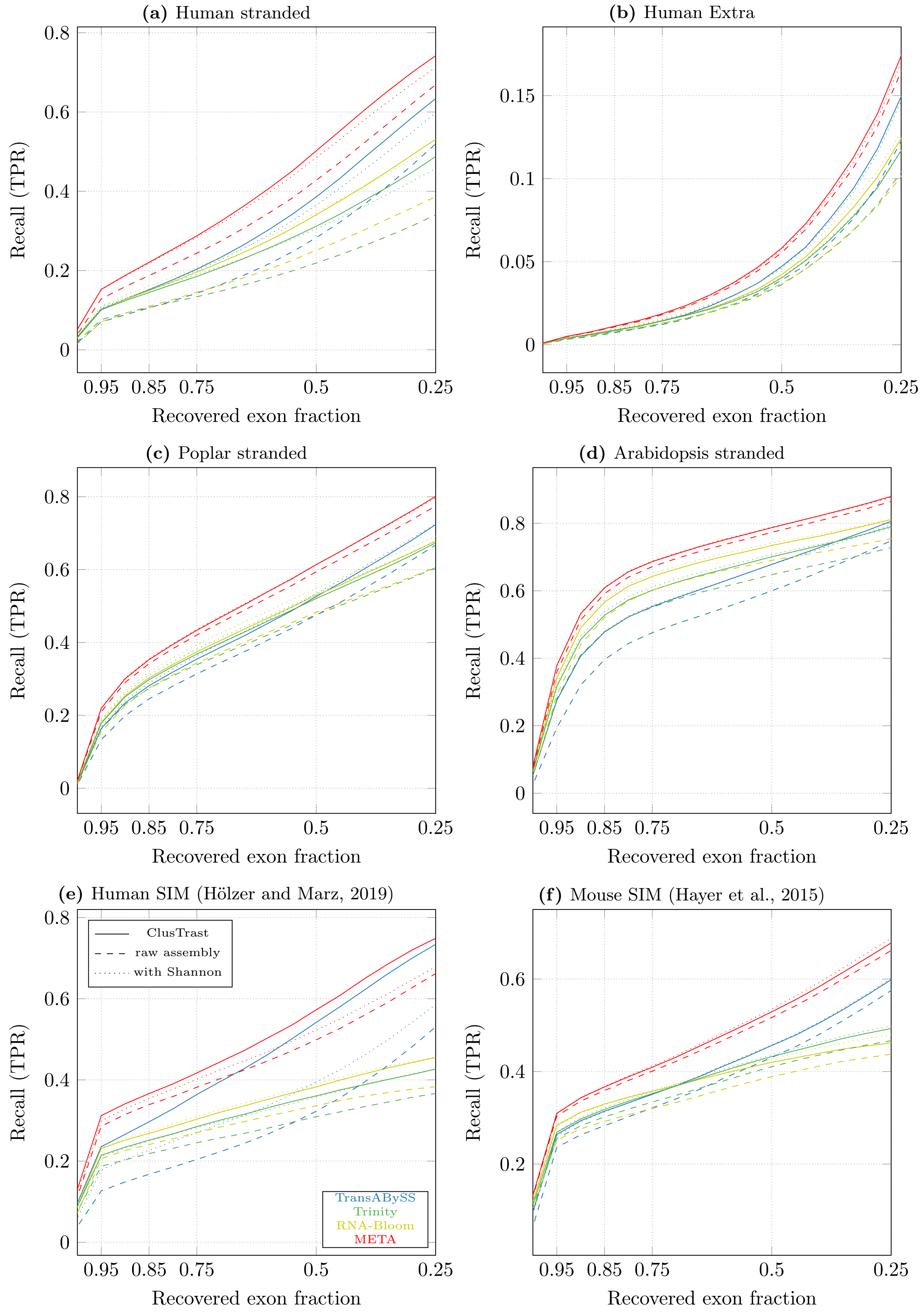


**Figure S.24:** Proportion of references with a CRBB hit vs. the cumulative proportion of recovered reference length, for ClusTrast when ran with different tools for primary assembly (solid lines), compared with these tools on their own (dashed lines) and concatenated with Shannon (dotted lines).

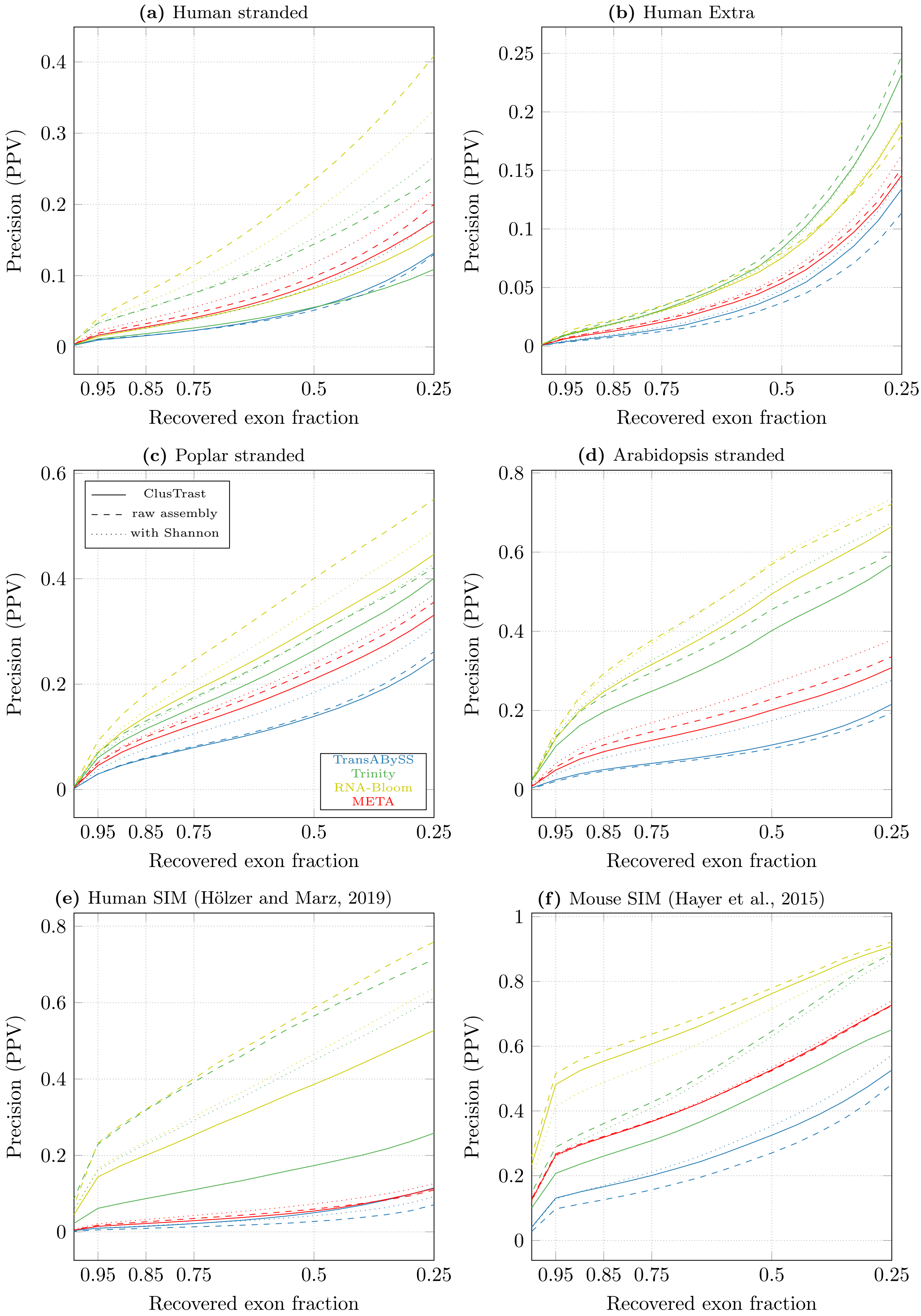


**Figure S.25:** Proportion of reconstructed isoforms with a CRBB hit vs. the cumulative proportion of recovered reference length, for ClusTrast when ran with different tools for primary assembly (solid lines), compared with these tools on their own (dashed lines) and concatenated with Shannon (dotted lines).

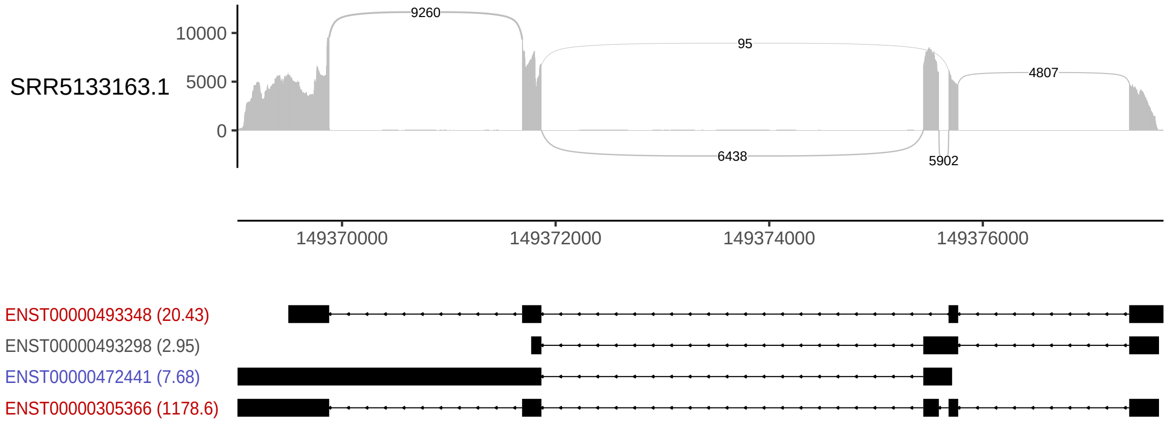


**Figure S.26:** Sashimi-plot of the human dataset SRR5133163.1 over the gene ENSG00000169908. TPM-values are shown within parethesis next to each isoform name. Red=detected solely by ClusTrast. Blue=detected by ClusTrast and at least one other assembler. Grey=not detected by any assembler.

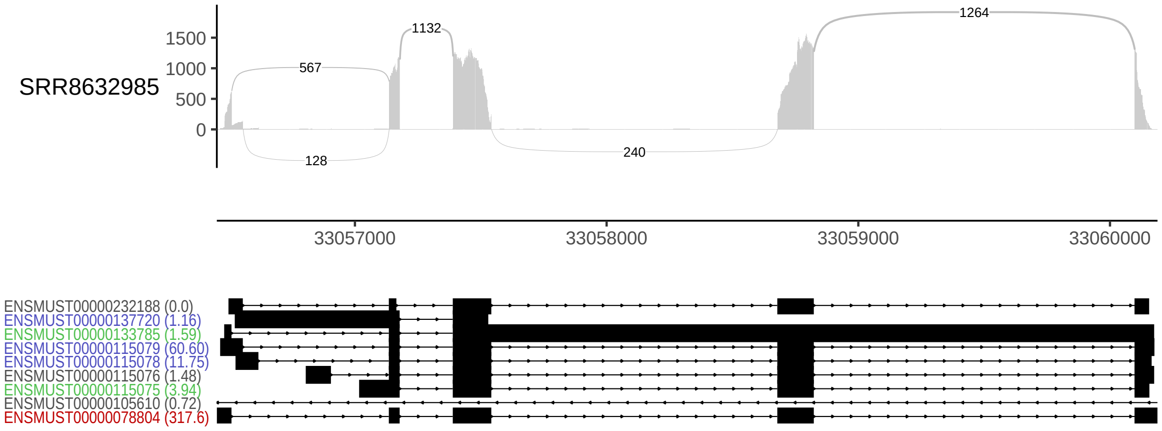


**Figure S.27:** Sashimi-plot of the mouse dataset SRR8632985 over the gene ENSMUSG00000060636. TPM-values are shown within parethesis next to each isoform name. Red=detected solely by ClusTrast. Blue=detected by ClusTrast and at least one other assembler. Green=deteced not by ClusTrast, but some other assembler. Grey=not detected by any assembler.

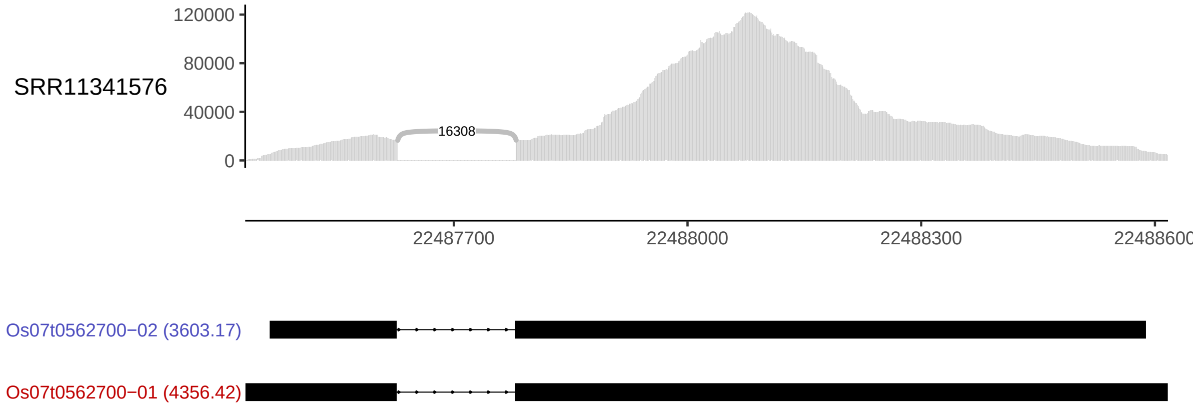


**Figure S.28:** Sashimi-plot of the rice dataset SRR11341576 over the gene Os07t0562700. TPM-values are shown within parethesis next to each isoform name. Red=detected solely by ClusTrast. Blue=detected by ClusTrast and at least one other assembler.

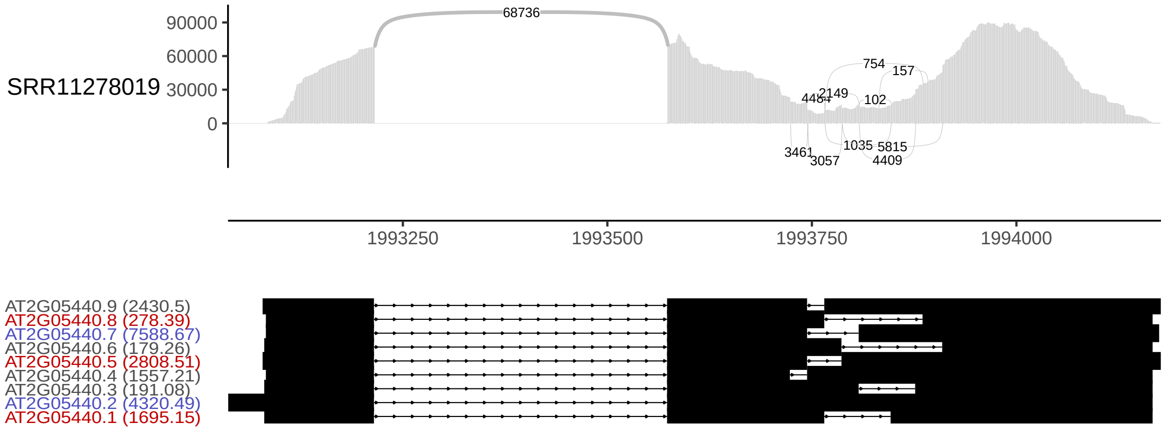


**Figure S.29:** Sashimi-plot of the arabidopsis dataset SRR11278019 over the gene AT2G05440. TPM-values are shown within parethesis next to each isoform name. Red=detected solely by ClusTrast. Blue=detected by ClusTrast and at least one other assembler. Grey=not detected by any assembler.

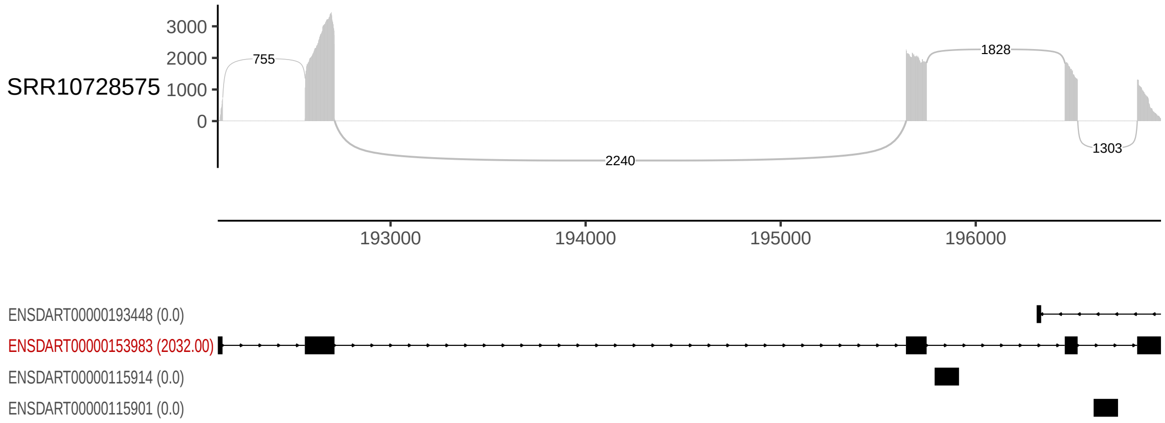


**Figure S.30:** Sashimi-plot of the zebrafish dataset SRR10728575 over the single-isoform gene ENSDART00000153983 (other isoforms shown are parts of other genes). TPM-values are shown within parethesis next to each isoform name. Red=detected solely by ClusTrast. Grey=not detected by any assembler.

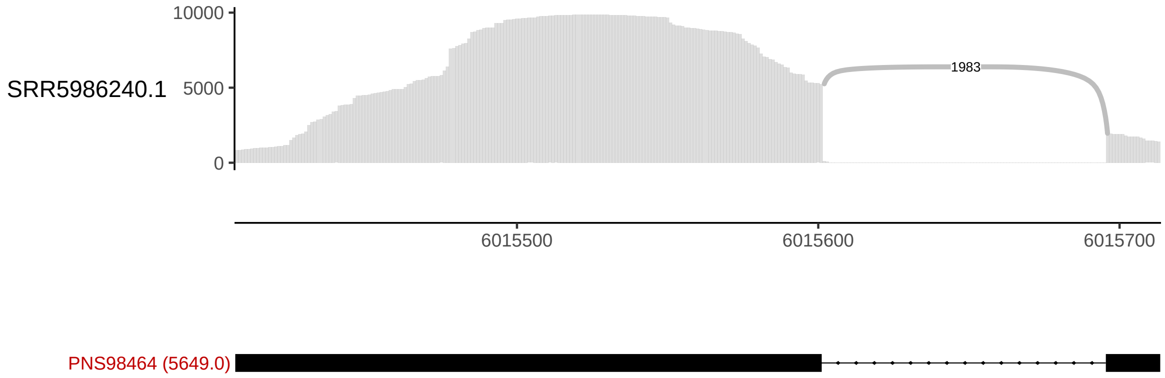


**Figure S.31:** Sashimi-plot of the poplar dataset SRR5986240 over the single-isoform gene POPTR_016G078900v3. TPM-values are shown within parethesis next to each isoform name. Red=detected solely by ClusTrast.

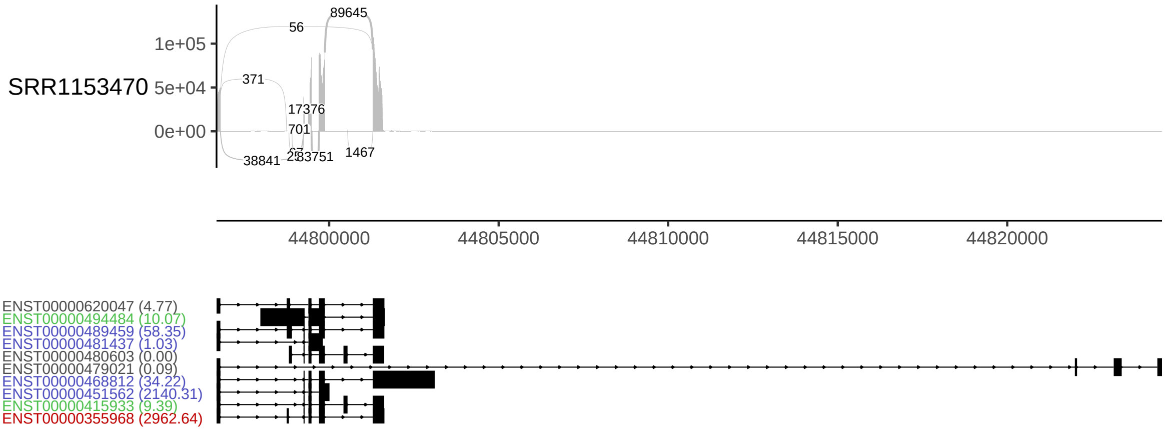


**Figure S.32:** Sashimi-plot of the stranded human dataset SRR1153470 over the gene ENSG00000196262. TPM-values are shown within parethesis next to each isoform name. Red=detected solely by ClusTrast. Blue=detected by ClusTrast and at least one other assembler. Grey=not detected by any assembler.

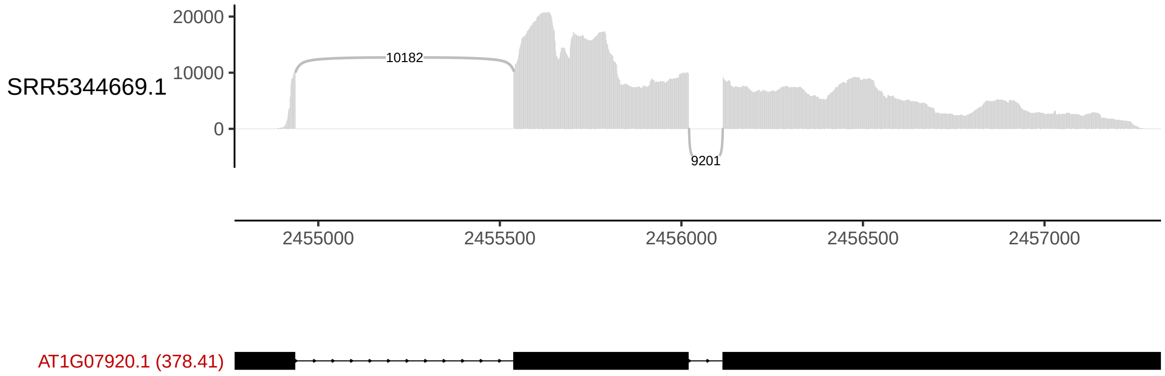


**Figure S.33:** Sashimi-plot of the stranded arabidopsis dataset SRR5344669.1 over the single-isoform gene AT1G07920. TPM-values are shown within parethesis next to each isoform name. Red=detected solely by ClusTrast. Blue=detected by ClusTrast and at least one other assembler. Grey=not detected by any assembler.

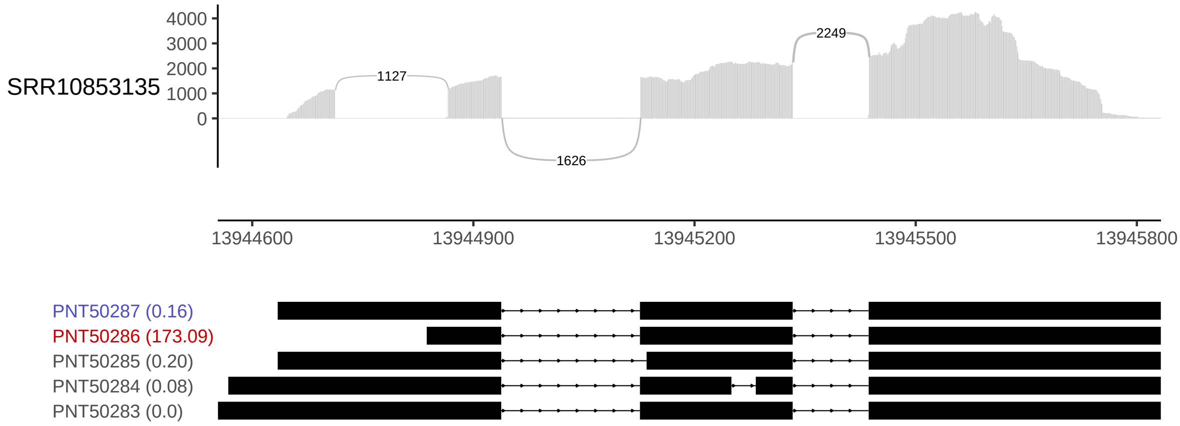


**Figure S.34:** Sashimi-plot of the stranded poplar (metatranscriptomic) dataset SRR10853135 over the gene POPTR_002G179400v3. TPM-values are shown within parethesis next to each isoform name. Red=detected solely by ClusTrast. Blue=detected by ClusTrast and at least one other assembler.

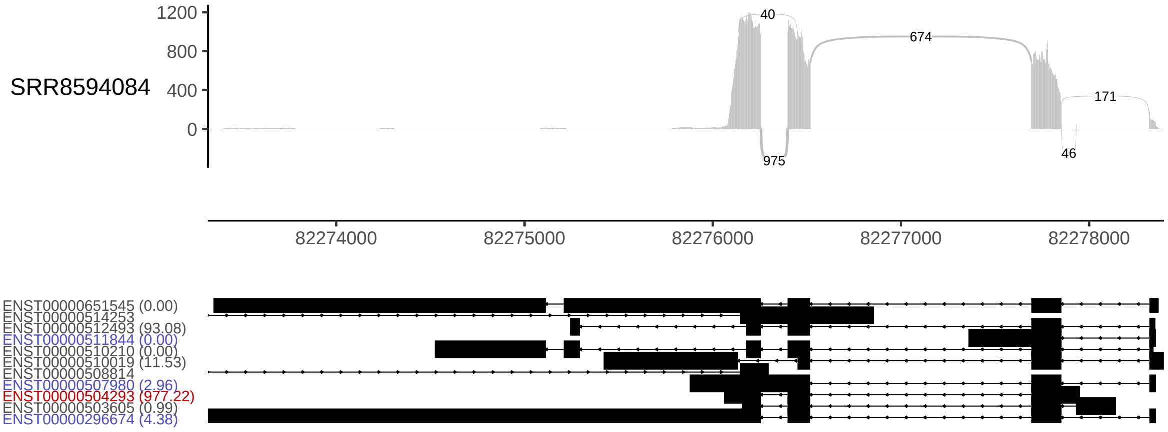


**Figure S.35:** Sashimi-plot of the extra human dataset SRR8594084 over the gene ENSG00000186468 (other isoforms shown are parts of other genes). TPM-values are shown within parethesis next to each isoform name. Red=detected solely by ClusTrast. Blue=detected by ClusTrast and at least one other assembler. Grey=not detected by any assembler.

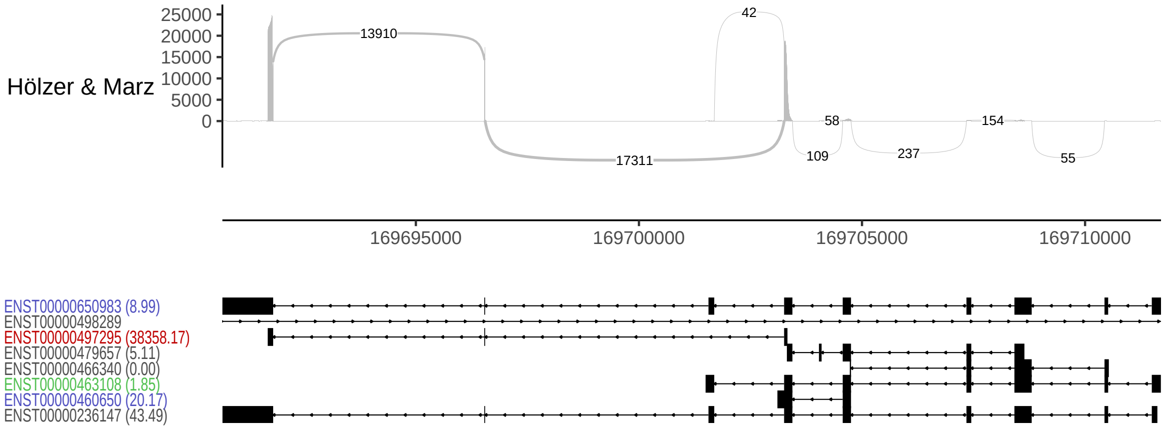


**Figure S.36:** Sashimi-plot of the simulated human dataset from Hölzer and Marz (2019) over the gene ENSG00000188404 (other isoforms shown are parts of other genes). TPM-values are shown within parethesis next to each isoform name. Red=detected solely by ClusTrast. Blue=detected by ClusTrast and at least one other assembler. Green=deteced not by ClusTrast, but some other assembler. Grey=not detected by any assembler.

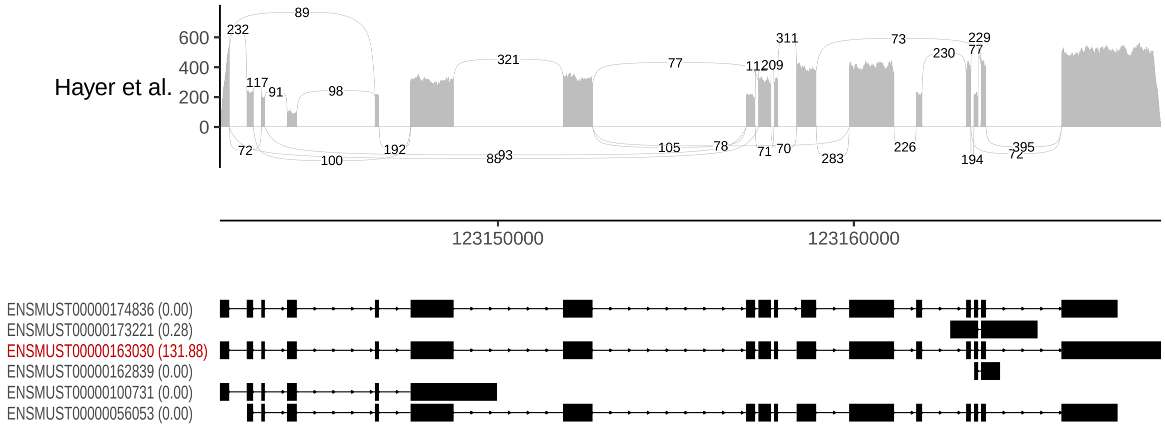


**Figure S.37:** Sashimi-plot of the simulated mouse dataset from Hayer et al. (2015) over the gene ENSMUSG00000038384. TPM-values are shown within parethesis next to each isoform name. Red=detected solely by ClusTrast.

**Table S.28:** Recall (TPR) according to SQANTI on each ClusTrast component separately: Prim.=Primary assembly (i.e. Trans-ABySS-**M**), Cl-w.=Clusterwise assembly (i.e. Shannon ran on each cluster), Merg.=The final assembly (i.e. the primary and clusterwise assemblies merged together).

| | Human | | | Arabidopsis | | | Mouse | | | Rice | | | Zebrafish | | | Poplar | | |
|---|---|---|---|---|---|---|---|---|---|---|---|---|---|---|---|---|---|---|
| | Prim. | Cl-w. | Merg. | Prim. | Cl-w. | Merg. | Prim. | Cl-w. | Merg. | Prim. | Cl-w. | Merg. | Prim. | Cl-w. | Merg. | Prim. | Cl-w. | Merg. |
| 50% | 0.3529 | 0.3937 | **0.5355** | 0.3337 | 0.2871 | **0.3923** | 0.6213 | 0.6267 | **0.7189** | 0.6069 | 0.5758 | **0.6719** | 0.5122 | 0.4827 | **0.5945** | 0.6359 | 0.6139 | **0.739** |
| 95% | 0.1831 | 0.1986 | **0.2683** | 0.2252 | 0.2036 | **0.2611** | 0.4938 | 0.4734 | **0.5641** | 0.5101 | 0.4888 | **0.5589** | 0.3277 | 0.3135 | **0.3769** | 0.4036 | 0.403 | **0.4823** |

**Table S.29:** Precision (PPV) according to SQANTI on each ClusTrast component separately: Prim.=Primary assembly (i.e. Trans-ABySS-**M**), Cl-w.=Clusterwise assembly (i.e. Shannon ran on each cluster), Merg.=The final assembly (i.e. the primary and clusterwise assemblies merged together).

| | Human | | | Arabidopsis | | | Mouse | | | Rice | | | Zebrafish | | | Poplar | | |
|---|---|---|---|---|---|---|---|---|---|---|---|---|---|---|---|---|---|---|
| | Prim. | Cl-w. | Merg. | Prim. | Cl-w. | Merg. | Prim. | Cl-w. | Merg. | Prim. | Cl-w. | Merg. | Prim. | Cl-w. | Merg. | Prim. | Cl-w. | Merg. |
| 50% | **0.1828** | 0.1452 | 0.1542 | **0.3349** | 0.3099 | 0.3205 | 0.3667 | **0.6005** | 0.5736 | **0.5737** | 0.5418 | 0.5529 | **0.1421** | 0.0843 | 0.1022 | 0.3069 | **0.4458** | 0.4031 |
| 95% | **0.0874** | 0.0684 | 0.0728 | **0.2089** | 0.1891 | 0.1973 | 0.2483 | **0.3817** | 0.3663 | **0.4133** | 0.335 | 0.3621 | **0.0741** | 0.0415 | 0.0515 | 0.1743 | **0.2456** | 0.2237 |

**Table S.30:** Recall (TPR) according to CRBB on each ClusTrast component separately: Prim.=Primary assembly (i.e. Trans-ABySS-**M**), Cl-w.=Clusterwise assembly (i.e. Shannon ran on each cluster), Merg.=The final assembly (i.e. the primary and clusterwise assemblies merged together).

| | Human | | | Arabidopsis | | | Mouse | | | Rice | | | Zebrafish | | | Poplar | | |
|---|---|---|---|---|---|---|---|---|---|---|---|---|---|---|---|---|---|---|
| | Prim. | Cl-w. | Merg. | Prim. | Cl-w. | Merg. | Prim. | Cl-w. | Merg. | Prim. | Cl-w. | Merg. | Prim. | Cl-w. | Merg. | Prim. | Cl-w. | Merg. |
| 50% | 0.2792 | 0.2870 | **0.392** | 0.2633 | 0.2107 | **0.2876** | 0.6140 | 0.5614 | **0.6648** | 0.5686 | 0.5324 | **0.6032** | 0.4723 | 0.4463 | **0.5237** | 0.5019 | 0.5174 | **0.5979** |
| 95% | 0.0826 | 0.0813 | **0.1139** | 0.103 | 0.0897 | **0.1154** | 0.3936 | 0.3175 | **0.4303** | 0.1361 | 0.1668 | **0.1906** | 0.2041 | 0.1958 | **0.2326** | 0.0968 | 0.1102 | **0.1333** |

**Table S.31:** Precision (PPV) according to CRBB on each ClusTrast component separately: Prim.=Primary assembly (i.e. Trans-ABySS-**M**), Cl-w.=Clusterwise assembly (i.e. Shannon ran on each cluster), Merg.=The final assembly (i.e. the primary and clusterwise assemblies merged together).

| | Human | | | Arabidopsis | | | Mouse | | | Rice | | | Zebrafish | | | Poplar | | |
|---|---|---|---|---|---|---|---|---|---|---|---|---|---|---|---|---|---|---|
| | Prim. | Cl-w. | Merg. | Prim. | Cl-w. | Merg. | Prim. | Cl-w. | Merg. | Prim. | Cl-w. | Merg. | Prim. | Cl-w. | Merg. | Prim. | Cl-w. | Merg. |
| 50% | **0.0454** | 0.0446 | 0.0451 | 0.1287 | **0.14** | 0.1332 | **0.0983** | 0.0482 | 0.0623 | **0.2558** | 0.2172 | 0.2357 | **0.0573** | 0.0381 | 0.0472 | **0.0977** | 0.0779 | 0.0863 |
| 95% | **0.0112** | 0.0078 | 0.0092 | 0.041 | **0.0447** | 0.0424 | **0.0462** | 0.0145 | 0.0234 | 0.0498 | **0.0539** | 0.0518 | **0.019** | 0.0117 | 0.0151 | **0.0137** | 0.009 | 0.0111 |

other isoforms from the same gene. This seems to support the idea that ClusTrast might be a useful tool to detect isoforms.
